# Live-cell quantitative monitoring reveals distinct, high-affinity Gβγ regulations of GIRK2 and GIRK1/2 channels

**DOI:** 10.1101/2025.10.15.682510

**Authors:** Reem Handklo-Jamal, Tal Keren Raifman, Boris Shalomov, Patrick Hofer, Uri Kahanovitch, Theres Friesacher, Galit Tabak, Vladimir Tsemakhovich, Haritha P. Reddy, Orna Chomsky-Hecht, Debi Ranjan Tripathy, Kerstin Zuhlke, Carmen W. Dessauer, Enno Klussmann, Yoni Haitin, Joel A. Hirsch, Anna Stary-Weinzinger, Daniel Yakubovich, Nathan Dascal

**Affiliations:** Department of Physiology and Pharmacology, Faculty of Health and Medical Sciences, Tel Aviv University, Tel Aviv, Israel; Sagol School of Neuroscience, Tel Aviv University, Tel Aviv, Israel; Department of Pharmaceutical Sciences, Division of Pharmacology and Toxicology, University of Vienna, Josef-Holaubek-Platz 2, 1090 Vienna, Austria; Department of Biochemistry & Molecular Biology, School of Neurobiology, Biochemistry and Biophysics, George S. Weiss Faculty of Life Sciences, Tel Aviv University, Tel Aviv, Israel; Max-Delbrück-Center for Molecular Medicine in the Helmholtz Association (MDC), Berlin, Germany; Department of Integrative Biology and Pharmacology, University of Texas Health Science Center, Houston, Texas, USA; DZHK (German Centre for Cardiovascular Research), partner site Berlin, Germany; The Adelson School of Medicine, Ariel University, Ariel 4077625, Israel

## Abstract

G_i/o_ protein-coupled receptors (GPCRs) inhibit cardiac and neuronal excitability via G protein-activated K^+^ channels (GIRK), assembled by combinations of GIRK1 - GIRK4 subunits. GIRKs are activated by direct binding of the Gβγ dimer of inhibitory G_i/o_ proteins. However, key aspects of this textbook signaling pathway remain debated. Recent studies suggested no G_i/o_-GIRK pre-coupling and low (>250 µM) Gβγ-GIRK interaction affinity, contradicting earlier sub-µM estimates and implying low signaling efficiency. We show that Gγ prenylation, which mediates Gβγ membrane attachment required for GIRK activation, also contributes to the Gβγ-GIRK interaction, explaining the poor affinity obtained with non-prenylated Gβγ. Using quantitative protein titration and electrophysiology in live *Xenopus* oocytes, Gβγ affinity for homotetrameric GIRK2 ranged from 4-30 µM. Heterotetrameric GIRK1/2 showed a higher Gβγ apparent affinity due to unique Gβγ-docking site (anchor) in GIRK1, which enriches Gβγ at the channel. Biochemical approaches and molecular dynamic simulations revealed that the Gβγ anchor is formed by interacting N-terminal and distal C-terminal domains of the GIRK1 subunits, distinct from the Gβγ-binding “activation” site(s) underlying channel opening. Thus, the affinity of Gβγ-GIRK interaction is within the expected physiological range, while dynamic pre-coupling of Gβγ to GIRK1-containing channels through high-affinity interactions further enhances the GPCR-G_i/o_-GIRK signaling efficiency.

## Introduction

G protein-activated inwardly rectifying K^+^ channels (GIRK; Kir3) mediate inhibitory effects of G_i/o_ protein-coupled receptors (GPCRs), controlling neuronal and cardiac excitability; GIRK malfunction is linked to neurological, cardiac and endocrine disorders^1–4^. GIRKs form homotetramers (GIRK2, GIRK4) or heterotetramers (GIRK1/2, 1/4, 1/3, 2/3), differing in tissue distribution and gating properties. Homotetrameric GIRK2 is best characterized in molecular terms, including a crystal structure in complex with Gβγ^5^. GIRKs are activated by direct, cooperative binding of up to 4 molecules of Gβγ^6–10^ (Fig. 1a). This membrane-delimited process requires posttranslational Gγ prenylation, essential for Gβγ accumulation at the plasma membrane (PM)^11^ and GIRK activation^12, 13^.

**Fig. 1.**
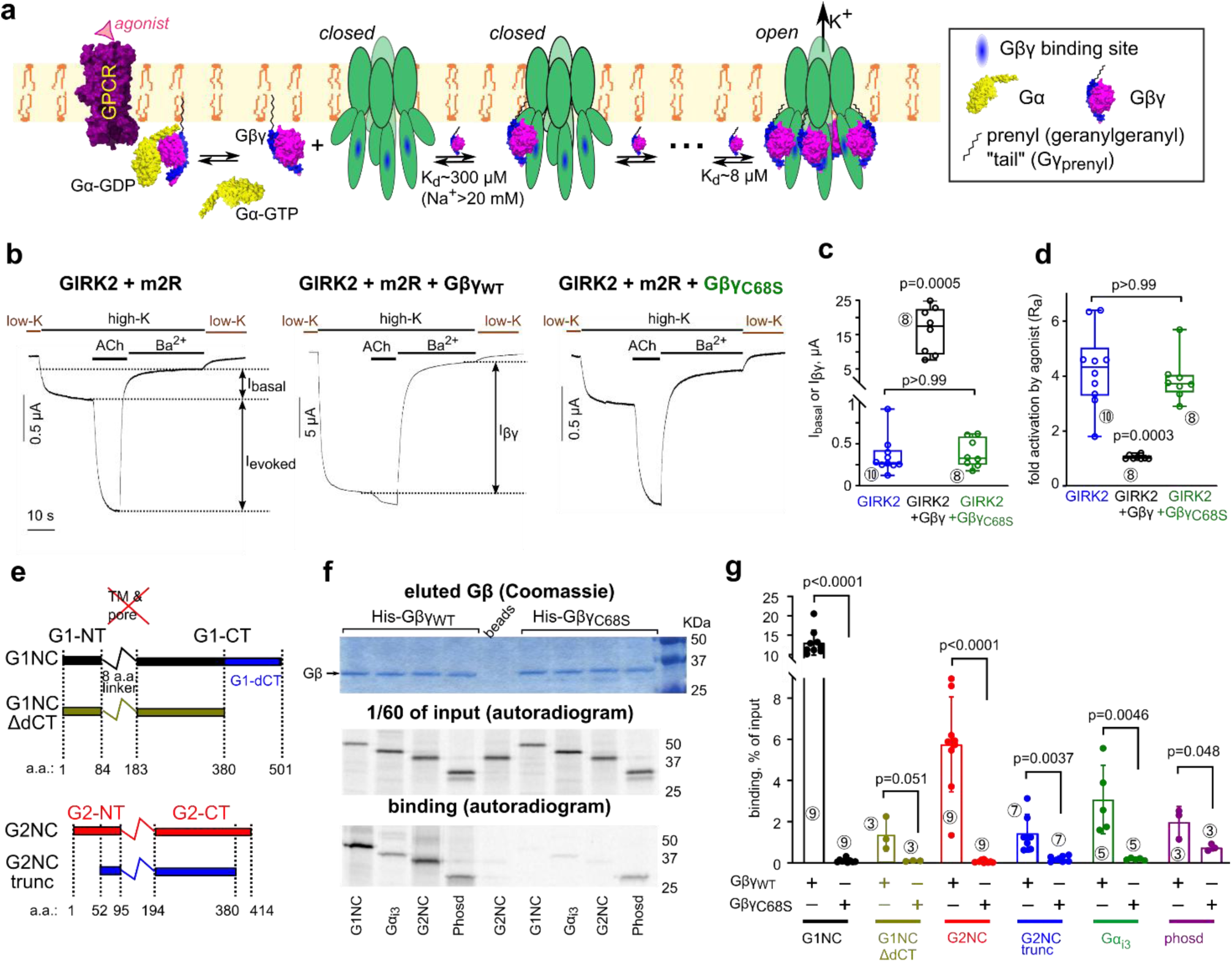
Lipid modification of Gγ is essential for GIRK activation and important for GIRK-Gβγ interaction. **a**, scheme of Gβγ activation of the GIRK2 channel. An agonist-bound GPCR (m2R) interacts with the Gα_i/o_βγ heterotrimer (Gα_i1_β_1_γ_2_, PDB: 1gp2), catalyzing the GDP-GTP exchange at Gα_i/o_ and its separation from Gβγ. Up to four Gβγ molecules bind sequentially to GIRK2. Channel opens when all four Gβγ-binding sites are occupied. The scheme shown represents the WTM model for the case of constant PIP_2_ and Na^+^ concentrations. **b**, whole-cell currents in oocytes expressing GIRK2 and m2R without Gβγ (left), with Gβγ (middle), or with Gβγ_C68S_ (right). Switching from a low-K to a high-K external solution (here 96 mM [K^+^]_out_) reveals I_basal_. ACh (10 µM) elicits I_evoked_, and then GIRK is blocked by 2.5 mM Ba^2+^, revealing the non-GIRK background current. RNA doses (ng/oocyte) were: m2R, 1; GIRK2, 2; Gβ, 5; Gγ or Gγ_C68S_, 2. **c, d**, only Gβγ, but not Gβγ_C68S_, increased I_basal_ (**c**) and abolished I_evoked_ (**d**). Statistics: Kruskal-Wallis test with Dunn’s multiple comparison vs. control (GIRK2+m2R). One experiment, representative of two. **e**, linear presentation of G1NC, G2NC and the truncated constructs. The transmembrane (TM) domains were replaced by a linker. **f,** purified prenylated His-Gβγ_WT_, captured on Ni-NTA beads, pulls down various [^35^S]Met-labeled *ivt* proteins better than the non-prenylated Gβγ_C68S_. *Top*, Coomassie staining of eluted proteins. Ni-NTA beads bound equal amounts of His-Gβγ and His-Gβγ_C68S_. *Middle*, autoradiogram of a separate gel of 1/60^th^ of the initial reaction mix (input). *Bottom*, autoradiogram of Gβγ-bound *ivt* proteins eluted from the beads (same gel as in upper image). Full gels are shown in Supplementary Fig. 2. **g**, summary of binding to Gβγ of *ivt* proteins (% of input of the same protein). Statistics for binding to His-Gβγ vs. His-Gβγ_C68S_: unpaired t-test or Mann-Whitney test (for G1NC).

GPCR-Gα_i/o_βγ-GIRK is an archetypal G protein-mediated cascade. The coupling between GPCR and Gα_i/o_βγ varies by receptor, G protein, cell type, and location, ranging from collision-coupling (e.g., muscarinic m2 receptor (m2R)^14–17^) to precoupling within dynamic multiprotein complexes (e.g., GABAB receptor with G_i/o_ and GIRK^18–20^), or a combination of both modes within protein-enriched membrane "hot spots"^21^, organized by specific scaffolds^22^ or driven by low-affinity protein interactions^23^.

Controversies linger regarding the affinity, specificity, and efficiency of Gαβγ-GIRK coupling. Early *in vitro* measurements of GIRK interaction with prenylated Gβγ yielded dissociation constants (K_d_) between 50-800 nM^8, 24^, comparable to other Gβγ interactors (typically 3 nM to 3 µM; Supplementary Table 1). Contrastingly, an NMR study reported a Kd of 250 µM for the interaction of non-prenylated Gβγ with GIRK1’s truncated cytosolic domain^25^. Wang, Touhara, MacKinnon and colleagues analyzed Gβγ activation of purified recombinant GIRK2 in-depth, while controlling the Gβγ surface density by titrating a non-prenylated His-tagged Gγ into GIRK2 and NTA lipid-containing bilayers. Their studies revealed high cooperativity of Gβγ binding and its allosteric enhancement by Na^+^ and PIP_2_^10, 15, 26, 27^. The resulting model, termed here WTM model, postulated sequential Gβγ binding to GIRK2, with channel opening when all four Gβγ sites are occupied^10, 15^ (Fig. 1a). Unexpectedly, binding of the first Gβγ showed an exceptionally low affinity, with K_d_ ∼1.9 mM at [Na^+^]=0 and ∼300 µM at saturating [Na^+^]^10^. (Due to cooperativity, the affinity increases for subsequent Gβγ bindings; Supplementary Table 2).

Low affinity entails inefficient signaling. With a K_d_>250 µM, GIRK activation (10-80%, depending on intracellular Na^+^ concentration, [Na^+^]_in_) would require free surface Gβγ exceeding 1200 µm^-2^ (molecules/µm^2^)^10^, hundredfold higher than the 2-10 µm^-2^ GIRK channel density in PM of neurons or atrial myocytes^16, 28^. While there is no evidence for such massive accumulation of Gαβγ around GIRKs, it could theoretically occur in membrane “hot spots”. Alternatively, higher affinity or dynamic (reversible) GIRK-G protein preassociation could enable fast and efficient signaling^29^. Several studies suggested preassociation of GIRKs with Gβγ or Gαβγ heterotrimers^18, 30–35^, while others support pure collision coupling^10, 16, 26^. Subunit-specific differences in GIRK-Gβγ interaction and gating may play a role. GIRK1, but not GIRK2, recruits Gβγ to the PM; the unique cytosolic distal C terminal segment of GIRK1 (G1-dCT) is essential for Gβγ recruitment^36^. We proposed that G1-dCT is part of a Gβγ-docking site (Gβγ anchor) that facilitates high-affinity, dynamic pre-association of GIRK1/2 with Gβγ^4, 17, 36–38^. However, the exact composition and interaction mode of the Gβγ-anchor remain unclear. Here we show that, besides its role in Gβγ attachment to the PM, Gγ’s prenylation directly contributes to GIRK-Gβγ interaction. We demonstrate distinct, low-micromolar interaction affinities of Gβγ with GIRK2 and GIRK1/2 in a living cell, the *Xenopus* oocyte, and determine the composition of the GIRK1’s Gβγ anchor and its role in higher apparent affinity of GIRK1/2 compared to GIRK2.

## Results

### Lipid modification of Gγ is essential for GIRK activation and important for GIRK-Gβγ interaction

All high-affinity estimates of Gβγ binding to GIRKs were obtained using prenylated Gβγ. We hypothesized that Gγ’s prenylation enhances Gβγ-GIRK interaction, as observed in Gβγ interactions with GPCRs, Gα, adenylyl cyclase and phospholipase Cβ^39–45^ (Supplementary Table 1).

In cells, the prenyl (geranylgeranyl in Gγ_2_) moiety, Gγ_prenyl_, is attached to Cys68 within the C-terminal CAAX motif, while the remaining residues are cleaved^11^. To assess the role of prenylation we used the non-prenylated mutant Gγ_C68S_ that associates with Gβ^42^; however, Gβγ_C68S_ fails to activate GIRK channels in excised PM patches^12, 13^. We expressed GIRK2 channels with m2R (adjusted to maximize I_evoked_^17^) and Gβ_1_γ_2_ (Gβγ) or Gβγ_C68S_ in *Xenopus* oocytes and measured whole-cell basal (I_basal_), agonist (acetylcholine; ACh)-evoked (I_evoked_), and Gβγ-induced (I_βγ_) GIRK currents in high-K^+^ solutions. As reported^37^, GIRK2 had a small I_basal_, which was enhanced 4-8-fold by ACh (by activating the endogenous Gα_i/o_βγ) and 30-60 fold by coexpressing nearly-saturating doses of Gβγ. In contrast, the non-prenylated Gβγ_C68S_ neither activated GIRK nor affected I_evoked_ (Fig. 1b-d, Supplementary Fig. 1a). To assess PM localization, we immunostained Gβ in oocytes’ excised giant membrane patches (GMP)^35, 46^ using wild-type (WT) Gβ or an N-terminally myristoylated Gβ (myr-Gβ). Only WT Gγ (Gγ_WT_), but not Gγ_C68S_, supported GIRK2 activation and, correspondingly, PM enrichment of WT-Gβ and myr-Gβ (Supplementary Fig. 1b-d).

These results confirm that prenylation of Gγ is essential for PM attachment of Gβγ and GIRK2 activation; but is it also involved in Gβγ interaction with GIRKs? We examined the interaction of purified, His-tagged Gβγ and Gβγ_C68S_ with *in vitro* translated (*ivt*) Gβγ-binding proteins: Gα_i3_; phosducin; cytosolic domains of GIRK1 and GIRK2 (G1NC and G2NC, respectively); and their truncated versions, G1NC_ΔdCT_ and G2NC_trunc_ (Fig. 1e). G1NC is a fusion protein of N- and C-terminal domains of GIRK1 (G1-NT and G1-CT). G1NC_ΔdCT_ lacks the G1-dCT and binds Gβγ much weaker than G1NC^36^ (Supplementary Fig. 2). G2NC_trunc_ lacks the distal segments of the N- and C-terminal domains (G2-NT and G2-CT, respectively), as in structural and bilayer studies^5, 26, 47, 48^. All *ivt* proteins bound Gβγ. Remarkably, lack of prenylation dramatically reduced Gβγ interaction with Gα_i3_ and phosducin, corroborating previous reports^39–41^, and with all GIRK constructs (Fig. 1f,g), suggesting that Gγ prenylation directly contributes to Gβγ-GIRK interaction.

### Estimating Gβγ density in PM using calibrated fluorescence and quantitative Western blotting

We aspired to quantitatively analyze the membrane-delimited GIRK-Gβγ interaction in intact cells, using prenylated Gγ. A major challenge was to accurately calibrate protein surface density. To this end, we extended our previously developed calibration methods in *Xenopus* oocytes^17, 38^, which use two independent approaches.

The calibrated fluorescence (CF) approach measures the surface density of yellow, cyan or Split-Venus fluorescent proteins (YFP, CFP or SpV; collectively xFP), using xFP-labeled channels as molecular calipers. We used Gβγ-activated xFP-GIRK1/2^38^, and additionally the constitutively active homotetrameric IRK1-xFP (usually IRK1-YFP; Fig. 2a,b). Calibration involved expressing these channels at varying RNA doses, measuring whole-cell currents, and calculating the surface density of functional channels based on open probability (P_o_), single-channel current (i_single_) and cell’s surface area^49^ (Eqn. 1 in Methods, Supplementary Fig. 3, Supplementary Table 3). YFP surface density was calculated assuming two or four YFP molecules per YFP-GIRK1/2 or IRK1-YFP channel, respectively. To avoid artifacts arising from any non-functional channels, we used channels’ RNA doses in the 0.01-1 ng range, ensuring a linear relationship between fluorescence and whole-cell current and, accordingly, the calculated YFP surface density (Fig. 2a). Deviations were observed only at high levels of YFP-GIRK1/2 (5 ng RNA; Supplementary Fig. 6a). Additionally, we compared the calibration with both YFP-GIRK1/2 and IRK1-YFP in the same experiment (Fig. 2a,b). The relationship between fluorescence and calculated YFP surface density was almost identical with both calipers.

**Fig. 2.**
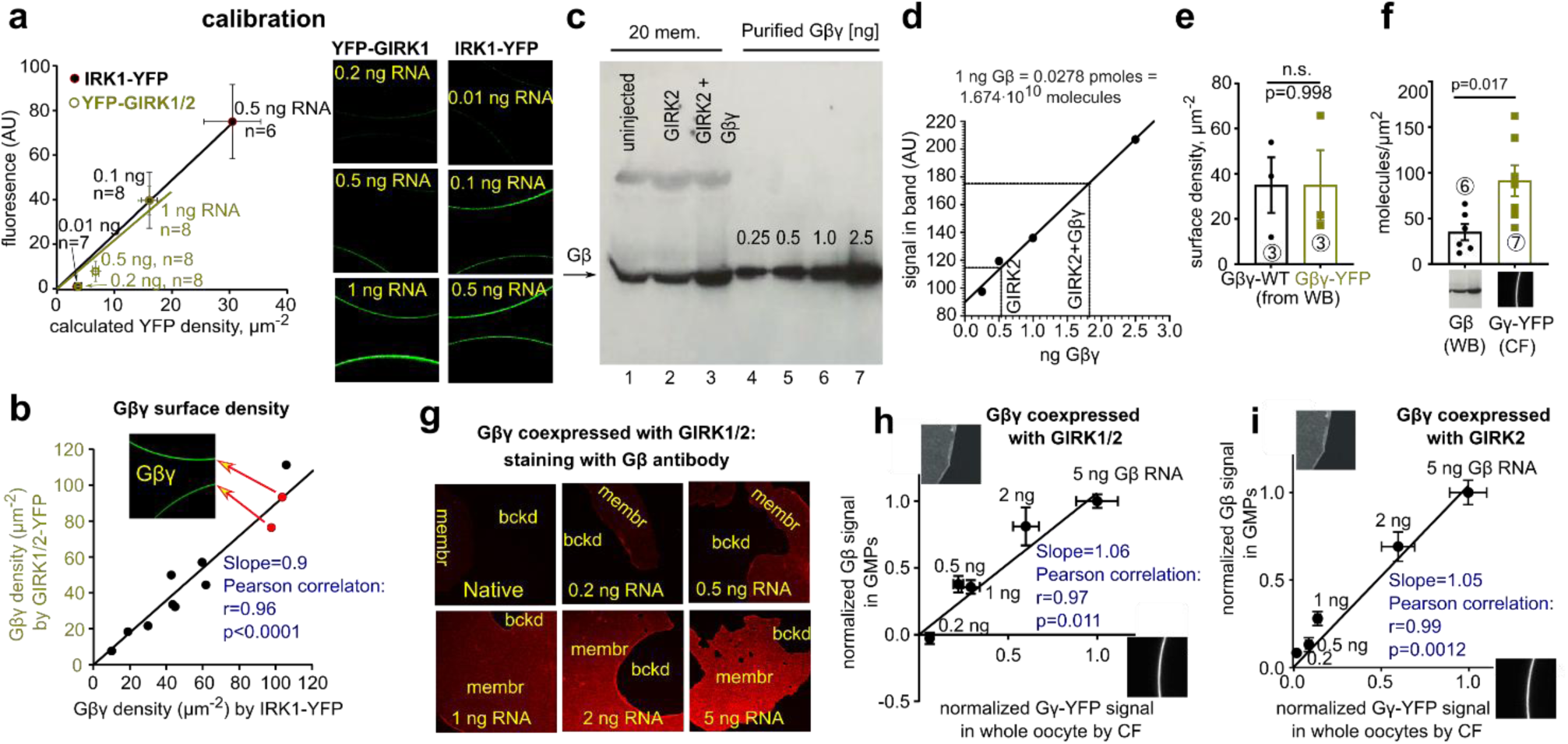
Estimating Gβγ density in PM using calibrated fluorescence (CF) and quantitative Western blotting (qWB). In oocyte experiments RNAs of YFP-Gγ and Gβ were injected at a constant ratio. **a,** calibrating surface YFP-Gγ density with YFP-GIRK1/2 coexpressed with Gβγ (5:2 ng RNA/oocyte) or IRK1-YFP. Amounts of channel RNA are shown near symbols. Surface density of channel-associated YFP was estimated from whole-cell currents. YFP fluorescence (in arbitrary units, AU) was measured from confocal images of intact oocytes (right panel). **b,** calibration with either IRK1-YFP or YFP-GIRK1/2 gives similar estimates of surface density of Gβ·_YFP_Gγ (same experiment in **a**). Data points represent individual oocytes. Inset shows representative oocytes (red symbols). **c,** measuring PM-attached Gβ (20 plasma membranes per lane) using WB with a Gβ antibody that well recognizes both endogenous and expressed Gβ^38^, from naïve (uninjected) oocytes, or injected with GIRK2 RNA (2 ng) without or with Gβγ (5:2 ng RNA/oocyte). Lanes 4-7: calibration with recombinant Gβγ (0.25-2.5 ng/lane). **d,** estimating the amounts of Gβγ in PMs for lanes 1-3 from the calibration plot drawn using linear regression of data from lanes 4-7. **e,** qWB-estimated surface density of Gβ, coexpressed with either Gγ or YFP-Gγ, is similar. Net amounts of Gβ were calculated in each experiment by subtracting the Gβ level of GIRK2-only expressing oocytes. Statistics: unpaired t-test. **f**, comparing the estimated levels surface density of YFP-Gγ (by the CF approach) and Gβ (by the qWB approach. Data with Gγ and YFP-Gγ were pooled). Statistics: unpaired t-test. **g**, representative confocal images of GMPs from oocytes expressing Gβ, YFP-Gγ, and GIRK1/2 or GIRK2. Amounts of Gβ RNA are shown. **h,i**, Gβ levels in GMPs and YFP-Gγ levels in intact oocytes are linearly correlated. Protein levels induced by different RNA doses were normalized to 5 ng Gβ in each experiment. Numbers of experiments and cells are shown in Supplementary Table 5.

Concomitantly, we expressed Gβ·_YFP_Gγ (Gβ and N-terminally labeled YFP-Gγ) in separate groups of oocytes, measured YFP fluorescence at the oocyte’s perimeter, and converted it to YFP-Gγ surface density with each caliper. The estimates of YFP-Gγ with both calipers showed strong linear correlation with a slope of 0.9 (Fig. 2b), validating the calibration protocol.

The CF procedure with Gβ·_YFP_Gγ monitors YFP-Gγ rather than Gβ. We directly assessed the surface density of Gβ using the independent approach^38^, quantitative Western blotting (qWB) of manually separated oocyte plasma membranes. We measured PM-associated Gβ with a Gβ antibody, using purified recombinant Gβγ for calibration (Fig. 2c,d). The PM density of the endogenous oocyte Gβ was 30±13 µm^-2^, consistent with previous estimates (Supplementary Table 4) and comparable to ∼40 µm^-2^ in HEK cells^50^. Expressed Gβ surface levels were similar with either coexpressed Gγ or YFP-Gγ (Fig. 2e, Supplementary Table 4). Overall, expressed surface Gβ (with 5 ng Gβ RNA) measured by qWB was 35±9 µm^-2^ (n=6), about 2.5-fold lower than surface YFP-Gγ estimated by CF (91±19 µm^-2^, n=7, Fig. 2f). The difference is probably not related to methodology, because previously both CF and qWB gave similar estimates of 22-28 µm^-2^ for a YFP-labeled Gβ^38^. Thus, evaluating YFP-Gγ may overestimate the coexpressed Gβ’s surface density, possibly because YFP-Gγ associates with endogenous Gβ, or exists as a separate protein^51, 52^. Therefore, we tested a variety of C- or N-terminally xFP-fused Gβ constructs (Supplementary Fig. 4). However, they yielded partial or no GIRK2 activation, and usually poorly activated GIRK1/2. SpV-Gβγ activated both GIRK1/2 and GIRK2 but induced smaller currents than WT-Gβγ. Only Gβ·_YFP_Gγ activated GIRK channels like the WT-Gβγ^36^.

We next varied expression levels of Gβ·_YFP_Gγ and examined changes in surface densities of YFP-Gγ in intact oocytes and Gβ in GMPs (Fig. 2g). Reassuringly, there was a linear correlation between surface levels of Gβ and YFP-Gγ with either GIRK2 or GIRK1/2 channels coexpressed (Fig. 2h,i). Thus, RNA dose-dependent changes in surface YFP-Gγ reflect corresponding changes in surface Gβ. Consequently, we routinely used Gβ·_YFP_Gγ in the following experiments.

### Affinity of Gβγ-GIRK2 interaction is in the low µM range

We investigated the dose-dependent activation of GIRK2 by Gβ·_YFP_Gγ using the CF approach. We expressed GIRK2 at a low density, with a range of Gβ·_YFP_Gγ RNA doses. Following calibration (Fig. 3a), we quantified surface Gβ·_YFP_Gγ density in oocytes expressing GIRK2 and Gβ·_YFP_Gγ and then measured single-channel P_o_ in cell-attached patches of the same oocytes (Fig. 3b-d). The activation of GIRK2 was steeply Gβ·_YFP_Gγ dose-dependent, with an initial slope of almost 3 on log-log coordinates (Fig. 3e). This indicates the requirement for at least 3 Gβγ molecules to open the channel, corroborating the WTM model^10^ (Figs. 1a, 3e). Therefore, we analyzed the dose-response data using the WTM model version adjusted for real-cell conditions^15^ (Fig. 3e, Supplementary Fig. 8a #2, Methods Eqn. 5) and, for comparison, the familiar but mechanistically less informative Hill equation (Eqn. 4). We added to the equations a constant component (c) corresponding to I_basal_. To convert the two-dimensional surface density to concentration we used a standard procedure^10, 15, 38, 53^ assuming a submembrane 10 nm thick interaction volume.

**Fig. 3.**
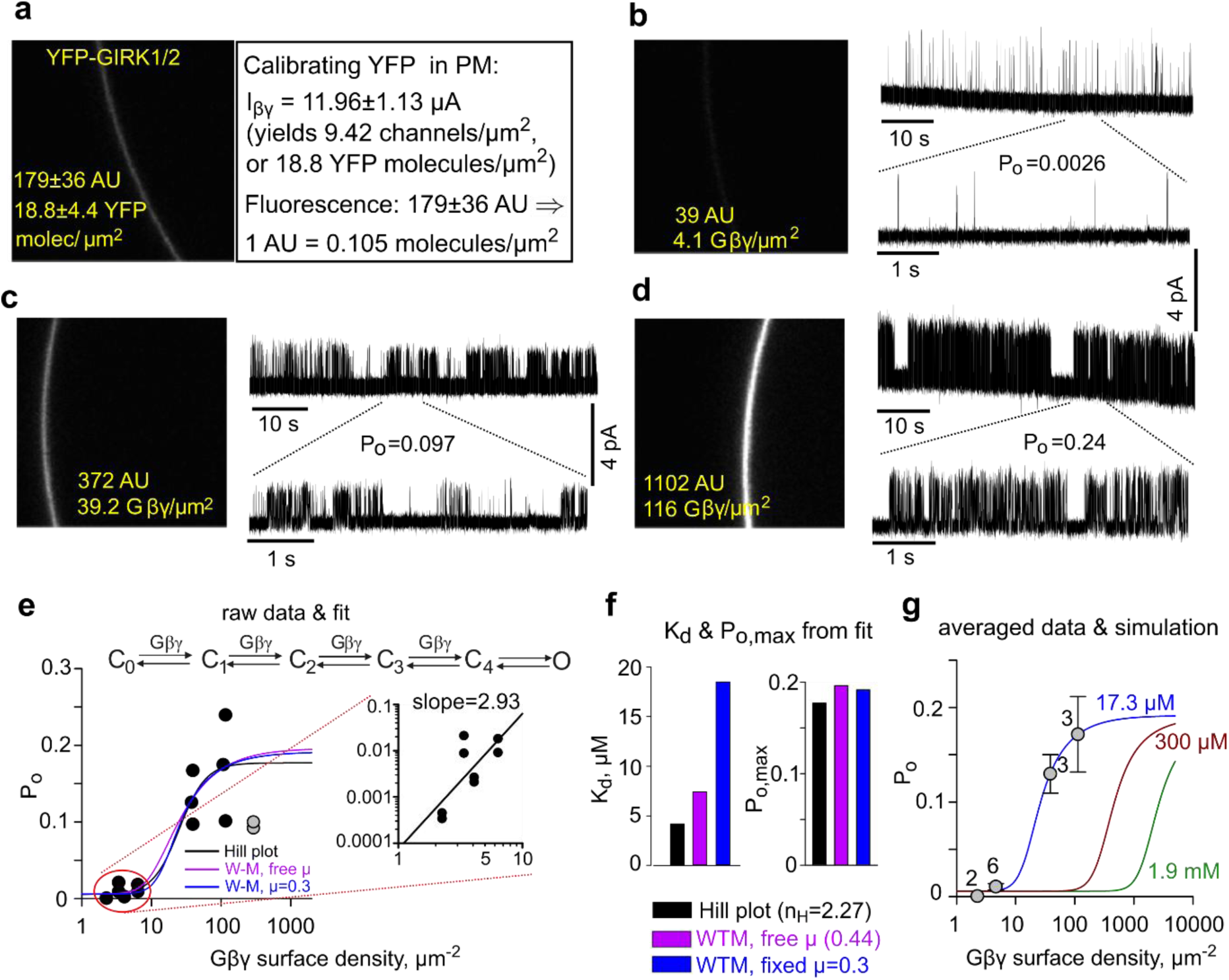
Coexpressed Gβ·_YFP_Gγ activates single GIRK2 channels with low-µM apparent affinity. P_o_ and Gβ·_YFP_Gγ expression were measured in the same oocytes, injected with RNA of GIRK2 (25 or 50 pg/oocyte), Gβ (0.2-20 ng/oocyte) and YFP-Gγ (40% of Gβ RNA). **a**, calibration of surface density of YFP using YFP-GIRK1/GIRK2 (1 ng RNA each) coexpressed with WT-Gβγ (5:2 ng RNA, respectively). **b-d**, representative confocal images of intact oocytes, and cell-attached patch records from these oocytes. **e**, changes in P_o_ vs. estimated Gβ·_YFP_Gγ PM density. Each circle represents P_o_ measurement in a separate patch. Low P_o_ observed in two patches from one oocyte (grey circles) with high surface Gβ·_YFP_γ (290 µm^-2^) was attributed to Gβγ-induced desensitization, as reported previously for high [Gβγ] for GIRK1/4 and GIRK1/2^9, 46^. These patches were excluded from fit. Lines show fits to Hill equation and to the WTM model, the latter with either fixed (µ=0.3) or free cooperativity factor µ. *Inset* (right) shows the log(P_o_)-log[Gβ·_YFP_Gγ] plot for the lowest Gβ·_YFP_Gγ expression levels. The slope of the linear regression (black line) was 2.93. Hill coefficient (n_H_) in the Hill plot fit was 2.37. The average Gβ·_YFP_Gγ density at 5 ng Gβ RNA was 39.7±6 μm^-2^ (n=12 oocytes). **f**, K_d_ and P_o,max_ values from fits shown in e. For a full set of WTM fit parameters, see Supplementary Table 6. **g**, simulated Gβγ dose-response curves with µ=0.3 and c=0.03, P_o,max_=0.19, K_d_=17.3 µM from the WTM fit of our data shown in **f**, compared to values reported by Wang et al.^10^: K_d_=1.9 mM for [Na^+^]_in_=0 and K_d_=300 µM for high [Na^+^]_in_ (>20 mM). For visualization purposes, P_o_ values from patches with similar Gβ·_YFP_Gγ levels were pulled and presented as mean±SEM.

Fitting the data with the WTM model (Fig. 3e) yielded cooperativity factor for each successive Gβγ binding (µ) of 0.44 and dissociation constant (K_d_) of 44 Gβγ µm^-2^ (7.4 µM). Fixing µ=0.3 as in Touhara et al.^15^, yielded a K_d_ of 17.3 µM, and Hill equation fit yielded a K_d_ of ∼4 µM (Fig. 3f). This is much lower than the 300 µM measured in bilayers even at saturating [Na^+^] of >20 mM^10^, as highlighted with simulated dose-response curves in Fig. 3g.

Similar K_d_ values were obtained for whole-cell currents of GIRK2 or HA-tagged GIRK2_HA_ (which is activated by Gβγ like GIRK2^36, 37^, Supplementary Fig. 1a). Fitting with WTM model (with fixed µ, to reduce the number of free parameters) yielded K_d_ of ∼11 µM with µ=0.44 and ∼31 µM with µ=0.3 (Fig. 4f; Supplementary Table 6). In one experiment we compared WT-GIRK2 and truncated GIRK2 (as used in lipid bilayers); they showed similar Gβγ sensitivity (Supplementary Fig. 5).

**Fig. 4.**
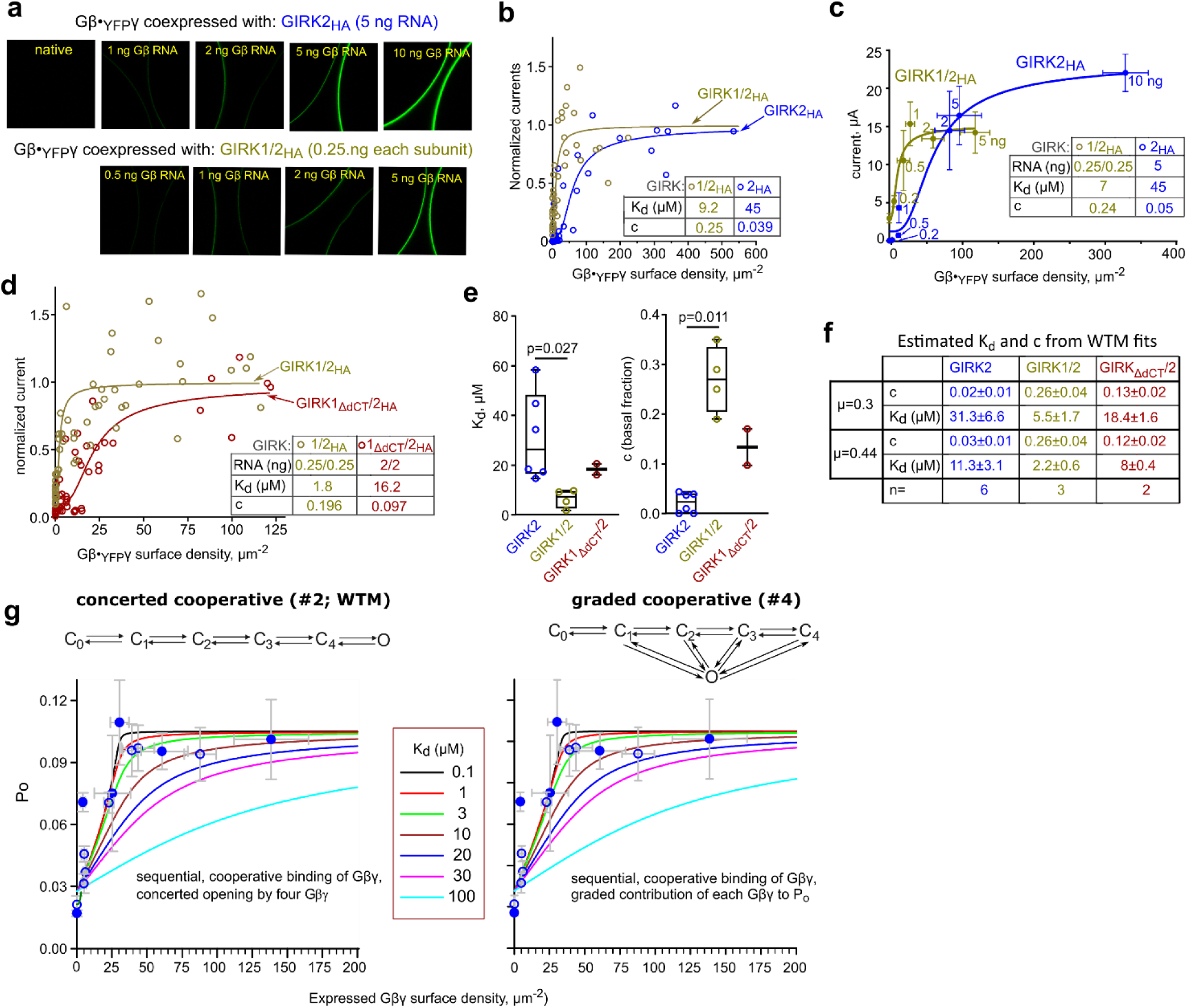
GIRK2 and dCT-truncated GIRK1 show lower apparent affinity to Gβγ than GIRK1/2. **a-d,** GIRK2_HA_ was used in these experiments. Gβ:YFP-Gγ RNA ratio was 2:1. RNA doses of GIRKs and WTM fit parameters are shown in insets in b-d. Surface density of YFP was calibrated using IRK1-YFP. Currents were measured in 24 mM [K^+^]_out_. **a-c,** dose-dependent activation of GIRK2_HA_ homotetramers and GIRK1/2_HA_ heterotetramers by Gβ·_YFP_Gγ (experiment #4). **a**, examples of confocal images in oocytes expressing Gβ·_YFP_Gγ with GIRK1/2_HA_ or GIRK2_HA_. **b,** dose-dependent activation of GIRK1/2_HA_ and GIRK2_HA_ by Gβ·_YFP_Gγ. Each point represents an individual oocyte. Currents were normalized to the maximal I_βγ_ (I_max_, Supplementary Table 6) and fitted to the WTM model (with µ = 0.3). The differences between the fitted K_d_ were significant (F(1, 81)= 18.95, p<0.0001). See additional analysis in Supplementary Fig. 7a. **c**, results of the same experiment were analyzed for groups of oocytes according to the amount of Gβ RNA (shown near each point). Data are presented as mean ± SEM of I_βγ_ and YFP-Gγ; numbers of oocytes are shown in Supplementary Table 5. **d**, dose-dependent activation of GIRK1/2_HA_ and GIRK1_ΔdCT_/2_HA_ by Gβ·_YFP_Gγ. (Experiment #7; additional details in Supplementary Fig. 7b). Analysis and presentation of data are as in b. The differences between fitted K_d_ were significant: F(1, 103)=14.18, P=0.0003). **e, f**, summary of parameters of the WTM fit with fixed µ=0.3 for all experiments (**e**; statistics: Kruskal-Wallis test) and with µ=0.3 or µ=0.44, presented as mean±SEM (**f**). See Supplementary Table 6 for full details. **g**, simulation of GIRK1/2_HA_ activation by Gβγ with a range of K_d_ values (solid lines) with the cooperative models (Supplementary Fig. 8a). The simulated curves are superimposed on data from experiments #4 (closed circles) and #7 (open circles). More details in Supplementary Fig. 8c.

### GIRK1/2 vs. GIRK2: higher apparent affinity to Gβγ and the role of Gβγ docking to GIRK1

Heterologously expressed GIRK1/2 has a high, Gβγ-dependent I_basal_, contrasting the smaller, Gβγ-independent I_basal_ of homotetrameric GIRK2^35, 37, 46, 54, 55^. Gβγ recruitment^36^ and high I_basal_ of GIRK1/2 and GIRK1/4 require an intact G1-dCT^37, 55, 56^, suggesting that Gβγ docking by GIRK1 increases the local concentration of Gβγ around GIRK^4, 17, 36, 38^. We hypothesized that this may also render higher apparent Gβγ affinity for GIRK1/2 compared to GIRK2.

We previously observed GIRK1/2 activation by expressing Gβγ at relatively low densities (5-50 µm^-2^)^38^. Here, we compared activation of GIRK2 and GIRK1/2 by Gβ·_YFP_Gγ in the same experiment (Fig. 4a-c). Surface levels of YFP-Gγ and, subsequently, GIRK currents were measured in individual intact oocytes. Fitting these data with the WTM model revealed a significant difference between K_d_ of GIRK2 and GIRK1/2 (45 and 9 µM, respectively, with µ=0.3, p=<0.0001; Fig. 4b, Supplementary Fig. 7a). Similar K_d_ values were obtained for data grouped according to the RNA dosage (Fig. 4c). On average, the K_d_ of GIRK1/2 was about 6-fold lower than GIRK2 (∼5.5 µM vs. ∼31 µM, p=0.027, Fig. 4e,f, Supplementary Table 6). We also observed an ∼8-fold difference in K_d_ of GIRK2-CFP and GIRK1/2-CFP (Supplementary Fig. 6). GIRK1/2 also exhibited the expected higher I_basal_ than GIRK2. The basal fraction (c) was ∼0.26 in GIRK1/2 and 0.02-0.03 in GIRK2 (Fig. 4e,f).

To investigate the role of Gβγ-anchor, we compared the Gβγ dose-dependence of GIRK1/2 to GIRK1_ΔdCT_/2. GIRK1_ΔdCT_/2 lacks the Gβγ-anchor, does not recruit Gβγ and has a reduced I_basal_^36^. Remarkably, the K_d_ of GIRK1_ΔdCT_/2 was 9-fold higher compared to GIRK1/2 (Fig. 4d; p=0.0003) and 3.8-fold higher in another experiment (Supplementary Fig. 7c; p=0.0009). Thus, GIRK1’s Gβγ-anchor contributes to the high apparent Gβγ affinity of GIRK1/2.

We added the c parameter to the original WTM model to account for I_basal_. Instead of fitting c, I_basal_ can be mechanistically explained and calculated using algorithms utilizing I_basal_, I_evoked_ and I_βγ_ to estimate basal Gβγ and Gα in GIRK1/2 microenvironment^17, 38^. We compared the modified WTM (concerted cooperative), the graded contribution (channel opens with one Gβγ and sequential Gβγ binding progressively increases P_o_^9, 57^), and two non-cooperative models (Fig. 4g, Supplementary Methods, Supplementary Fig. 8). With each model, we calculated basal Gα, Gβγ and I_basal_ for a range of K_d_ values, and subsequently simulated dose-response curves for expressed Gβγ with µ=0.3. Both cooperative models matched the experimental data with K_d_ between 1-10 µM (Fig. 4g). Expectedly, the non-cooperative models predicted lower K_d_. The cooperative models also provided stable estimates of basal Gα and Gβγ across a wide K_d_ range, 0.1-30 µM (Supplementary Fig. 8).

The interactions of Gβγ with the PM and the channel are reversible. Therefore, we expected that removing the cytosolic Gβγ reserve by excising a membrane patch into a Gβγ-free solution would reduce the PM- and GIRK-associated Gβγ, deactivating GIRK channels. We anticipated slower deactivation in channels with a high-affinity Gβγ-anchor.

To test this hypothesis, we recorded Gβγ-activated channels in cell-attached patches and then excised them into an ATP and Na^+^-containing bath solution (Fig. 5a-d). GIRK1/2 activity decayed (deactivated) slowly, with 30-50% persisting after 5 minutes (Fig. 5a,d). The decay followed a single exponent with a time constant (τ) of >2 min and a non-deactivating fraction (C) of 0.34. In contrast, GIRK2 and GIRK1_ΔdCT_/2 exhibited faster and more complete decay (Fig. 5b-d,f). Excising patches into an ATP-free solution, which could deplete PIP_2_ in the PM^58^, had a minimal impact on GIRK2 and GIRK1_ΔdCT_/2 decay, and slightly affected GIRK1/2 (Fig. 5e,f). This suggests that GIRK deactivation is mainly governed by the depletion of Gβγ associated with or surrounding the channel, rather than PIP_2_ depletion.

**Fig. 5.**
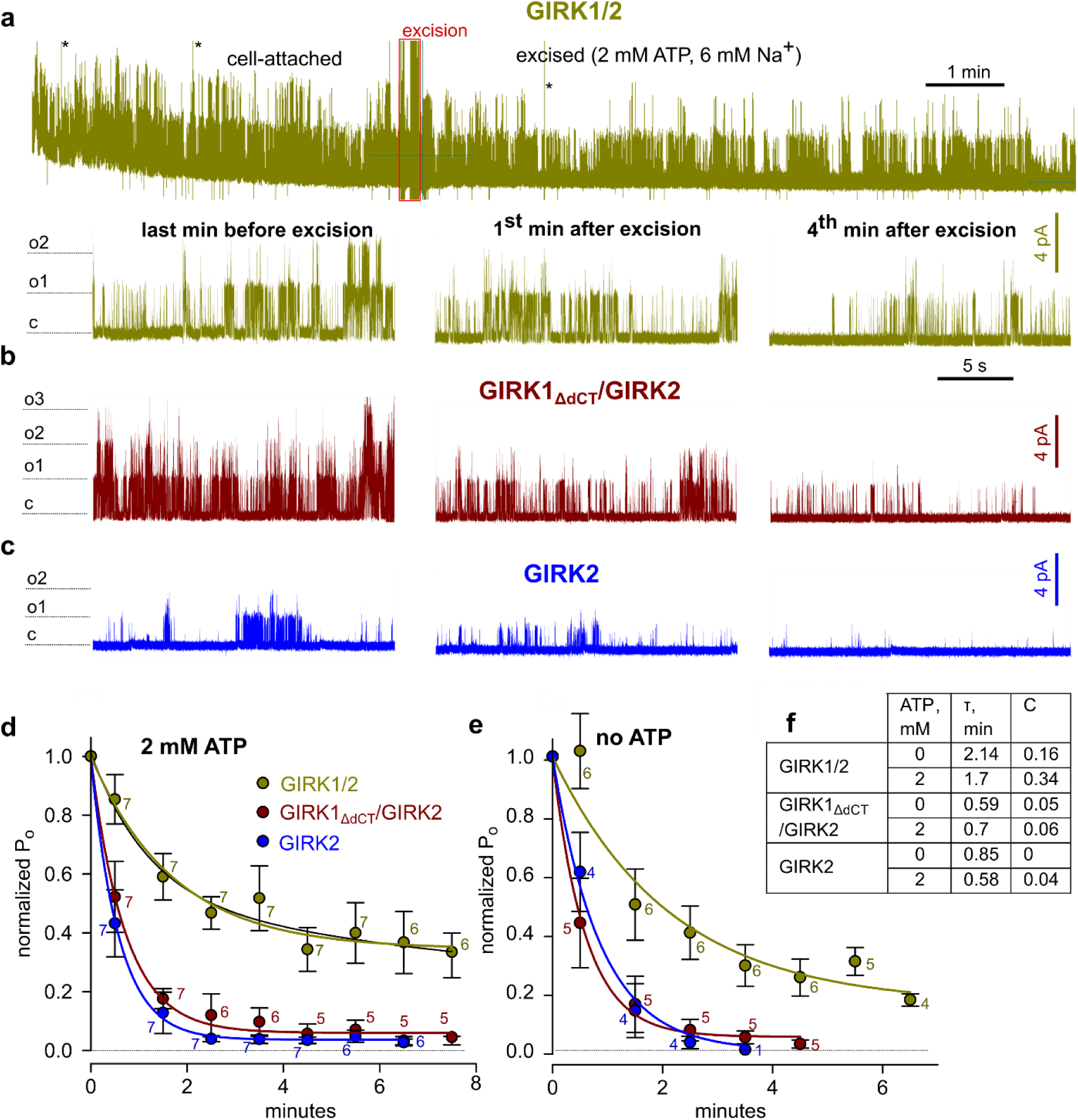
Different patterns of deactivation of GIRK2 and GIRK1/2 after patch excision and the role of G1-dCT. Channels were expressed at low densities, with a high dose of Gβγ or SpV-Gβγ (5 ng Gβ and 1 ng Gγ). **a**, representative recording of GIRK1/2. *Top*, the complete original recording that lasted 13.5 min. After ∼4 min in cell-attached mode, the patch was excised into bath solution containing 2 mM ATP and 6 mM NaCl, causing a gradual decay of activity. *Bottom*, zoom on 20 s segments of the record during the indicated times before and after excision. **b**, **c**, similar stretches from recordings of representative GIRK1_ΔdCT_/2 and GIRK2 recordings. **d**, time course of deactivation after excision summarized as NP_o_ within consecutive 60 s segments of record, normalized to NP_o_ during the last minute before excision. (NP_o_ is a measure of total activity in the patch, i.e. number of channels times P_o_). Each point is mean±SEM, with number of patches shown near each symbol. Lines show single-exponential fits; fitting with two exponents did not produce better results (exemplified for GIRK1/2 with ATP, black line). **e**, similar results were obtained when the patches were excised into an ATP-free solution. **f**, comparison of exponential fit parameters for the three channel types, with and without ATP. τ is the time constant of the exponential decay and C is the extrapolated non-deactivating fraction.

### G1-NT and G1-dCT form a Gβγ-binding site and contribute to channel’s interaction with Gγ’s prenylation tail, Gγ_prenyl_

Although deleting G1-dCT thwarts Gβγ binding, G1-dCT alone does not strongly bind Gβγ^59^, indicating that the Gβγ-anchor includes additional Gβγ-binding segments^36^. To identify these regions, we scanned arrays of overlapping peptides covering the cytosolic domains of GIRK1 and GIRK2 (G1NC, G2NC) for His-Gβγ binding (Fig. 1e, 6a-c, Supplementary Fig. 9). Scanning revealed three Gβγ-binding segments mainly overlapping the C1 and C3 segments from previous biochemical studies^30, 59^. Two segments fully (in GIRK2) or partially (in GIRK1) overlapped the Gβγ-binding amino acid (a.a.) clusters from the crystallized GIRK2/Gβγ complex^5^ (Fig. 6d). Additionally, Gβγ bound segments in G1-NT (a.a. ∼20-50), parts of G1-dCT (a.a. ∼390-440 and ∼485-501), and G2-NT and G2-dCT.

**Fig. 6.**
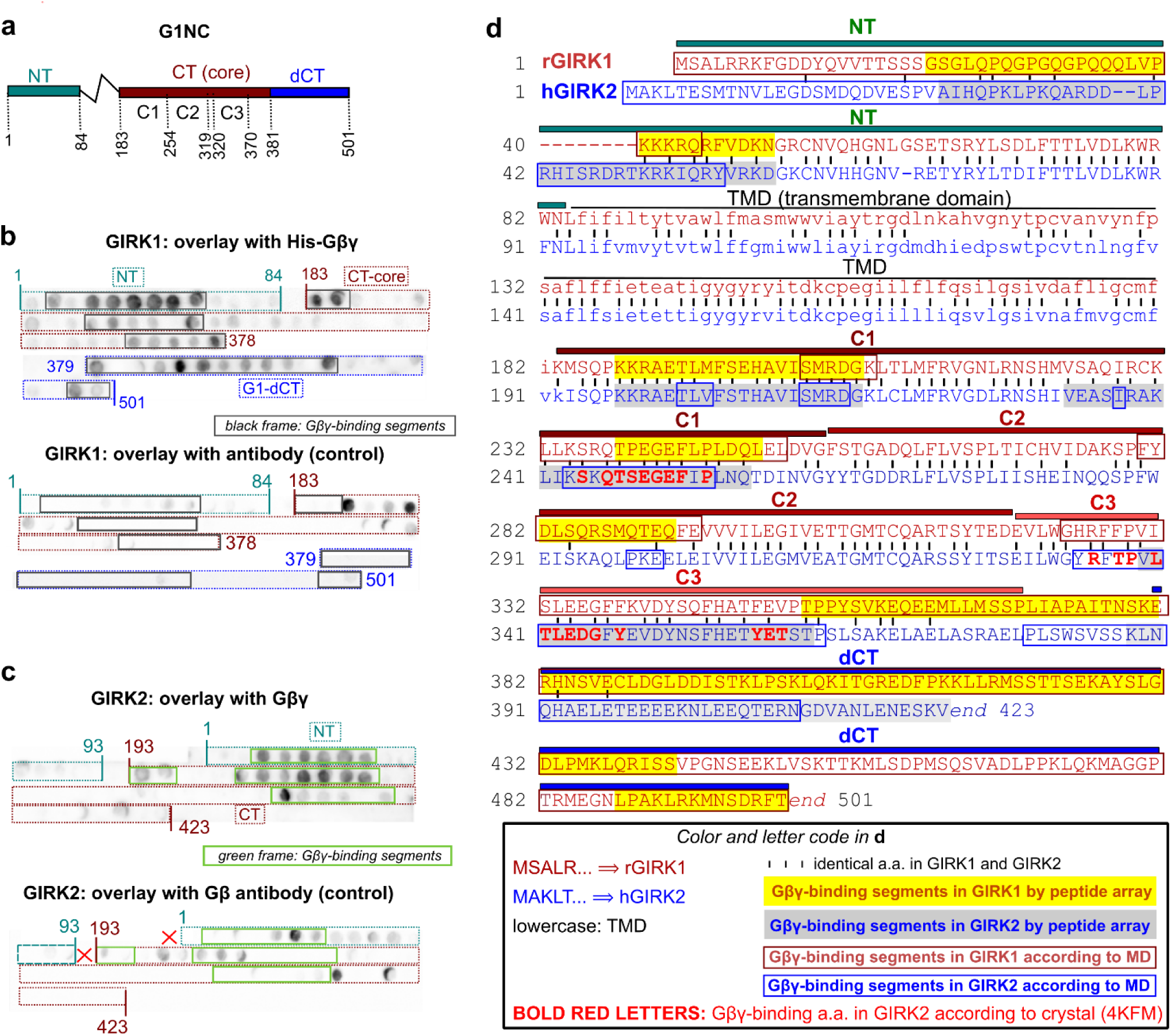
Peptide array scanning for Gβγ binding sites in the cytosolic domains of GIRKs. **a,** linear scheme of G1NC incorporating segment names (NT, CT, etc.) and a.a. numbers illustrating the design of the peptide array (b) and the constructs used in pull down experiments of Fig. 7. **b**, **c,** arrays of 25-mer overlapping peptides with a 5 a.a. shift of G1NC (b) and G2NC (c), spotted onto a membrane. Upper images show overlays with purified His-Gβγ, probed with the Gβ antibody (4 experiments for G1NC, 3 for G2NC). Gβγ-binding segments are enclosed within solid-border rectangles. Bottom images show control arrays overlayed with Gβ antibody only (two experiments for each channel). In GIRK2 some non-specific labeling (without Gβγ) was observed in segments designated as Gβγ-binding. The non-specific labeling was weaker and appeared in fewer spots, therefore we have not discarded these spots from the area assigned as Gβγ-binding. **d,** alignment of rGIRK1 (rat GIRK1) and hGIRK2 (human GIRK2) a.a. sequences used in peptide array scans. The Gβγ-binding segments suggested by peptide arrays are highlighted in yellow (GIRK1) and gray (GIRK2). A weakly labeled potential Gβγ-binding segment in the distal CT of hGIRK2 is labeled with a lighter gray background. Gβγ-binding segments suggested by molecular dynamics (MD) simulations (from Fig. 8) are framed by dark red (GIRK1) and blue (GIRK2) rectangles. Amino acids in GIRK2 that make contacts with Gβγ according to the crystal structure of the GIRK2-Gβγ complex, 4KFM^5^, were determined using the Prodigy software (https://rascar.science.uu.nl/prodigy/) and are highlighted in bold red letters.

If any of the GIRK1’s Gβγ-binding segments combines with G1-dCT to form the Gβγ-anchor, deleting it from G1NC should reduce Gβγ binding. We used prenylated His-Gβγ to pull-down the full-length *ivt* G1NC or G1NC with specific segment deletions, and a fusion protein of G1-NT and G1-dCT, G1NdCT (Fig. 7). Gβγ binding was unaffected by the deletion of internal segments C1-C3 and tended to decrease after the deletion of G1-NT (G1CT construct). G1-dCT and G1-NT showed weak and negligible Gβγ binding, respectively. However, both G1NdCT and the fusion of the second half of G1-NT (a.a. 40-84) with G1-dCT bound Gβγ very strongly, suggesting that the GIRK1’s Gβγ-anchor comprises G1-dCT and part(s) of G1-NT.

**Fig. 7.**
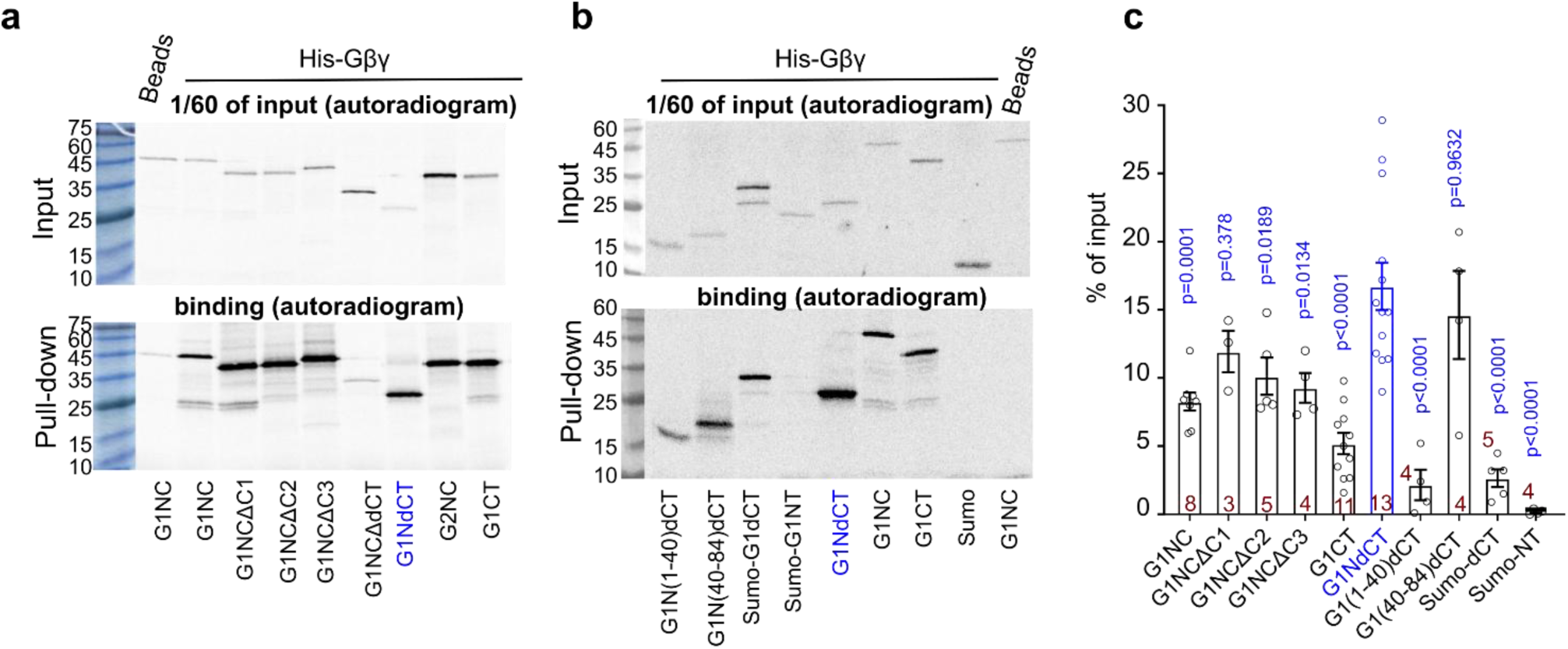
Fused G1-NT and G1-dCT of GIRK1 form a high-affinity Gβγ-binding site. **a, b,** SDS-PAGE autoradiograms of pull-down of [^35^Met]-labeled *ivt* G1NC, G1NC-derived constructs and additional controls by His-Gβγ_WT_ from two representative experiments. G1-NT and G1-dCT were fused to Sumo for stability. **c**, summary of pull-down experiments. Binding of each construct was calculated as percentage of input of that construct in the same experiment. Statistics: One Way ANOVA followed by Dunnet’s multiple comparison method vs. control group, G1NdCT. Statistics for G1NC comparisons are shown in Supplementary Table 7.

We conducted coarse-grain molecular dynamics (MD) simulations to further investigate the involvement of G1-NT, G1-dCT and Gγ_prenyl_ in GIRK-Gβγ interactions. These elements are missing from the available high-resolution structures. Creating a system where Gβγ is added *ab initio* and equilibrates with the channel and PM is challenging. Therefore, the initial system included four Gβγ molecules bound to a G1NC or G2NC tetramer without the PM and bulk Gβγ in the cytosol.

We modeled full-length and truncated G1NC and G2NC tetramers complexed with Gβγ using AlphaFold3 and manually added the prenylation tails (Fig. 8, Supplementary Fig. 10a). MD simulations accurately captured the two Gβγ-interacting surfaces from the GIRK2-Gβγ crystal structure^5^ and predicted additional Gβγ-binding segments. Most of these segments showed excellent (in G2NC) or considerable (in G1NC) agreement with peptide arrays (Fig. 6d, 8b, Supplementary Fig. 10b), lending credibility to the combined analysis. Further analysis revealed that Gγ_prenyl_ spent 100% of the simulation time interacting with G1NC, mainly with the beginning of G1-NT, as compared to only 6.4% with G2NC (Fig. 8c,d). This interaction likely accounts for most of the Gβγ binding to the first NT segment predicted by the MD (compare Fig. 8b and 8c), explaining the poor Gβγ labeling of a.a. 1-25 in peptide array overlays, where solid support-spotted peptides may be less accessible to Gγ lipid moiety. Additionally, Gγ_prenyl_ also interacted with hydrophobic a.a. in the C-terminus of Gβ (Supplementary Fig. 10c). If PM were present, it would attract Gγ_prenyl_ and reduce its contact time with channel parts; but binding site mapping would remain unaffected.

**Fig. 8.**
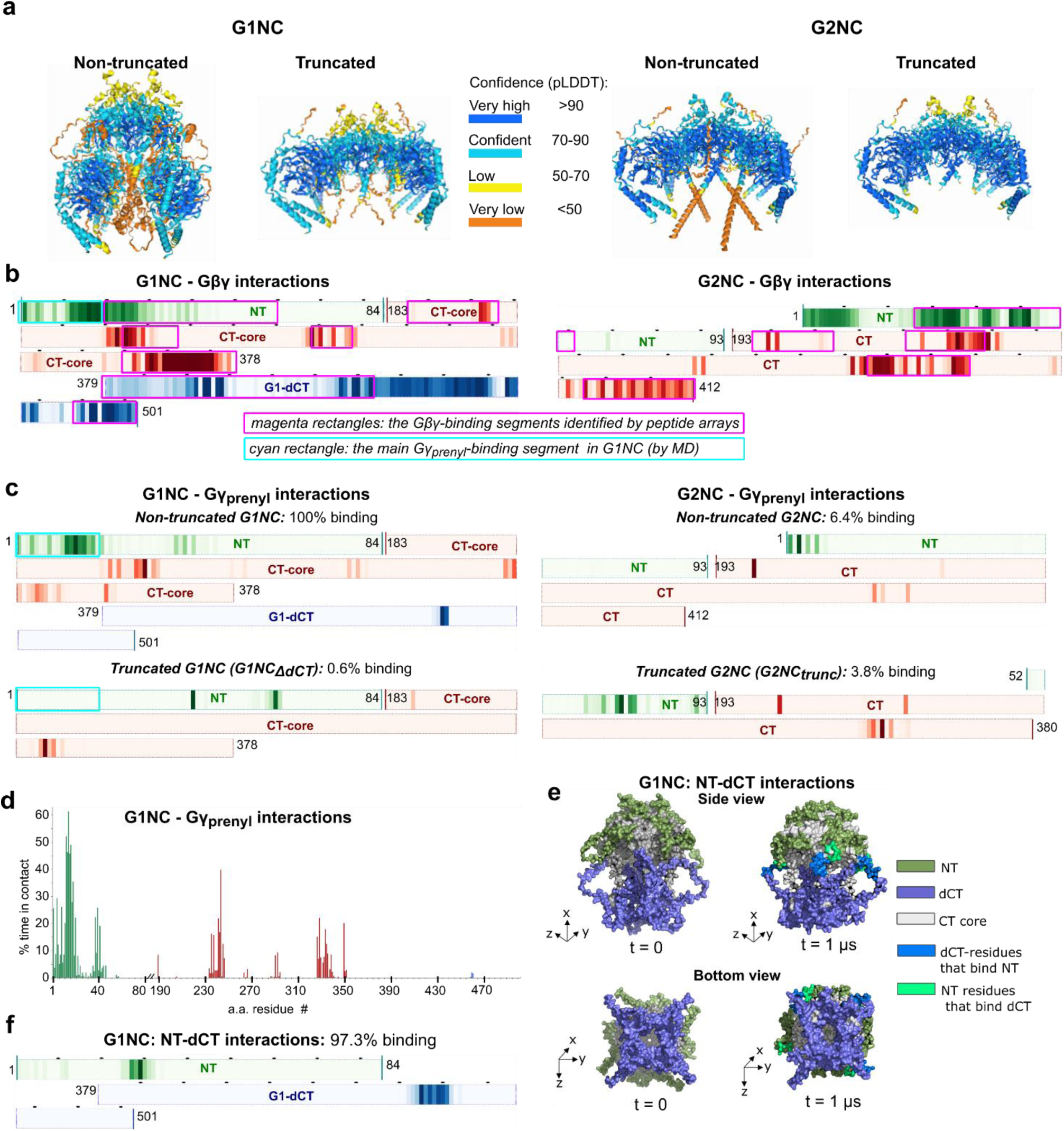
MD simulations corroborate the role of G1-NT and G1-dCT in interactions with Gβγ and the prenylation tail, Gγ_prenyl_. **a**, the initial AlphaFold 3 models of complexes of G1NC and G2NC with prenylated Gβγ. **b,** heatmaps illustrating the G1NC and G2NC residues contributing to Gβγ binding. Analysis was carried out on five 1-µs production runs for G1NC and ten for G2NC. Black dots above the bars are placed at 10 a.a. intervals. Darker coloring corresponds to greater overall contacts between the channel and Gβγ across all production runs. The magenta rectangles superimposed onto the heatmaps correspond to the Gβγ-binding segments identified by the peptide arrays (Fig. 6). The cyan rectangle outlines the main Gγ_prenyl_-binding segment, the beginning of G1-NT. **c**, heatmaps of interactions of G1NC and G2NC and their truncated versions with Gγ_prenyl_. % binding is the percentage of time when at least one prenylation tail is bound to the channel. Note that the Gγ_prenyl_ interaction with the most prominent site, a.a. 1-20 of G1-NT (cyan rectangle), is lost after G1-dCT removal. **d**, the histogram shows % of time spent by G1NC a.a. residues in contact with the Gγ_prenyl_. **e, f**, the interaction between G1-NT and G1-dCT in G1NC. A frame with a contact was defined as one in which at least one G1-dCT chain is bound to the G1-NT, with a cutoff of 6 Å. G1-NT andG1-dCT were in contact in 97.3% of the frames in the five runs. The structures of G1NC (e) are shown at the beginning and at the end (1 µs) of a representative run. Areas of contact are highlighted. The heatmap (f) indicates that the main interaction segment in G1-NT is a.a. 25-32.

Remarkably, deleting dCT abolished Gγ_preny_-G1-NT binding (Fig. 8c), reinforcing the idea that G1-NT and G1-dCT form a Gβγ-binding unit. Truncating G2NC reduced the Gγ_prenyl_ binding by ∼40%. MD simulations also revealed details of the GIRK1’s NT-dCT structural unit, with segments of a.a. 27-31 (NT) and ∼450-460 (dCT) interacting 97.3% of the simulation time (Fig. 8e,f). Notably, this NT-dCT unit is not predicted in the G1NC-Gβγ model by AlphaFold but assembles dynamically during the simulation.

## Discussion

In this study we address two key issues in the GPCR-Gαβγ-GIRK signaling cascade: the Gβγ-GIRK interaction affinity and the subunit-dependent GIRK-Gβγ preassociation. We hypothesized that Gγ prenylation contributes to Gβγ-GIRK interactions and demonstrated that elimination of prenylation thwarts Gβγ interaction with cytosolic domains of GIRK1 and GIRK2 (G1NC and G2NC; Fig. 1). Expectedly, PM targeting was also abolished (Supplementary Fig. 1). However, since our Gβγ binding assays were performed in membrane-free detergent solutions, membrane targeting was not involved. The importance of Gγ_prenyl_ in full channel context in PM is supported by higher GIRK2-Gβγ affinity in intact oocytes (Figs. 3, 4) compared to non-prenylated Gβγ in bilayers^10^. We conclude that, besides its well-established role in membrane attachment of Gβγ, Gγ prenylation enhances Gβγ-GIRK interaction, as in other Gβγ binding partners^39–45^. The mechanism could involve transient interactions of Gγ_prenyl_ with hydrophobic sites in Gβγ’s partner^60^ or Gβ itself, stabilizing the conformation favoring Gβγ function^41, 43, 45, 61^. In support, MD simulations reveal interactions of Gγ_prenyl_ with both, specific sites in GIRK1 and GIRK2, and C-terminal hydrophobic residues of Gβ (Fig. 8, Supplementary Fig. 10).

The dual role of Gγ prenylation complicates the interpretation of *in vitro* affinity measurements. Measuring GIRK’s K_d_ in excised PM patches with prenylated Gβγ in bath solution grossly overestimates affinity (K_d_=2-11 nM, Supplementary Table 8) due to Gβγ’s preferential partitioning to the PM. We addressed the challenge of quantitating GIRK activation by prenylated Gβγ in intact cells utilizing *Xenopus* oocytes, which are exceptionally suitable for accurate titration and monitoring of expression and function of GPCRs, transporters and ion channels^38, 62^. We constructed Gβγ-GIRK dose-response relationships by varying Gβγ expression and measuring surface densities of Gβγ and GIRK responses. Our results support the WTM model^10, 15^ of collision-coupled, cooperative activation of GIRK2 by four Gβγ molecules. However, our affinity estimates are substantially higher.

K_d_ estimates rely on accurate calibrations used to measure Gβγ surface levels. We validated our CF calibrations using two molecular calipers, YFP-GIRK1/2 and IRK1-YFP (Fig. 2). These results, along with previous compatibility tests between CF and qWB methods^38^, enhance confidence in both calibration procedures. The CF approach is advantageous for measuring protein expression and function in individual, intact cells but requires using fluorescently labeled proteins. Disappointingly, xFP-Gβ constructs poorly activated GIRKs, especially GIRK2, calling for caution in using xFP-labeled Gβ in functional studies. Consequently, in most dose-response experiments we used Gβ·_YFP_Gγ, which activated GIRKs like WT Gβγ. When expressing Gβ·_YFP_Gγ, the surface densities of Gβ and YFP-Gγ were linearly related, but measuring YFP-Gγ might overestimate coexpressed Gβ, and accordingly the K_d_, by up to 2.5-fold (Fig. 2). To avoid overinterpretation, we did not apply the YFP-Gγ correction (for measuring YFP-Gγ as a proxy for Gβγ) in our tables and figures.

Even before formal curve fitting, the Gβγ-GIRK2 dose-responses clearly show that only 10 to 150 µm^-2^ of free Gβγ is needed for 10% to 80-90% GIRK2 activation in intact oocytes (Figs. 3, 4), much less than the >1200 µm^-2^ predicted by bilayer results^10^. Applying the 2.5-fold YFP-Gγ correction shifts the activation range to 4-60 µm^-2^. We propose that the higher affinity that we find is mainly due to Gγ prenylation. Truncation of G2NC somewhat reduces Gβγ binding (Supplementary Fig. 2), but the functional impact appears minor (Supplementary Fig. 5).

Interestingly, GIRK2’s I_evoked_ (via m2R) is only 10% of Gβγ-evoked (Supplementary Fig. 1a). Thus, activation of endogenous G_i/o_ (Gα_i/o_βγ) releases 10-15 molecules/µm^-2^ of free Gβγ, corresponding to 30-50% of total endogenous Gβγ in oocyte’s PM, ∼30 µm^-2^ (Fig. 2). Importantly, coexpressing Gα_i3_ and Gβγ with m2R yields I_evoked_ matching I_βγ_^63^. Clearly, endogenous G_i/o_ is insufficient to activate all GIRKs; but m2R can activate all channels when enough G_i/o_ is present.

Comparing K_d_ for a multistep cooperative reaction is complex, even with the same kinetic model. The K_d_ derived from dose-response data is interdependent with the Gβγ cooperativity factor µ: higher µ gives a lower K_d_. µ is Na^+^-dependent^10^ but can be considered constant at stable cytosolic [Na^+^]^15^. (We consider [Na^+^]_in_ in oocytes, 10-20 mM, as close to saturating for GIRK2).

Our average K_d_ estimates for GIRK2 are 11 µM with µ=0.44 (from Fig. 3) and 31 µM with µ=0.3^15^. These are likely overestimates, for two reasons. First, Hill and WTM models assume ligand excess over receptors. This is uncommon in cellular protein-protein interactions, leading to ligand depletion and K_d_ overestimation: more receptors (GIRK) mean less free ligand (Gβγ) per receptor^64^. This is relevant to our whole-cell experiments, where GIRK2 surface density was 17±5 µm^-2^ (Supplementary Table 6), comparable to the functional Gβγ range. Second, applying the 2.5-fold YFP-Gγ correction would shift K_d_ to 4-12 µM, quite close to the most accurate *in vitro* measurement available for prenylated Gβγ, 0.8 µM (interaction with CT of GIRK4, by surface plasmon resonance)^24^.

Notably, less Gβγ is needed for GIRK1/2; 50 µm^-2^ yields full activation (Fig. 4), confirming previous results^38^. The 10-15 Gβγ molecules/µm^2^ released by GPCR activation would yield I_evoked_ of about 50% of I_βγ_ (Fig. 4), consistent with experiments^38^. Accordingly, GIRK1/2’s apparent K_d_ from WTM fits is 5-6-fold lower than GIRK2’s. We further analyzed the GIRK1/2 dose-response data by including explicit calculations of Gα and Gβγ needed to produce the observed I_basal_ and I_evoked_^17, 38^. Across a broad K_d_ range (0.1 to 10 µM), the two cooperative models (Fig. 4g) predicted that both I_basal_ and I_evoked_ could be generated by physiologically relevant amounts of 1-2 Gα and 3-4 Gβγ per channel (Supplementary Fig. 8). This corresponds to less than 40 µm^-2^ of Gβγ assuming physiological densities of GIRKs (2-10 µm^-2^)^16, 28^.

GIRK1’s Gβγ docking site (anchor) emerges as the major factor determining the higher affinity of GIRK1/2. This is suggested by (i) the 4-9-fold affinity drop in GIRK1_ΔdCT_/2, which lacks the main part of the anchor, G1-dCT^36^ (Fig. 4); (ii) the fast deactivation after patch excision of GIRK1_ΔdCT_/2, mirroring GIRK2, indicating faster Gβγ dissociation (Fig. 5). These results, along with the preservation in GIRK1_ΔdCT_ of Asn-217 that renders GIRK1 Na^+^-insensitive^65^, imply a minor role for the differences in Na^+^-dependence of Gβγ affinity in GIRK1 and GIRK2^27^ in our experiments.

The anchor probably increases the apparent affinity through local enrichment of Gβγ (see below). We proposed that Gβγ-anchors are distinct from the Gβγ-binding “activation” sites, which induce channel opening^4^ and are located at the interface between core-CTs of two adjacent GIRK subunits^5, 66, 67^. Removal of G1-dCT preserves maximal Gβγ activation and P_o_ but eliminates Gβγ recruitment and high I_basal_^36^, suggesting functional separation of docking and activation. Structural separation is suggested by strong Gβγ binding to G1NC that persists after removing major components of the activation site (C1-C3) and even the whole core-CT, leaving only the fused NT and dCT (G1NdCT) (Fig. 7). Thus, the anchor dominates the overall Gβγ affinity of GIRK1’s cytosolic domain and does not include elements from core-CT. Both G1-NT and G1-dCT bind Gβγ^59, 68^ (Fig. 6) but much weaker than their fusion protein, G1NdCT (Fig. 7). These results suggest that the Gβγ-anchor is formed jointly by G1-NT and G1-dCT. Interestingly, adding G1-dCT to GIRK2 increased I_basal_ and conferred Gβγ recruitment^36^, suggesting that G2-NT may form Gβγ anchors with G1-dCT.

Peptide array scan and MD simulations provide additional insights. Both approaches identify known Gβγ-binding sites in core-CT, and new NT and dCT Gβγ-binding sites in GIRK1 and GIRK2. Our MD analysis used AlphaFold-models including unstructured but essential elements absent from crystal structures: Gγ_prenyl_ and GIRKs’ NT and dCT. Despite the low-confidence of AlphaFold predictions for these elements, MD calculates interactions based on physical parameters and can capture dynamic interactions even if the initial structure is uncertain. Importantly, the simulations reveal a dynamically arising structural unit formed by G1-NT and G1-dCT, and extensive interactions of Gγ_prenyl_ with G1NC, particularly G1-NT, and some with G2NC (Fig. 8). Remarkably, Gγ_prenyl_–G1-NT interaction is lost, and Gβγ–G1-NT interaction is reduced after deleting G1-dCT, although G1-dCT itself barely interacts with Gγ_prenyl_. These results corroborate the idea that the Gβγ-anchor is a standalone structural and functional unit formed by G1-NT and G1-dCT, with G1-dCT essential for its integrity. Notably, Gγ assists Gβ in GIRK activation^52, 61^. Gγ_prenyl_-anchor interaction may also be involved, since removing G1-dCT or Gγ’s C-terminal region, which includes the prenylation site, eliminates Gγ’s enhancing effect^52, 61^.

Fig. 9 summarizes our view of Gβγ-GIRK2 vs. Gβγ-GIRK1/2 interactions, gating, and the anchor’s role. The dynamic equilibrium between channel-bound, membrane-associated and cytosolic Gβγ determines the local Gβγ concentration within the channel’s microdomain. Free Gβγ can reversibly partition from the cytosolic reserve to the PM, activating GIRKs. Comparing K_d_ for GIRK1/2 activation by Gβγ in whole oocytes (Fig. 4f) and excised oocyte’s patches^46^ yields a Gβγ PM/cytosol partition coefficient between 140 and 425 (Supplementary Fig. 11), close to earlier estimates of ∼300^69^.

**Fig. 9.**
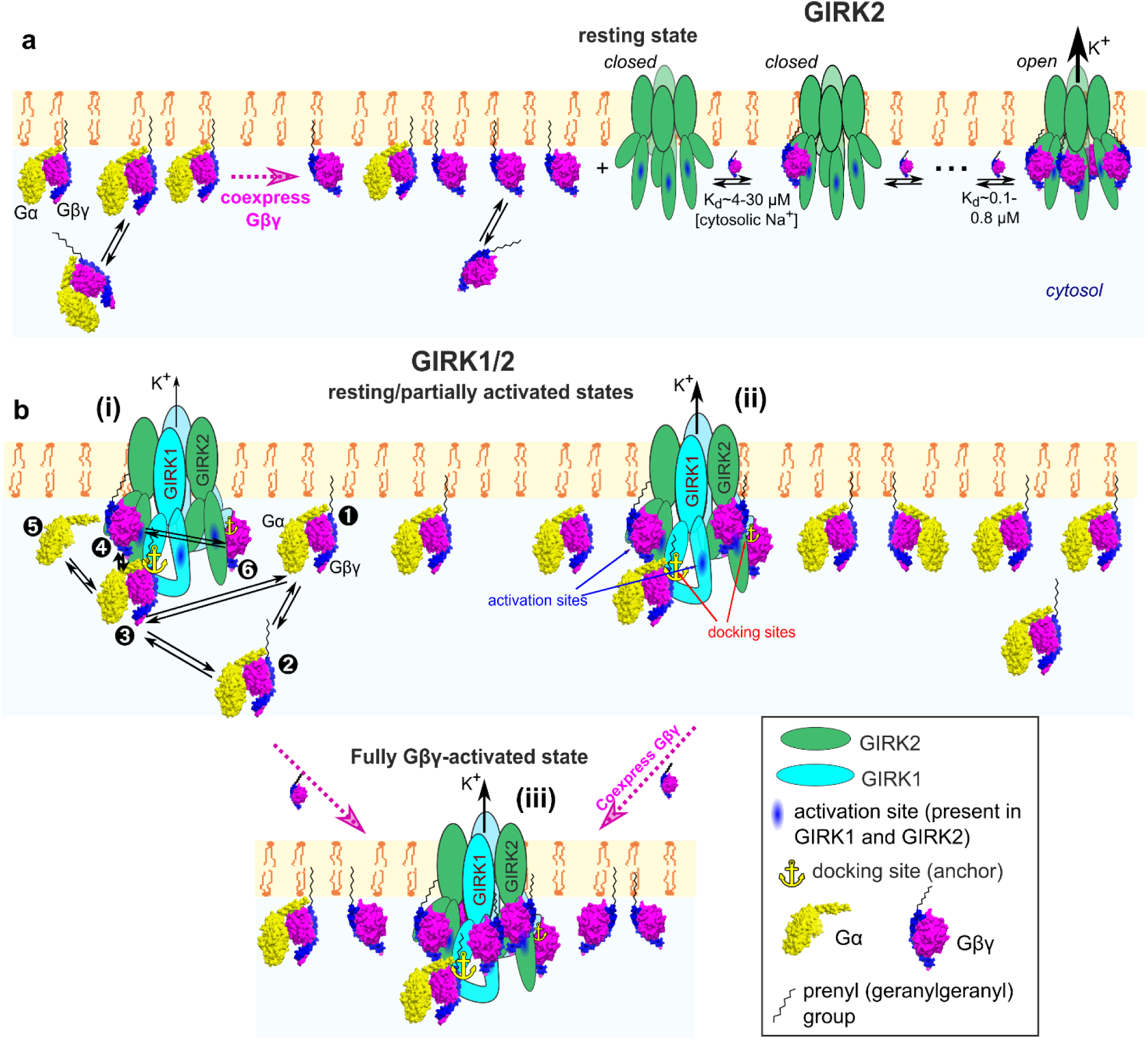
Differences between GIRK2 and GIRK1/2 in their interaction and gating by Gβγ. **a**, GIRK2 homotetramer does not preassociate with Gβγ and has low I_basal_. Channel opening requires the binding of four Gβγ. The affinity of first Gβγ binding is ∼4-30 µM and increases with the binding of each additional Gβγ. **b**, GIRK1/2 reversibly preassociates with Gβγ or Gαβγ due to two Gβγ-docking sites (anchors) formed by G1-dCT and NT (❸,❻) and opened following Gβγ binding to its activation sites (e.g. ❹). In the “graded contribution” scenario shown, binding of even one Gβγ to an activation site induces opening, and P_o_ as well as K^+^ flux are increased with each additional bound Gβγ. GIRK1/2 operates within a complex dynamic system that includes the channel and membrane-associated (❶), cytosolic (❷) and channel-bound Gαβγ and Gβγ, and free Gα_GDP_ or Gα_GTP_ (❺). Gγ_prenyl_ plays an important part in the emerging equilibrium by interacting with the PM or, alternatively, Gα, the anchor, and Gβ C-terminus (most of these interactions are not shown). The anchors attract Gβγ, leading to an enrichment of Gβγ and, potentially, Gαβγ in channel’s microenvironment even in the absence of GPCR activation (basal states i, ii). Free Gβγ is in excess over Gαβγ because the presence of the anchor renders the channel with an overall higher affinity to Gβγ than Gα. Because of excess of free Gβγ, 1-3 out of the 4 activation sites of the GIRK1/2 tetramer are already occupied by Gβγ in basal state, I_basal_ is high, and full activation (state iii) is achieved by binding of additional 1-3 Gβγ molecules.

Our findings confirm that GIRK2 is gated through collision-coupling with Gβγ, cooperative Gβγ binding, and concerted activation after Gβγ occupies four activation sites (Fig. 9a), consistent with the WTM model^10, 15, 16^. However, in intact *Xenopus* oocytes (at estimated [Na^+^ ] of 10-20 mM), the Gβγ-GIRK2 affinity is significantly higher than the bilayer estimates, primarily due to Gγ prenylation, which enhances Gβγ functionality and interaction with GIRKs. The high affinity guarantees efficient G_i/o_-GIRK2 signaling without the need for obligatory hotspots to account for physiological response (although we cannot exclude hotspots or crowding in oocyte PM, which could upshift our K_d_ estimates).

GIRK1/2, on the other hand, operates within a more complex dynamic system featuring two kinds of binding sites, docking (Gβγ-anchors) and activating. The anchor is formed jointly by G1-NT and G1-dCT and is functionally and topologically separate from the activation sites. The similarity of K_d_ values and deactivation rates in GIRK1_ΔdCT_/2 and GIRK2 indicates that the activation sites in GIRK2 and GIRK1/2 have similar Gβγ affinities. If the anchor does not participate in channel opening, how does it increase the apparent affinity? We propose that this is the consequence of the local enrichment of Gβγ around GIRK1/2 due to Gβγ recruitment^4^, through kinetic scaffolding-like mechanisms^70–73^, functionally equivalent to dynamic preassociation. The increased local Gβγ concentration, in excess over Gα, leads to partial occupation of the activation sites and high I_basal_^36, 38^. Moreover, the added Gβγ will bind to the subsequent (unoccupied) sites with higher affinity due to cooperativity, explaining the leftward shift in GIRK1/2’s Gβγ dose-response curve. Added efficiency could arise if Gβγ’s binding surfaces for docking and activation sites are non-overlapping, allowing the docked Gβγ to repeatedly contact the nearby activation site before Gβγ dissociation from the anchor. Mapping the anchor-Gβγ interface is a challenge for the future.

What is the role of Gα? Gα_i/o_ interacts with GIRKs and has been hypothesized to dock the Gα_i/o_βγ heterotrimer to GIRKs^32, 46, 74^. However, the affinity of Gα to GIRK1 is lower than Gβγ^75, 76^. Importantly, binding of Gα_i_ to G1NC is enhanced by added Gβγ, suggesting that the heterotrimer is docked via Gβγ^4, 37, 75^. Both Gβγ-dependent Gα_i3_-GIRK1 interactions and the speed and amplitude of I_evoked_ are maximized when both G1-NT and G1-dCT are present^31, 55, 56, 75^, indicating that the NT-dCT anchor is involved in docking the heterotrimer (Fig. 9b). The stoichiometry of anchor-associated Gαβγ and Gβγ in heterologous models and neurons likely varies with GIRK1/x density, constitutive GPCR activity, and other factors^4^.

### Summary

We quantitated the interaction between Gβγ and GIRK channels in live *Xenopus* oocytes by combining accurate titrated protein expression, PM level monitoring and concurrent functional assays with biochemical and computational approaches. We discovered a novel role for Gγ prenylation in Gβγ-GIRK interaction, resolved the controversy over interaction affinity, and determined the composition of GIRK1’s unique Gβγ-docking site. Our findings reveal efficient, subunit-specific GIRK regulation by Gβγ and will facilitate further quantitative and structure-function analysis of this important signaling cascade.

## Methods

### Ethical approval and Xenopus oocytes handling

Experiments have been approved by Tel Aviv University Institutional Animal Care and Use Committee (permit #01-20-083). Maintenance and surgery of female frogs were done as described^46^. Female frogs, aged 1.5-5 years, were kept at 20 ± 2°C at 10/14-hour light-dark cycle. During surgeries, frogs are anesthetized with a 0.25% Tricaine methanesulfonate (MS-222, Sigma-Aldrich #886-86-2) solution, and parts of ovary are removed through a small abdominal incision. Oocytes were defolliculated with collagenase in Ca^2+^ free ND96 solution (in mM: 96 NaCl, 2 KCl, 1 MgCl_2_, 5 HEPES, pH 7.5). 2 hours later oocytes were washed with NDE solution (in mM: 96 NaCl, 2 KCl, 1 MgCl_2_, 1 CaCl_2_, 5 HEPES, 2.5 mM sodium pyruvate, 50 mg/ml gentamycin, pH 7.5) and left in NDE for 2-24 hours before injection. Oocytes were injected with 50 nl RNA using microinjection pipette (Drummond Scientific, Broomall, PA, USA) and incubated at 20°C for 72 hours for two-electrode voltage clamp, or 48-72 hours for single-channel patch clamp experiments.

### DNA constructs, RNA, antibodies

DNA constructs are summarized in Supplementary Table 9. Antibodies are described in relevant sections of the Methods and summarized in Supplementary Table 10. Gβγ stands for Gβ_1_γ_2_ throughout the paper. All DNA constructs used to produce RNA were inserted in vectors containing 5’ and 3’ untranslated sequences of *Xenopus* β-globin (pGEM-HE, pGEM-HJ or pBS-MXT)^75^. New constructs were prepared using standard PCR-based procedures and fully sequenced. We used the mouse isoform GIRK2A, which is 11 a.a. shorter than the longer isoform (mouse and human) not studied here, which includes a PDZ-binding consensus sequence at the dCT^77^. The truncated GIRK2 construct (GIRK2_trunk_) was prepared by deleting a.a. 1-51 and 381-414 from the GIRK2A construct by PCR. G2NC_trunc_ was prepared by deleting, from G2NC, of the same regions. Myristoylated Gβ_1_ (myr-Gβ) was created by adding the myristoylation signal (the first 15 aa of Src added to the N terminus of Gβ_1_)^35^. GIRK2-CFP was created by fusing CFP_A207K_ to the CT of GIRK2 via a Ser-Arg linker. IRK1-YFP and IRK1-CFP were created by fusing YFP_A207K_ and CFP_A207K_, respectively, to the CT of IRK1 via a Lys-Leu linker, as described^78^. N-terminally Split Venus labeled Gβ_1_ (SpV-Gβ) and N-terminally Split Venus labeled Gγ_2_ (SpV-Gγ)^63^ were subcloned into pGEM-HJ. G1NdCT (the fused cytosolic G1-NT and G1-dCT), G1N(1-40)dCT (the first 40 a.a. of G1-NT fused to G1-dCT), G1N(40-84)dCT (the last 44 a.a. of G1-NT fused to G1-dCT), Sumo-G1NT and Sumo-G1dCT (Sumo fused to G1-NT or G1-dCT). In all cases the fusion was via the 8-a.a. linker, QSTASQST. The Sumo construct used here was a truncated version of human Sumo 2 protein (a.a. 3-95; PDB: 5ELU_B).

RNAs were transcribed *in vitro* as described^46^. The amounts of injected RNAs varied according to the experimental design. For whole-cell electrophysiology experiments we used, in ng/oocyte: 0.01-1 of GIRK1 or YFP-GIRK1, 0.2-10 GIRK2, 0.2-10 Gβ, 0.04-2.5 Gγ, 0.08-5 YFP-Gγ. Equal amounts of GIRK1 and GIRK2 RNAs were injected to express GIRK1/2 channels. In all experiments where several Gβγ expression levels were tested, the ratio of Gβ:Gγ RNA was kept constant: for Gβ:Gγ, the RNA ratio was 5:1 or 2.5:1, and for Gβ:YFP-Gγ the ratio was 2:1 or 2.5:1. For single channel patch clamp, the injected RNAs (in ng/oocyte) were: 0.005-0.01 IRK1-CFP; for GIRK2 alone, 0.02-0.05; for GIRK1/2_HA_, GIRK1 0.01-0.02 of GIRK1 with 0.01-0.02 GIRK2_HA_. In the experiments of Fig. 5, we injected, in ng/oocyte: GIRK2 alone, 0.2-0.5; GIRK1/2, 0.02-0.05 of GIRK1 and 0.01-0.025 of GIRK2; for GIRK1ΔdCT/2, 0.02-0.05 of GIRK1ΔdCT and 0.01-0.025 of GIRK2. In all patch clamp experiments with Gβγ-activated GIRKs, we injected 5 ng Gβ_1_ and 1-2 ng Gγ_2_ RNA, and 25-50 ng/oocyte of the GIRK5 antisense oligonucleotide^38^ to prevent the formation of GIRK1/5 channels.

### Gβγ expression and purification

The pFastBac Dual vector (Invitrogen) was utilized to coexpress the bovine Gγ_2_ (WT and C68S) and Gβ_1_ genes in Sf9 insect cells. The bovine Gγ2 gene was subcloned downstream of a His-tag and a Tobacco Etch Virus (TEV) protease recognition site in the first multiple cloning site, under the control of the polyhedrin promoter. The Gβ1 gene was inserted into the second MCS, under the control of the p10 promoter.

His_6_-Gβγ and His_6_-Gβγ_C68S_ were purified essentially as described^79^. The pFastBac construct was transformed into DH10Bac competent E. coli to generate a recombinant bacmid. The recombinant bacmid was isolated and used for transfection to SF9 cells in presence of CellFectin II reagent (Thermo Fisher Scientific) to generate recombinant baculovirus. Further, this baculovirus stock was amplified and optimized for maximum protein expression via infection to Trichoplusia ni (T.ni) cells. The infected T.ni cells were grown for 60-72 hrs. Infected T.ni cells were harvested by centrifugation at 1000 rpm and stored at -80°C until further use.

Purification of Gβγ: Cells were suspended and homogenized using glass homogenizer in 20 mM Hepes pH 8.0, 100 mM NaCl, 3 mM MgCl2, 100 mM EDTA, 10 mM 2-mercapto ethanol, cocktail protease inhibitors, 1% Triton X-100. Then the cells were lysed in sonicator using a program 5/25 sec on/off pulse for 30 min and subjected to centrifugation at 42000 rpm for 45 min. The soluble fraction was filtered using 0.4 µM filter and His-tag Gβγ was purified by sequential Ni^2+^ chelate, size-exclusion [Superdex-75 HiPrep (GE Healthcare)] column chromatography. Final buffer conditions were: 20 mM Hepes pH 8.0, 100 mM NaCl, 1 mM MgCl_2_, 10 mM 2-mercaptoethanol. The fraction purity was analyzed using SDS-PAGE. The protein was also characterized by Western blot using anti-Gβ (GTX114442) and anti-His tag (Roche 11 965 085 001) antibody (Supplementary Table 10).

### Electrophysiology

Whole-cell GIRK currents were measured using standard two-electrode voltage clamp at 20-22°C using GeneClamp 500B amplifier (Molecular Devices, Sunnyvale, CA, USA) and digitized using Axon Digidata 1440a using pCLAMP software (Molecular Devices). Agarose cushion microelectrodes were filled with 3M KCl, with resistances of 0.1–1 MΩ^37^. GIRK currents were measured in either low-[K^+^] solution ND96 (same as Ca^2+^-free but with 1 mM CaCl_2_) or high-K solution with 24 mM [K]_out_ (in mM: 24 KCl, 72 NaCl, 1 CaCl_2_, 1 MgCl_2_ and 5 Hepes). In experiments of Fig. 1, to maximize GIRK2’s I_basal_, we used a 96 mM high-[K]_out_ solution (in mM: 96 KCl, 2 NaCl, 1 CaCl_2_, 1 MgCl_2_ and 5 Hepes). Net GIRK currents (I_basal_ and I_βγ_) were determined by subtraction of currents recorded in presence of 1-2.5 mM Ba^2+^ that blocked GIRK currents. The pH of all solutions was 7.5–7.6. Cell-attached patch clamp recordings were performed as previously described^38^, at 20–23°C, using borosilicate glass pipettes with resistances of 1.5–3.5 MΩ. The electrode solution contained (in mM): 146 KCl, 2 NaCl, 1 CaCl_2_, 1 MgCl_2_, 10 Hepes and 1 GdCl_3_ (pH 7.6). Bath solution contained (in mM): 146 KCl, 2 MgCl_2_, 6 NaCl, 10 Hepes and 1 EGTA (pH 7.6). Block of stretch-activated channels by GdCl_3_ was confirmed by recording currents at +80 mV. Single channel currents were recorded at −80 mV in cell-attached patches with the Axopatch 200B amplifier (Molecular Devices) at −80 mV, filtered at 2 or 5 kHz and sampled at 10 or 25 kHz.

### Giant membrane patches (GMPs)

GMPs were prepared and imaged as described^63^. Oocytes were devitellinized using tweezers in hypertonic solution (in mM: 6 NaCl, 150 KCl, 4 MgCl2, 10 Hepes, pH 7.6). The devitellinized oocytes were transferred onto a Thermanox^TM^ coverslip (Nunc, Roskilde, Denmark) and immersed in Ca^2+^-free ND96 solution with their black hemisphere facing the coverslip, for 30– 45 min. The oocytes were then suctioned using a Pasteur pipette, leaving a GMP attached to the coverslip, with the cytosolic part facing the medium. The coverslip was washed thoroughly with fresh ND96 solution, and fixated using 4% formaldehyde for 30 min. Fixated GMPs were immunostained in 5% milk in PBS and non-specific binding was blocked with Donkey IgG 1:200 (Jackson ImmunoResearch, West Grove, PA, USA). Primary rabbit anti-Gβ (1:200; Santa Cruz, SC-378 or GeneTex, GTX114442) was applied for 45 min at 37°C either alone or with blocking peptide supplied with the antibody. Then DyLight549 or DyLight® 650-conjugated anti-rabbit secondary antibodies (KPL) were applied at 1:300 dilution for 30 min at 37°C, washed with PBS and mounted on a slide for visualization. Immunostained slides were kept at 4°C in the dark.

### Pull-down assay

Pull-down binding experiments were performed as described^36^. Briefly, in vitro translated (*ivt*) [^35^S]methionine-labelled proteins were prepared in rabbit reticulocyte lysate (Promega, Madison, WI, USA). *Ivt* proteins were mixed with ∼2 µg of either purified His-Gβγ_WT_ or purified His-Gβγ_C68S_ in 300 μl of the incubation buffer (in mM: 150 KCl, 50 Tris, 0.6 MgCl_2_, 1 EDTA, 0.1% Lubrol or 0.5% CHAPS and 10 imidazole; pH 7.4). The mixture was incubated while shaking for 45 min at room temperature, then 30 μl beads were added, and incubated for 30 min at 4°C. His-Gβγ was pulled-down using HisPurTM Ni-NTA Resin affinity beads (ThermoFisher Scientific, Rockford, IL, USA). The beads were washed three times with 500 μl buffer. Elution was done with 30 μl elution buffer (incubation buffer supplemented with 250 mM imidazole). After washing, the samples were analyzed on 12% gels by SDS-PAGE. Also, 1/60 of the mixture before the pull-down was loaded, usually on a separate gel (‘input’). Gels were imaged using Sapphire™ Biomolecular Imager (Azure Biosystems, Dublin, CA, USA). Autoradiograms were analyzed using ImageQuant 5.2 (GE Healthcare) and ImageJ/Fuji (https://imagej.net/software/fiji/). Binding was calculated as percentage or fraction of the input of this construct in the same experiment.

### Confocal imaging

Confocal imaging and analysis were performed as described^78^, with Zeiss 510, Ziess 710 or Leica TCS SP5 confocal microscopes, using a 20× objective. Live oocytes were images at their animal hemisphere in ND96 or NDE solutions. Giant membrane patches were imaged at their edges, so both the membrane and the background were visible. Images were acquired using spectral (λ)-mode or channel mode. For imaging the following wavelength parameters were used: YFP, excitation 514 nm, imaging at 525–540 nm; DyLight 650, excitation 633 nm, imaging at 663-673 nm. For whole oocytes, fluorescence signals at the maximum emission wavelength are averaged from three regions of interest using Zeiss LSM Image Browser (Carl Zeiss Jena GmbH), ImageJ or LAS AF (Lecia Microsystems CMC GmbH) Image software. For giant membrane patches, the entire visible membrane and background were averaged. Averaged background for each signal, and the average net signal from uninjected (native) oocytes of the same experiment, were subtracted to obtain net signal. For the effect of Gβγ on the expression of GIRK2 and GIRK2HA, signals were normalized by dividing the signal of each oocyte with the average of the group that did not contain Gβγ.

### Western blots of Gβ in oocytes’ plasma membranes

Quantitative Western blots of Gβ in manually separated oocytes’ PMs have been performed as described^38^. Briefly, PMs together with the vitelline membranes (extracellular collagen-like matrix) were manually separated from the rest of the oocyte (“cytosol”) with fine forceps, after a 5-15 min incubation in a low osmolarity solution (5 mM NaCl, 5 mM HEPES, and protease inhibitors (Roche Complete Protease Inhibitors Cocktail (Merck), 1 tablet/50 ml, pH=7.5). PMs of ∼20 oocytes were pooled for each sample (lane on polyacrylamide-SDS gel). The cytosol was processed separately: the nuclei were separated by centrifugation for 10 min at 700×g at 4°C and removing the pellet (nuclei). Plasma membranes and cytosols were then solubilized in 35 µl running buffer (2% SDS, 10% glycerol, 5% β-mercaptoethanol, 0.05% Bromophenol Blue, 62.5 mM Tris-HCl pH 6.8) and heated to 65°C for 5 min. Samples were electrophoresed on 12% polyacrylamide-SDS gel and transferred to nitrocellulose membranes for standard Western blotting. Known amounts of purified His-Gβγ were run on the same gel for the construction of the calibration curve. Gβγ was detected with anti-GNB1 antibody (GTX114442) at 1:500 or 1:1000 dilution on Fusion FX7 (Witec AG, Sursee, Switzerland) and quantitated using Fiji software. Signal in each band was assessed as (mean intensity)×(area) using ImageJ/Fiji. To quantify the expressed Gβ, endogenous Gβ signal from oocytes expressing the channel alone was subtracted from the total signal.

### Peptide spot array

Peptide arrays were generated by automatic SPOT synthesis and blotted on a Whatman membrane as described^80^. Briefly, N-terminal and C-terminal parts of GIRK1 and GIRK2 were spot-synthesized as 25-mer peptides overlapping sequences, shifted by 5 a.a. along the sequence, using AutoSpot Robot ASS 222 (Intavis Bioanalytical Instruments, Cologne, Germany). The peptides were designed according to human GIRK2 (NCBI: NM_002240.5) (NT: a.a. 1-93, CT: a.a. 193-423) and rat GIRK1 (NCBI: NP_113798.1) (NT: a.a. 1-84, CT: a.a. 183-501). The interaction with spot-synthesized peptides was investigated by an overlay assay. Following blocking of 1 hour at room temperature with 5% BSA in 20 mM Tris and 150 mM NaCl with 0.1% Tween-20 (TBST), 0.016-0.16 μM purified His-Gβγ were incubated with the immobilized peptide-dots, overnight at 4 °C. His-Gβγ was detected by anti-GNB1 antibody (GTX114442) at 1:500 or 1:1000 dilution, and anti-rabbit HRP-coupled secondary antibody (1:40000) incubated with 5% BSA/TBST, and the membrane was imaged using Fusion FX7, as for Western blotting.

### Electrophysiological data analysis and surface density calibration

Whole-cell and single-channel data were analyzed using Clampex and Clampfit (pCLAMP suite, Molecular Devices, Sunnyvale, CA, USA). In oocytes expressing the m2 receptor, the fold activation by agonist, R_a_, was measured in each cell and defined as

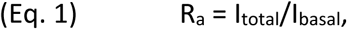

where I_total_=I_basal_+I_evoked_. R_a_=1 when there is no response to agonist.

The fold activation by Gβγ, R_βγ_, was defined as

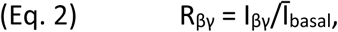

where I_βγ_ is the net GIRK current in a Gβγ-expressing oocyte, and Ῑ_basal_ is the average GIRK current in oocytes of control group, that express only the channel, from the same experiment^37^.

Single channel amplitudes were calculated from Gaussian fits of all-points histograms of 30–90 s segments of the record. The open channel probability (*P*_o_) was estimated from 1–5 min segments of 4–20 min recordings from patches containing one to three channels using a standard 50% idealization criterion^38^.

The PM density of functional channels was determined from the whole-cell current, I, using the classical equation^49^

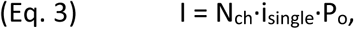

where N_ch_ is the total number of channels in the cell, i_single_ is the single-channel current and P_o_ is the open probability. P_o_ and i_single_ for Gβγ-activated GIRK1/2 are known, and for GIRK2, GIRK1/2_HA_ and IRK1-xFP we determined them here (Supplementary Fig. 3, Supplementary Table 3). The surface density, in channels/µm^2^ (µm^-2^) was calculated by dividing N_ch_ by the membrane surface area of the oocyte^81^, 2·10^7^ µm^2^. Protein surface densities were converted to concentrations using the standard procedure based on a submembrane interaction space 10 nm deep. i_single_ was measured in cell-attached patches in 146 mM [K^+^]_out_, whole-cell currents were measured in 24 mM [K^+^]_out_. The amplitude translation factor for these solutions was 4.63. The conversion factor from surface density to sub-PM space concentration was 1 µm^-2^ = 0.166 µM^38^. In calculating the surface density of channel-attached YFP (two for YFP-GIRK1/2 and four for IRK1-YFP), we assumed similar levels of fluorescence maturation of channel- and Gβ-attached YFP molecules, therefore no correction for such maturation was made. For CF calibrations with YFP-GIRK1/2 or IRK1-YFP, the linear fit included the zero-fluorescence point (with no expressed channels).

In the analysis of Gβγ dose-response data in intact oocytes, we assumed that the PM level of the GIRK2 channels was not significantly altered by Gβγ, as shown previously^37, 63^ and confirmed for CFP-GIRK2 (Supplementary Fig. 6). In one experiment we monitored GIRK2HA and observed changes at different doses of Gβγ, and corrected the currents accordingly (Supplementary Table 6). Similarly, coexpression of Gβγ causes no significant changes in PM levels of GIRK1/2 up to 2 ng RNA of Gβ^63^. In most experiments, the maximal GIRK1/2 current was observed already with 1 or 2 ng Gβ RNA. With 5 ng Gβ RNA, a 20-30% decrease in channel expression is occasionally seen^63^. No correction for this potential change has been made.

### Modeling, simulation and curve fitting for Gβγ dose-response data

Standard fitting for Gβγ-GIRK dose-response curves with Hill or modified WTM models was done assuming that, in the absence of GPCR simulation, the endogenous G proteins are in the form of heterotrimers. Data were fitted to Hill equation in the following form:

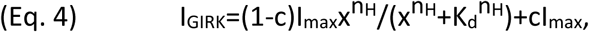

where x is the concentration of coexpressed Gβγ ([Gβγ]), I_GIRK_ is GIRK current, I_max_ is the maximal GIRK current at saturating concentrations of coexpressed Gβγ, h_H_ is the Hill coefficient, c is a constant component corresponding to I_basal_;

or a modified WTM model^15^ with the addition of a constant component c:

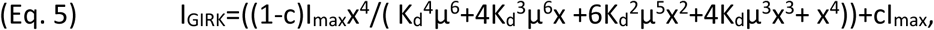

where x, c and I_max_ have the same meaning as in Eq. 4, K_d_ is the dissociation constant of the first Gβγ binding to the one of the four sites in GIRK molecule, µ is the cooperativity factor for each successive Gβγ binding^10^ for the specific case of a constant Na^+^ concentration^15^. In whole-cell of cell-attached recordings from intact *Xenopus* oocytes, both intracellular Na^+^ and the membrane PIP_2_ can be assumed constant during the experiment. Therefore, in most WTM model fits, we utilized a constant cooperativity factor µ=0.3^15^ or µ=0.44 (from Fig. 3). In two experiments with GIRK2 we were able to obtain independent estimates of µ from fit, which were 0.44 and 0.62 (Supplementary Table 6, “free µ”).

To simulate Gβγ activation of GIRK1/2, we tested four kinetic schemes (models) (Supplementary Fig. 8a, Supplementary Methods). First, we calculated the basal available Gβγ and Gα from the experimentally observed I_basal_ as described previously^17, 38^. For simulation, we constructed systems of differential equations based on these schemes and solved them numerically in Berkley Madonna (Berkeley Madonna, Inc., Albany, CA) ^82^ utilizing 4th order Runge-Kutta integration method. The simulations were run till apparent steady state was achieved.

### Molecular dynamics simulations

Primary structures of G1NC and G2NC were generated by fusing the NT and CT of human GIRK1 (UniProt-ID: P48549; a.a.1-84 and 183-501) and human GIRK2 (UniProt-ID: P48051, a.a. 1-93 and 193-414), respectively (Fig. 8). The heatmaps in Fig. 8 show G412 as the last a.a., which corresponds to G414 of mGIRK2 (Supplementary Fig. 9). Additionally, Gβγ (UniProt-ID: P62873, P59768) units were incorporated into the sequences. Similarly, truncated constructs were obtained by omitting a.a. 379-501 and a.a. 1-52 and 381-412 for G1NC and G2NC, respectively. The truncated G1NC and G2NC are the same as G1NC_ΔdCT_ and G2NC_trunc_ used in biochemical experiments (see Fig. 1e). Heterotetramers bound to four Gβγ were modeled using AlphaFold 3^83^. Coarse-grained constructs were modeled with CHARMM-GUI^84^. Simulations were conducted with Gromacs 2022.3^85^ based on the Martini force field Elnedyn22p^86^ and polarizable water^87^ in 100mM KCl. For the geranylgeranyl moiety, previously published parameters were used^88^. Minimization and equilibration procedures followed the CHARMM-GUI protocol. Simulation runs of 1 μs each were performed with a time step of 20 fs. During the production runs, the temperature was set to 310 K using the v-rescale thermostat^89^, and the pressure was kept at 1 bar with the Parrinello-Rahman barostat^90^. Lennard-Jones potentials and Coulombic interactions are switched off between 9 to 12 Å and 0 to 12 Å, respectively. Contacts were analyzed using MDAnalysis^91^ with a distance cut-off of 6 Å.

### Statistical analysis

Statistical analysis was performed using GraphPad Prism (GraphPad, La Jolla, CA, USA). For normally distributed data (by Shapiro-Wilk test), pairwise comparison was done by t-test and multiple comparisons by one-way ANOVA, and data were presented as bar graphs with individual data points and mean ± SEM (except if non-normally distributed data were presented on the same panel, in which case box plots were shown). If the data did not pass the normal distribution test, they were analyzed using Mann-Whitney (pairwise) and Kruskal-Wallis non-parametric ANOVA tests, and data were presented as box plots and individual data points. The boxes represent the 25th and 75th percentiles, the whiskers show the smallest and maximal values, and the horizontal line represents the median. Statistical analysis for differences between dose-response curves for two different GIRK compositions was done on WTM model fits of normalized dose-response data from individual oocytes for two fits (as in Fig. 4b,d), as well on three fits (details in Supplementary Fig. 7).

### Graphics

Structures of GIRK2, Gα and Gβγ were drawn with PyMOL (Schrodinger LLC). All final figures were produced with Inkscape (inkscape.org).

## Acknowledgements

This work was supported by grants 1282/18 and 581/22 from Israel Science Foundation (ND), KL1415/14-1 from the Deutsche Forschungsgemeinschaft (EK), R35GM145921 from the National Institute of General Medical Science, National Institutes of Health, W1232 from the Austrian Science Fund (ASW, TF). The computational results presented have been achieved in part using the Vienna Scientific Cluster (VSC).

## Authors contributions

ND, DY, ASW – conceptualization

RH, TKR, BS, UK, TF, CWD, YH, JAH, ASW, DY, ND – experimental design

RH, TKR, BS, PH, UK, TF, GT, VT, HRP, ASW, DY, ND – investigation

RH, GT, OCH, DRT, KZ, CWD, EK, JAH – resources

RH, THK, PH, UK, TF, ASW, ND – data curation

EK, YH, JAH, ASW, DY, ND – supervision

CWD, EK, YH, JAH, ASW, ND – funding acquisition

RH, ASW, DY, ND – writing the initial version of the paper

RH, HRP, UK, EK, CWD, YH, JAH, ASW, DY, ND - review and editing

All authors read and confirmed the paper.

## Materials & correspondence

All materials created in this paper, such as DNA constructs, are fully available upon reasonable request. Correspondence regarding experimental procedures and results to ND, kinetic modeling and analysis to DY, molecular dynamics simulations AWS.

## Data availability

All data are presented in figures and tables in the main paper and in Supplementary Material. MD simulation data has been stored on Zenodo: doi: 10.5281/zenodo.15075373.

