## Supplementary Material for "Live-cell quantitative monitoring reveals distinct, high-affinity Gβγ regulations of GIRK2 and GIRK1/2 channels"

### ***Supplementary Methods***

*Modeling GIRK1/2 basal activity and activation by expression of Gβγ with four kinetic models*

To describe GIRK1/2 activation by Gβγ with the inclusion of a mechanistic explanation of $I_{basal}$, we utilized 4 gating models (Supplementary Fig. 8a): #1, concerted activation, non-cooperative binding; #2, concerted activation, cooperative binding; #3, graded contribution, non-cooperative binding; and #4, graded contribution, cooperative binding model. Concerted activation models are based on the assumption of four Gβγ required for channel opening. The concerted activation, cooperative binding model #2 is the modified WTM model used throughout this paper. Graded contribution models are based on increasing contribution to $P_{o,max}$ of each sequential Gβγ-bound state. We have previously described the graded contribution non-cooperative binding model[1], and the graded contribution cooperative binding model was described in detail by Berlin et al.[2].

Since the basal activity of GIRK12 is highly Gβγ-dependent, and this phenomenon was shown to be dependent on differential recruitment of Gα and Gβγ to the GIRK1/2 microenvironment[3], we first utilized each of the four models of Supplementary Fig. 8a to estimate the basal endogenous Gα and Gβγ available for channel's gating ($G\alpha_{endo}$ and $G\beta\gamma_{endo}$, where endo stays for endogenous).

For calculations we used the following parameters: $P_{o,max}$ (open probability value, observed with 5 ng Gβ RNA and 1-2 Gγ RNA) ~ 0.105[1], $P_o$ in the absence of Gβγ ~0.00273 (assuming 10% Gβγ-independent activity out of total $I_{basal}$ of GIRK1/2 , $P_{o,basal}$ = 0.0273 (open probability in absence of Gβγ expression, c=26 % of maximal open probability; Fig. 4F) and Po, agonist = 0.0525 (open probability corresponding to full endogenous Gβγ dissociation from Gα induced by agonist, which is 50% of that of $P_{o,max}$ with 5 ng Gβ RNA[1]. Utilizing the $P_{o,max}$ and single channel current, we estimated channel density to be 13.7 channels/$\mu m^2$. The above described data were substituted to system of equations relevant for each model as described[1, 2] and solved in Matlab 6.5 for Windows. This procedure was conducted for a range of $K_d$ values, thus generating initial values matrix containing [$K_d$, $G\beta\gamma_{endo}$, $G\alpha_{endo}$]. Estimated values are shown in Supplementary Fig. 8b. Each [$K_d$, $G\beta\gamma_{endp}$, $G\alpha_{endo}$] was subsequently utilized for simulation of a complete dose response to expressed Gβγ. In order to make calculations more time-efficient, the dose response was simulated as a steady-state solution of a system of differential equations. Gating schemes described in Fig.S8a were combined with agonist-independent G-protein dissociation reaction:

(1) $$G\alpha_{GDP} + G\beta\gamma \underset{k_{off,G}}{\overset{k_{on,G}}{\rightleftarrows}} G\alpha_{GDP}G\beta\gamma$$

where $G\alpha_{endo}=G\alpha_{GDP}+G\alpha_{GDP}G\beta\gamma$ and $G\beta\gamma_{endo}=G\beta\gamma+G\alpha_{GDP}G\beta\gamma$, and $G\alpha_{endo}$ and $G\beta\gamma_{endo}$ are calculated values obtained for each $K_d$ value as described above.

Simulation of models based on cooperative binding of Gβγ to the channel was based on solution of the following differential equations system:

(2) $dG\alpha_{GDP}/dt=k_{off,G}\cdot G\alpha_{GDP}G\beta\gamma-k_{on,G}\cdot G\alpha_{GDP}\cdot G\beta\gamma$

(3) $dG\beta\gamma/dt= k_{off,G}\cdot G\alpha_{GDP}G\beta\gamma+k_{off}\cdot(C_1+2\cdot\mu\cdot C_2+3\cdot\mu^2\cdot C_3+4\cdot\mu^3\cdot C_4)-$

$- G\beta\gamma\cdot(k_{on,G}\cdot G\alpha_{GDP}+4\cdot k_{on}\cdot C_0+3\cdot k_{on}\cdot C_1+2\cdot k_{on}\cdot C_2+ k_{on}\cdot C_3)$

(4) $dG\alpha_{GDP}G\beta\gamma/dt= k_{on,G}\cdot G\alpha_{GDP}\cdot G\beta\gamma- k_{off,G}\cdot G\alpha_{GDP}\cdot G\beta\gamma$

(5) $dC_0/dt= -4\cdot k_{on}\cdot G\beta\gamma\cdot C_0+k_{off}\cdot C_1$

(6) $dC_1/dt=4\cdot k_{on}\cdot G\beta\gamma\cdot C_0-3\cdot k_{on}\cdot G\beta\gamma\cdot C_1-k_{off}\cdot C_1+2\cdot\mu\cdot k_{off}\cdot C_2$

(7) $dC_2/dt=3\cdot k_{on}\cdot G\beta\gamma\cdot C_1-2\cdot k_{on}\cdot G\beta\gamma\cdot C_2-2\cdot\mu\cdot k_{off}\cdot C_2+3\cdot\mu^2\cdot k_{off}\cdot C_3$

(8) $dC_3/dt=2\cdot k_{on}\cdot G\beta\gamma\cdot C_2-1\cdot k_{on}\cdot G\beta\gamma\cdot C_3-3\cdot\mu^2\cdot k_{off}\cdot C_3+4\cdot\mu^3\cdot k_{off}\cdot C_4$

(9) $dC_4/dt=1\cdot k_{on}\cdot G\beta\gamma\cdot C_3-4\cdot\mu^3\cdot k_{off}\cdot C_4$

Simulation for model based on non-cooperative binding of Gβγ to channel was based on solution of the following differential equations system:

(10) $dG\alpha_{GDP}/dt=k_{off,G}\cdot G\alpha_{GDP}G\beta\gamma-k_{on,G}\cdot G\alpha_{GDP}\cdot G\beta\gamma$

(11) $dG\beta\gamma/dt=k_{off,G}\cdot G\alpha_{GDP}G\beta\gamma+k_{off}\cdot(C_1+2\cdot\mu\cdot C_2+3\cdot\mu^2\cdot C_3+4\cdot\mu^3\cdot C_4)-$
$-G\beta\gamma\cdot(k_{on,G}\cdot G\alpha_{GDP}+4\cdot k_{on}\cdot C_0+3\cdot k_{on}\cdot C_1+2\cdot k_{on}\cdot C_2+k_{on}\cdot C_3)$

(12) $dG\alpha_{GDP}G\beta\gamma/dt=k_{on,G}\cdot G\alpha_{GDP}G\beta\gamma-k_{off,G}\cdot G\alpha_{GDP}G\beta\gamma$

(13) $dC_0/dt=-4\cdot k_{on}\cdot G\beta\gamma\cdot C_0+k_{off}\cdot C_1$

(14) $dC_1/dt=4\cdot k_{on}\cdot G\beta\gamma\cdot C_0-3\cdot k_{on}\cdot G\beta\gamma\cdot C_1-k_{off}\cdot C_1+2\cdot k_{off}\cdot C_2$

(15) $dC_2/dt=3\cdot k_{on}\cdot G\beta\gamma\cdot C_1-2\cdot k_{on}\cdot G\beta\gamma\cdot C_2-2\cdot k_{off}\cdot C_2+3\cdot k_{off}\cdot C_3$

(16) $dC_3/dt=2\cdot k_{on}\cdot G\beta\gamma\cdot C_2-1\cdot k_{on}\cdot G\beta\gamma\cdot C_3-3\cdot k_{off}\cdot C_3+4\cdot k_{off}\cdot C_4$

(17) $dC_4/dt=1\cdot k_{on}\cdot G\beta\gamma\cdot C_3-4\cdot k_{off}\cdot C_4$

For models based on graded contribution of each Gβγ occupied state to channel activity the open probability was calculated according to:

(18)

$$P_o=P_{o,max}\cdot\sum_{1}^{4}\varphi_i\cdot\frac{C_i}{C_{total}}$$

where $\varphi_i$ is the contribution factor of each occupied Gβγ state to open probability, and $C_{total}$ is channel concentration ($C_{total}=\sum_{i=0}^{i=4}C_i$). $\varphi_i$ values were adopted from Yakubovich et al.[1] and Berlin et al.[2] and based on data published by Ivanova-Nikolova et al. and Sadja et al.[4, 5].

For models based on concerted gating the open probability was calculated according to:

(19)

$$P_o=P_{o,max}\cdot\frac{C_4}{C_{total}}$$

These models are based on the same assumption as used in WTM model, i.e. only 4 Gβγ-occupied channel is available for opening.

In all models $C_0$-$C_4$ correspond to 0-4 Gβγ occupied state of the channel, $k_{on}$ =1e7 $M^{-1}s^{-1}$ (similar to value utilized by Berlin at al. and in agreement with the diffusion limit[2, 6], $k_{off}=K_d/k_{on}$, μ is the cooperativity factor of Gβγ binding to each consecutive Gβγ occupied state[6]. $k_{on,G}$ =0.7e6 $M^{-1}s^{-1}$ and $k_{off,G}$ =0.0013 $s^{-1}$ as reported by Sarvazyan et al. [7]. For simulation of response to expressed Gβγ initial values of [$K_d$ $G\alpha_{endo}$, $G\beta\gamma_{endo}$+$G\beta\gamma_{expressed}$] were utilized for each consequtive run of differential equation system solution, thus generating matrix of [$G\beta\gamma_{expressed}$ $P_o$] values for each $K_d$. Differential equations systems were solved in Berkeley Madonna for Windows utilizing 4th order Runge-Kutta integration method. All systems were allowed to reach steady-state. The results of simulation were compared to experimental dose-response curves. For selection of optimal [$K_d$ $G\alpha_{endo}$ $G\beta\gamma_{endo}$] values we utilized two criteria: a) the stability of G-protein concentration estimation as seen from Supplementary Fig. 8b – i.e. an optimal model is expected to generate stable estimation of G protein concentration over a wide range of tested $K_d$ values, and b) resemblance to superimposed dose-response by visual inspection. Simulations of all models are shown in Fig. 4g and Supplementary Fig. 8d,e.

*Extracellular HA staining*

Extracellular HA staining of the oocytes was performed as described[8]. Briefly, oocytes were fixated with 4% formaldehyde for 30 min, blocked in 5% milk in $Ca^{2+}$-free ND96 solution for 1 hour, incubated with mouse anti-HA antibody (1:333; Santa Cruz Biotechnology, Dallas, TX, USA) in 2.5% milk-$Ca^{2+}$-free ND96 for 1 hour, washed thrice, and incubated with DyLight405-conjugated anti-mouse antibody (KPL, Gaithersburg, MD, USA). Oocytes were then kept at 4°C in the dark in $Ca^{2+}$-free ND96 solution until imaged.

### *Supplementary Figures*

*Fig. S1*

**a**

| | $I_{\beta\gamma}$, µA | $I_{basal}$, nA | | | $R_a$ | | | $R_{\beta\gamma}$ | | | N |
|---|---|---|---|---|---|---|---|---|---|---|---|
| | | mean | SEM | n | mean | SEM | n | mean | SEM | n | |
| GIRK2 | 2-4 | 64.2 | 7.1 | 49 | 8.2 | 0.8 | 49 | 67.3 | 7.9 | 52 | 7 |
| | >4 | 137.7 | 19.2 | 48 | 5.1 | 0.4 | 48 | 68.9 | 9.3 | 31 | 5 |
| GIRK2HA | 3-10 | 69.8 | 16.4 | 16 | 5.7 | 0.7 | 16 | 78.0 | 9.6 | 21 | 3 |
| | >10 | 348.7 | 34.0 | 23 | 4.9 | 0.7 | 23 | 49.4 | 7.1 | 18 | 3 |

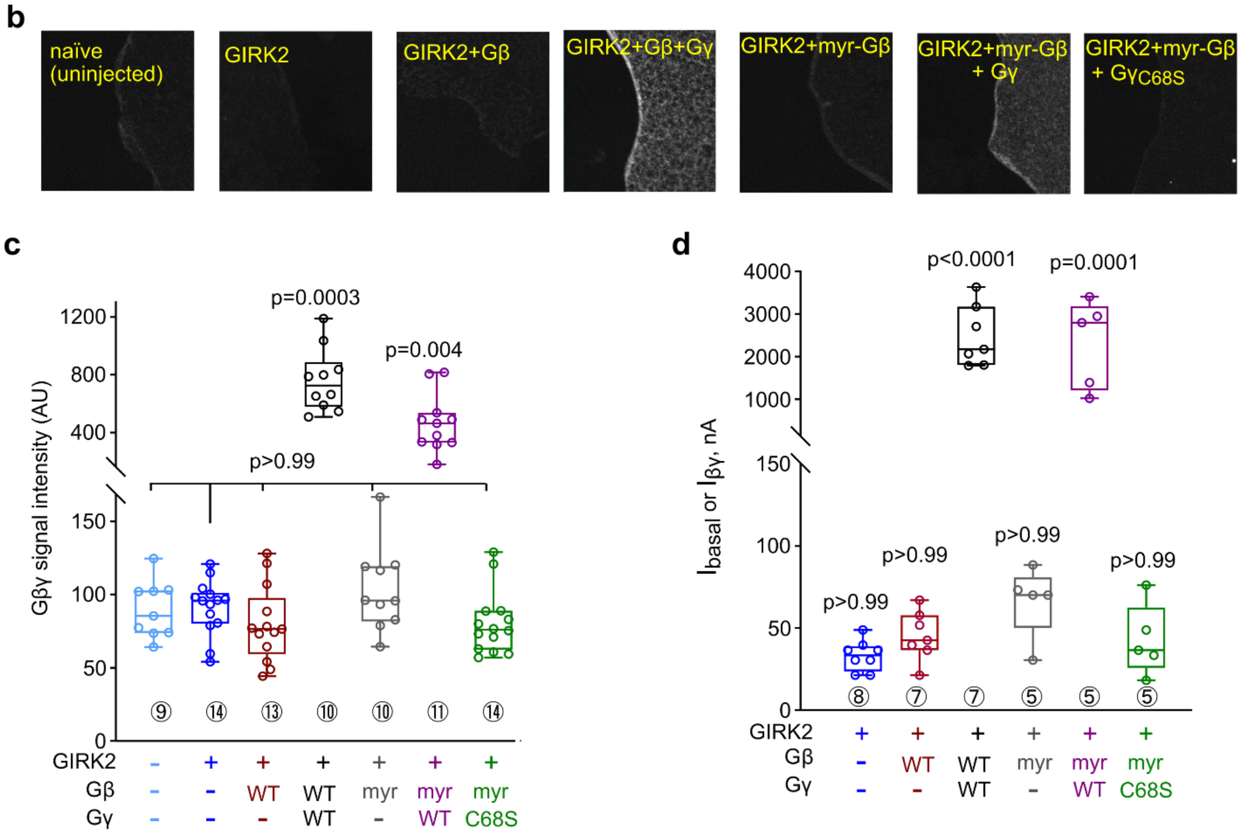


**Supplementary Fig. 1. GIRK2 activation by ACh and Gβγ and the requirement for prenylation of Gγ. a**, basal currents ($I_{basal}$), fold activation by ACh ($R_a$) and fold activation by Gβγ ($R_{\beta\gamma}$). The table shows summary of a series of experiments in which, on the same day, $I_{basal}$ and $I_{evoked}$ were measured in one group of oocytes expressing m2R and GIRK2, and $I_{\beta\gamma}$ in another group of expressing Gβγ, in 24 mM $[K^+]_{out}$ solution. m2R was expressed at 0.5-1 ng RNA, which ensures full maximal attainable $I_{evoked}$[2]. Gβ RNA was 5 ng and Gγ or Gγ-YFP were 1 or 2-2.5 ng, accordingly. Results were grouped according to $I_{\beta\gamma}$ as indicated in the 2$^{nd}$ column. **b**, examples of GMPs from oocytes injected with the indicated RNAs (YFP-GIRK2 was used in this experiment). RNAs injected were (in ng): YFP-GIRK2, 5; Gβ, 5; Gγ or $G\gamma_{C68S}$, 2. 1 ng m2R RNA was present in all groups except native oocytes. Note that, unlike Western blots, the Gβ antibody used here poorly recognized the endogenous oocyte's Gβ in GMP immunostaining[1]. Only $G\gamma_{WT}$ ensures the PM attachment of Gβ or myr-Gβ. Only a weak signal, reflecting the PM-attached endogenous Gβ, is observed in uninjected (native) oocytes or after expression of Gβ or myr-Gβ without Gγ. Note that myr-Gβγ is a functional protein that reaches PM and activates GIRK2 when expressed with $G\gamma_{WT}$. $G\gamma_{S68S}$ is unable to assist in enriching Gβγ in the PM. **c**, summary of Gβ measurements from **b**. Statistics: Kruskal-Wallis test on ranks followed by Dunn's multiple comparison vs. control group (native oocytes). **d**, Summary of GIRK currents measured in 24 mM $[K]_{out}$ in the same experiment as a, b. Statistics: as in c. In **c** and **d**, number of cells tested is shown near data boxes (encircled numbers).

*Fig. S2*

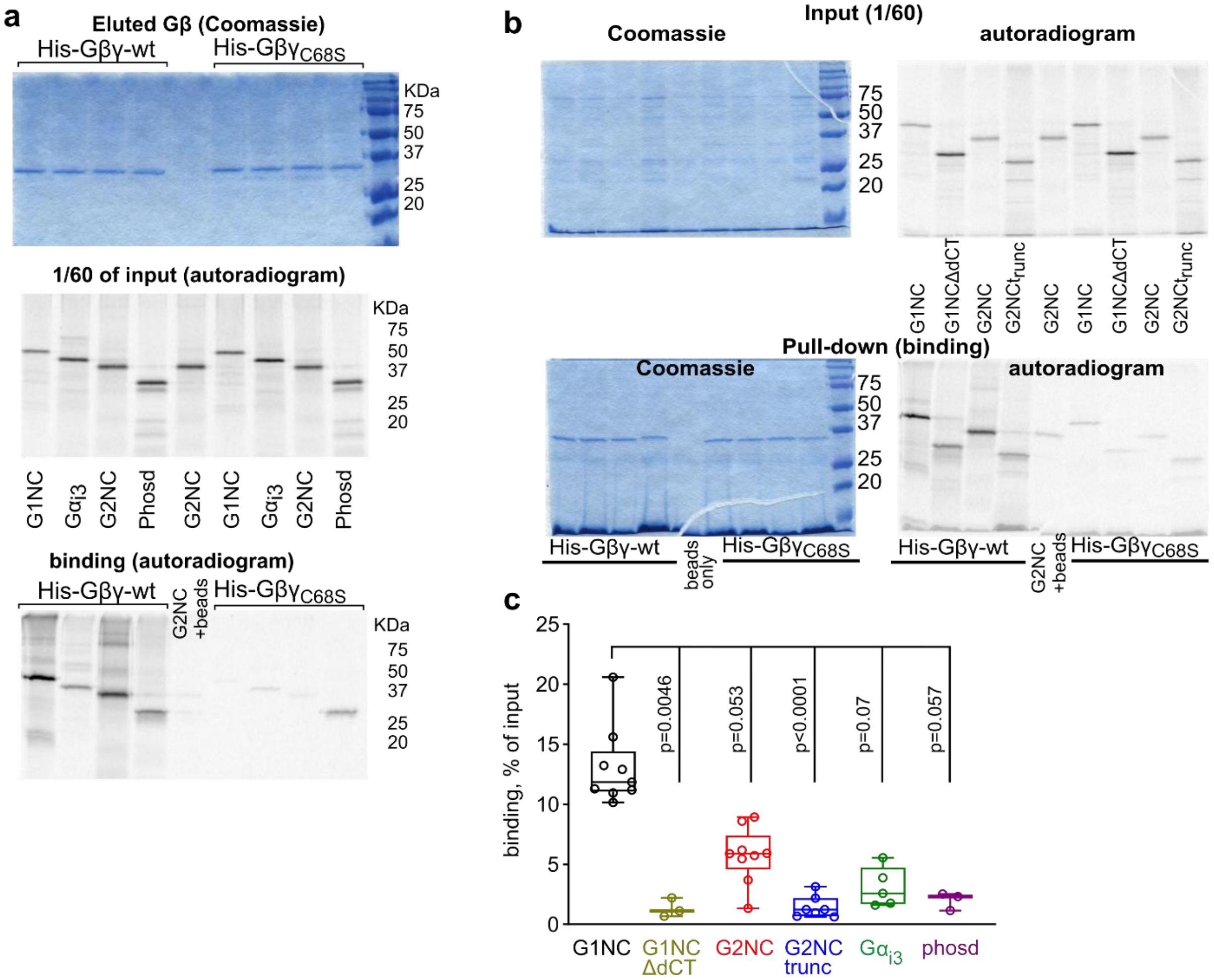


**Supplementary Fig. 2. Differences in binding of Gα$_{i3}$-GDP, phosducin and cytosolic segments of GIRK1 and GIRK2 to WT Gβγ and non-prenylated Gβγ. a,** the full gel of the experiment shown in Fig. 1c. **b**, a representative experiment comparing binding of *ivt* full-length G1NC and G2NC and their truncated versions, G2NC$_{trunc}$ and G1NC$_{ΔdCT}$, to WT His-Gβγ and non-prenylated His-Gβγ$_{C68S}$. Upper and lower images represent two separate gels. Left images show Coomassie staining of proteins in reaction mix (1/60 of total; "input (1/60)", top) and eluted proteins (binding; bottom). Right images show autoradiograms of the same gels. **c**, comparison of binding of the various interactors to Gβγ$_{WT}$. Shown are the same data as in Fig. 1d but without the binding to Gβγ$_{C68S}$. Statistics: Kruskal-Wallis test followed by Dunn's multiple comparison test vs. control (G1NC).

*Fig. S3*

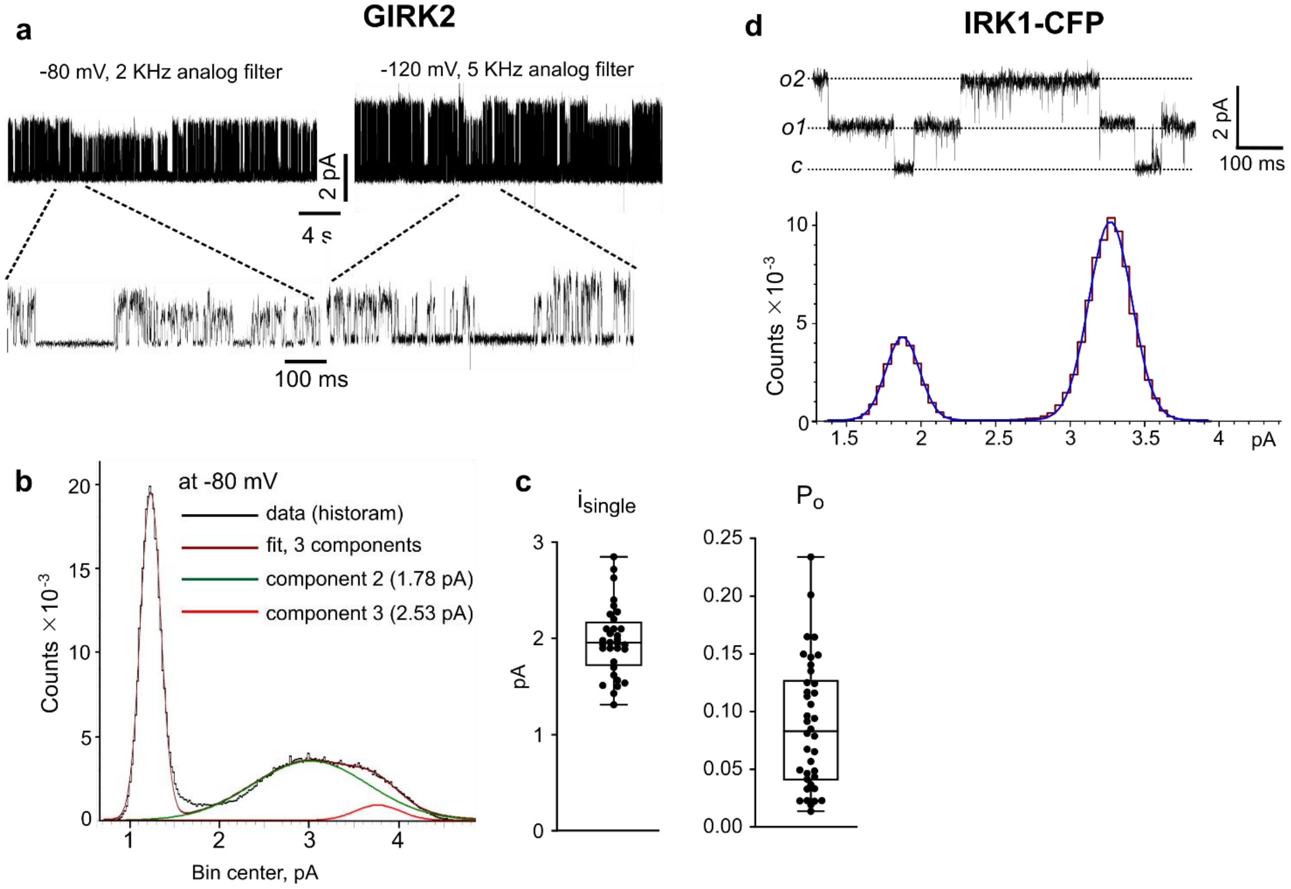


**Supplementary Fig. 3. Characterization of single-channel parameters of GIRK2 and IRK1-CFP. a-c**, analysis of a cell-attached record from an oocyte injected with 25 ng of the anti-GIRK5 oligonucleotide and the following RNAs (in ng/oocyte): GIRK2, 0.017; m2R, 2; Gβ, 5; Gγ, 1. The patch contained one active channel, as assessed from lack of overlaps during the ~10-minute record. **a,** representative segments of the record at -80 mV and -120 mV acquired at 20 KHz with either 2 KHz or 5 KHz analog filter, as indicated. Inward $K^+$ currents are shown as upward deflections. **b,** all-points amplitude histogram of the left record from **a** (at -80 mV). The left peak corresponds to the background noise. The parameters of the 3 components of the Gaussian fit were: $\mu_1$=1.23 pA, $\sigma_1$=0.11 pA; $\mu_2$=3.01 pA, $\sigma_2$=0.64 pA; $\mu_3$=3.76 pA, $\sigma_3$=0.25 pA. The two components of the fit corresponding to the two subconductance levels are shown with green and red lines. Two conductance levels (substates) have been observed in a minority of GIRK2 records. The proportion of the two levels varied among patches, but usually either the larger or the smaller one was predominant (see, for example, Fig. 3c,d). In most cells the channel current in all-point amplitude histograms was well fitted with one Gaussian component. **c,** single channel parameters of GIRK2. $P_o$ was calculated from 2 to 4 min segments of idealized traces from patches containing 1 to 3 channels. The weighted averaged $i_{single}$ was calculated from all-points histograms. Amplitude analysis was limited to patches with $P_o$>0.05 to avoid filtering artifacts with very short openings. **d**, section of a record from an oocyte injected with 5 pg IRK1-CFP RNA. Holding potential was -80 mV. Two channels were present; *c* denotes closed channel current level, *o1* – one open channel, *o2* – two open channels' current. *Bottom*, all-points histogram of a section of the record from the same patch, fitted with a two-component Gaussian for determining the single channel amplitude, $i_{single}$. In this patch, $P_o$ was 0.81 and $i_{single}$ was 1.39 pA. A summary of single channel parameters for all channels used here is presented in Supplementary Table 3.

*Fig. S4*

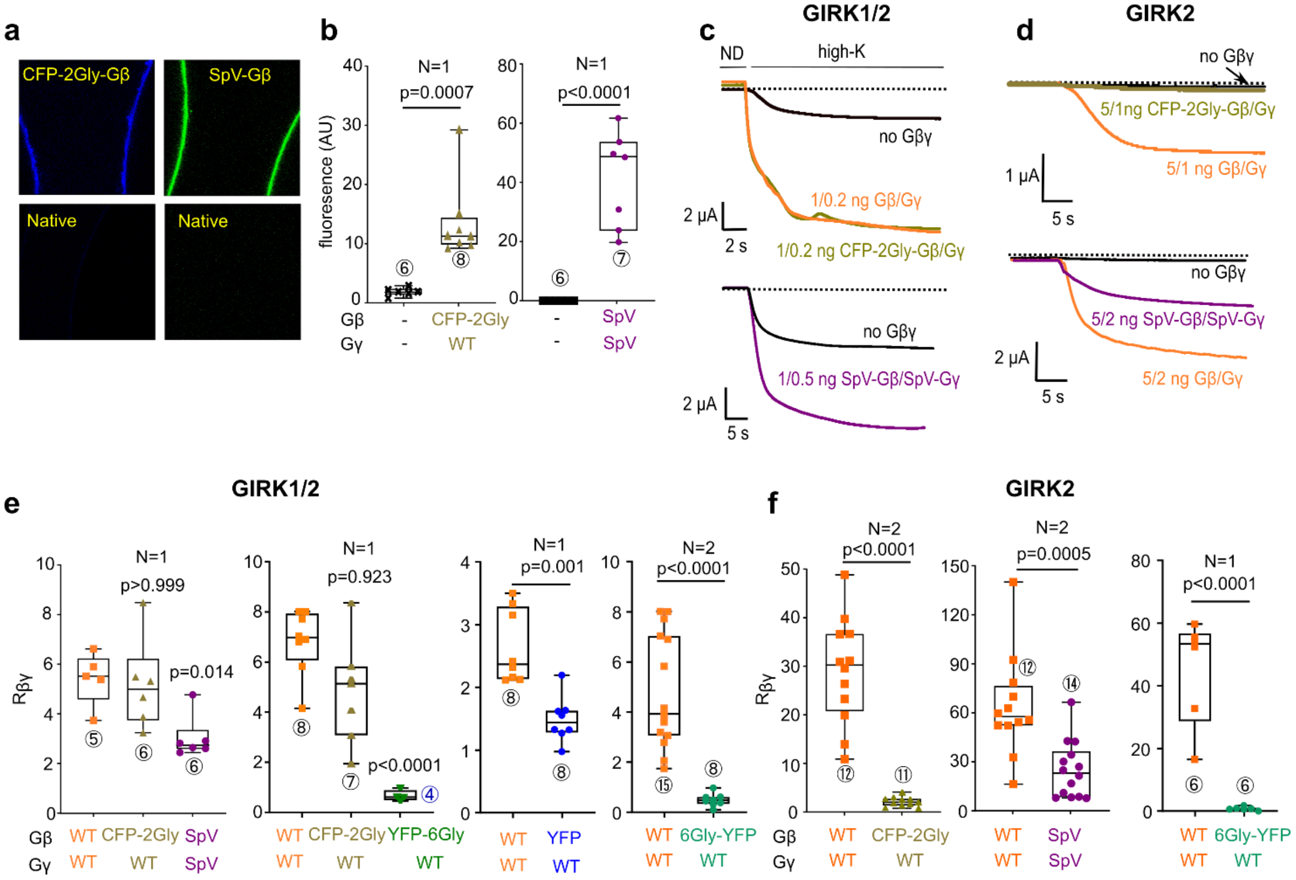


**Supplementary Fig. 4. xFP-fused Gβ constructs perform poorly in activating GIRKs. a,** examples of confocal images of oocytes injected with GIRK2 (2 ng RNA) and RNAs of CFP-2Gly-Gβ (1 ng) and Gγ (0.4 ng), or SpV-Gβ (1 ng) and SpV-Gγ (0.5 ng). Both SpV-Gβγ and CFP-2Gly-Gβ were expressed in the plasma membrane. The intensity of images from the CFP-2Gly-Gβ experiment (both native and Gβ-expressing oocytes) was enhanced 2-fold using Corel PhotoPaint, for better visibility. **b,** summary of expression levels of CFP-2Gly-Gβ and SpV-Gβγ in whole oocytes, compared to the background fluorescence of native oocytes. AU, arbitrary units. Unpaired t-test was used for CFP-2Gly-Gβ and Mann-Whitney test was used for SpV-Gβγ. **c, d**, representative whole-cell currents in oocytes expressing GIRK1/2 (**c**) and GIRK2 (**d**). The amounts of injected RNAs were: GIRK1, 0.05 ng; GIRK2, 0.05 ng as heterotetramer and 2 ng as homotetramer. The amounts of Gβ or xFP-Gβ were: 1 ng when coexpressed with GIRK1/2 and 5 ng with GIRK2. The record started in a low-$K^+$ external solution (ND96; ND) which was then switched to high-K solution (24 mM $[K^+]_{out}$). **e, f**, summary of fold-Gβγ activation ($R_{\beta\gamma}$) by Gβγ vs. the various xFP-labeled Gβ with WT Gγ (except for SpV Gβγ). Gβ RNA doses were as in **c**, except Gβ-6Gly-YFP, which was 5 ng RNA/oocyte in all cases. Of all constructs, only CFP-2Gly-Gβ activated GIRK1/2 similarly to Gβγ (**e**), but it did not activate GIRK2 (**f**). SpV-Gβγ activated both GIRK2 and GIRK1/2 but significantly less than $G\beta\gamma_{WT}$. With Gβ-6Gly-YFP (YFP fused to Gβ's C terminus via a 6-glycine linker), no activation at all was seen with GIRK2 (**f**, right panel), and GIRK1/2 was even reduced relative to its $I_{basal}$ (**e**, right panel; $R_{\beta\gamma}$=0.49±0.09). Statistical analysis included the Shapiro-Wilk normality test, and the following analysis was done using the appropriate tests as detailed in Methods. Number of cells (encircled) is shown near the columns, and number of experiments is shown as N.

*Fig. S5*

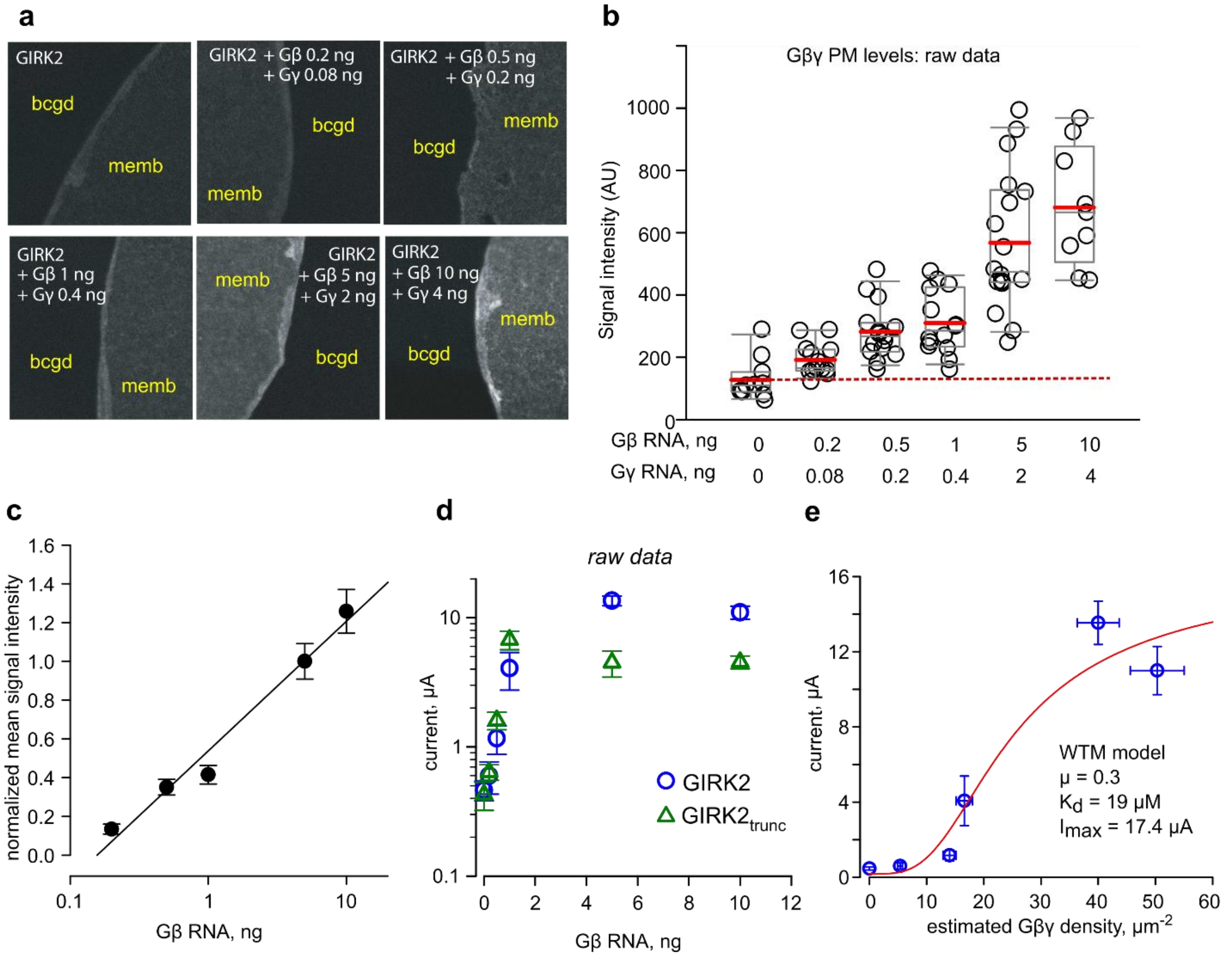


**Supplementary Fig. 5. Dose-dependent Gβγ activation of GIRK2 and GIRK2$_{trunc}$.** Whole-cell currents of the two constructs were compared in one experiment. **a,** representative images of giant plasma membrane patches (GMP) stained with Gβ antibody (Santa Cruz, SC-378). GIRK2 channel (2 ng RNA/oocyte) was present in all groups, and Gβ and Gγ RNAs were coinjected in the indicated amounts. bckg, background; memb, membrane. **b**, summary of measurements of Gβγ in GMPs. AU, arbitrary units. Nine to 18 GMPs were measured with each Gβγ dose. Boxes show 25-75 percentiles and whiskers 5-95 percentiles. Red lines within the boxes show the mean and the grey lines the median values. **c**, relation between the amount of injected Gβ RNA and Gβ protein measurement in the GMPs. Circles show mean±SEM from the same measurements shown in b. In this experiment, the surface levels of expressed Gβγ were empirically found to be linearly related to log[Gβ RNA dose]. Data are shown as mean±SEM, with linear regression line. Net values of expressed Gβγ, after subtraction of the average background signal measured in native oocytes, were normalized to the signal seen with 5 ng Gβ RNA. **d**, whole-cell GIRK2 and GIRK2$_{trunc}$ (2 ng RNA/oocyte) currents measured in 96 mM $K^+$ solution. GIRK2$_{trunc}$ lacks the first 51 amino acid residues (a.a.) of the cytosolic N-terminus and the last 24 a.a. of the CT. **e,** dose-dependent activation of GIRK2 by Gβγ. The density of expressed Gβγ in the PM obtained with 5 ng Gβ RNA was assumed to be 40 molecules/μm$^2$ (as in Fig. 3 and close to the average 35 molecules/μm$^2$ from qWB data, Supplementary Table 4), and Gβγ PM densities for other RNA doses were calculated based on the regression line from Fig. 2c. The circles show mean estimated Gβγ density (±SEM) on the X axis (n=8-18) and mean±SEM current on the Y-axis (n=7-13). The parameters of WTM model fit with fixed μ=0.3 (red line) are shown in the inset.

*Fig. S6*

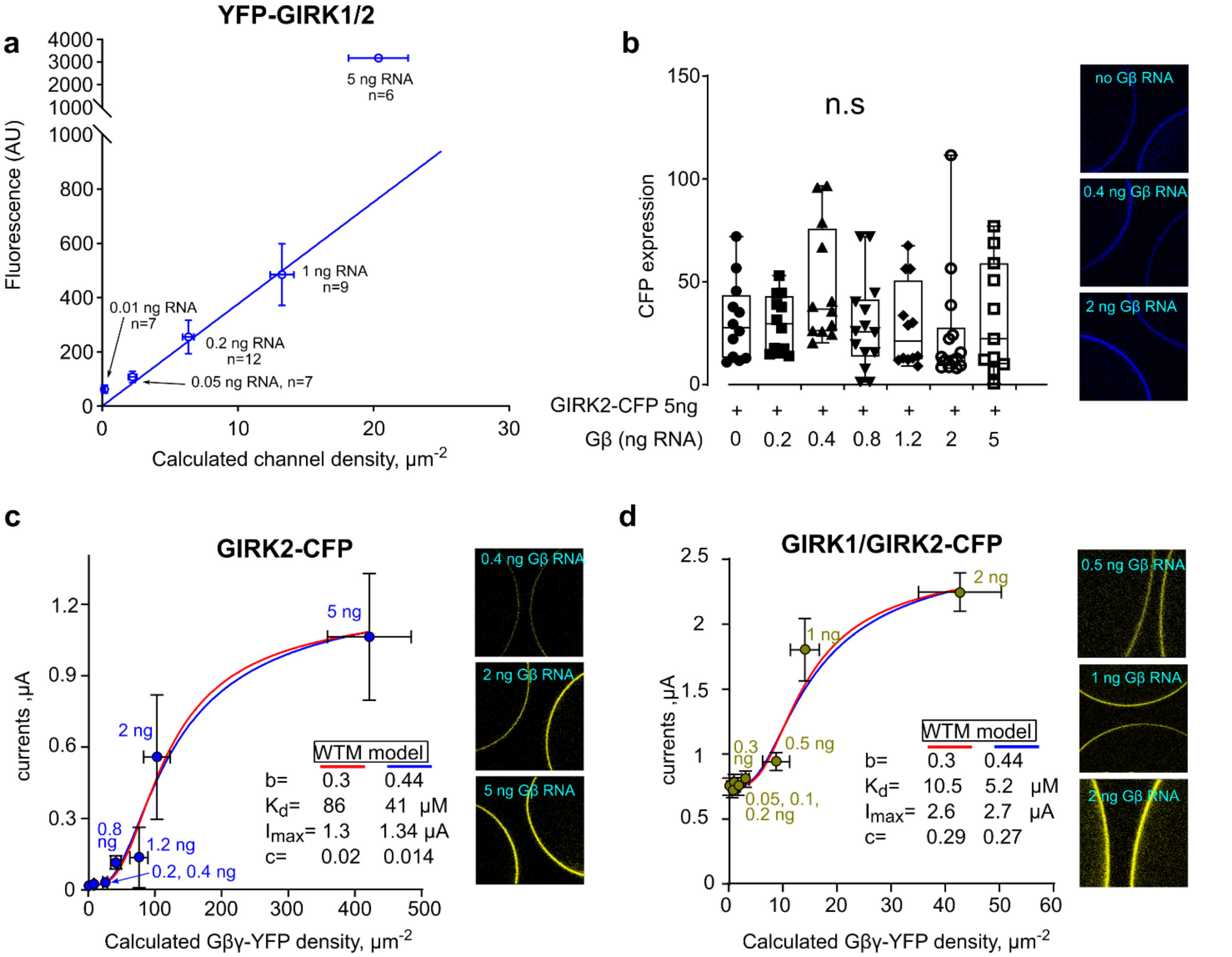


**Supplementary Fig. 6. Testing linear range of CE calibration (a) and dose-dependent activation of GIRK2-CFP and GIRK1/2-CFP by coexpressed $G\beta\cdot_{YFP}G\gamma$ (b-d). a**, testing linearity of surface fluorescence of YFP vs. surface density of YFP-GIRK1/2 in a wide range of channel subunits' RNA (0.01-5 ng/oocyte of each subunit, shown near each point) coexpressed with Gβγ (5:1 ng RNA). YFP fluorescence was measured in 10 oocytes in each group; for current measurements, n is shown near the points. Data are shown as mean±SEM. The linear regression line was drawn via zero and all data points except 5 ng RNA (we never used more than 1 ng in GIRK1/2 experiments). Deviations from linearity were observed in all 3 experiments where 5 ng GIRK1/2 RNAs were tested. **b-d**, data are from one experiment. Calibration was done with YFP-GIRK1/2 coexpressed with Gβγ (5:1 ng RNA; not shown). **b**, GIRK2-CFP (5 ng RNA/oocyte) was expressed with $G\beta\cdot_{YFP}G\gamma$ (2:1 ratio Gβ to YFP-Gγ RNA), and its expression was measured from confocal images of intact oocytes (right panel). Coexpression of $G\beta\cdot_{YFP}G\gamma$ did not significantly affect the expression of GIRK2-CFP in the PM. Boxes show 25-75 percentiles, whiskers the full data range, lines within the boxes show the median. Statistics: Kruskal-Wallis ANOVA followed by Dunn's multiple comparison test. **c**, dose-dependent activation of GIRK2-CFP by $G\beta\cdot_{YFP}G\gamma$. The expression of Gγ-YFP was measured in 11-14 intact oocytes for each group (exemplary images are in the right panel). GIRK currents were measured in 8-10 oocytes. WTM fits with μ=0.3 (red line) and μ=0.44 (blue line) are shown. **d,** dose-dependent activation of GIRK1/GIRK2-CFP (50 pg RNA/oocyte of each subunit) by $G\beta\cdot_{YFP}G\gamma$. $G\beta\cdot_{YFP}G\gamma$ expression in the PM was measured from 9-12 intact oocytes for each $G\beta\cdot_{YFP}G\gamma$ dose. Representative images are shown on the right. GIRK currents were measured in 5-18 oocytes. WTM fits with μ=0.3 (red line) and μ=0.44 (blue line) are shown. This result was not included in the summary of Fig. 4h and Supplementary Table 6, because GIRK2-CFP showed lower sensitivity to Gβγ than WT GIRK2, as judged both by the small magnitude of $I_{\beta\gamma}$ and the high $K_d$.

*Fig. S7*

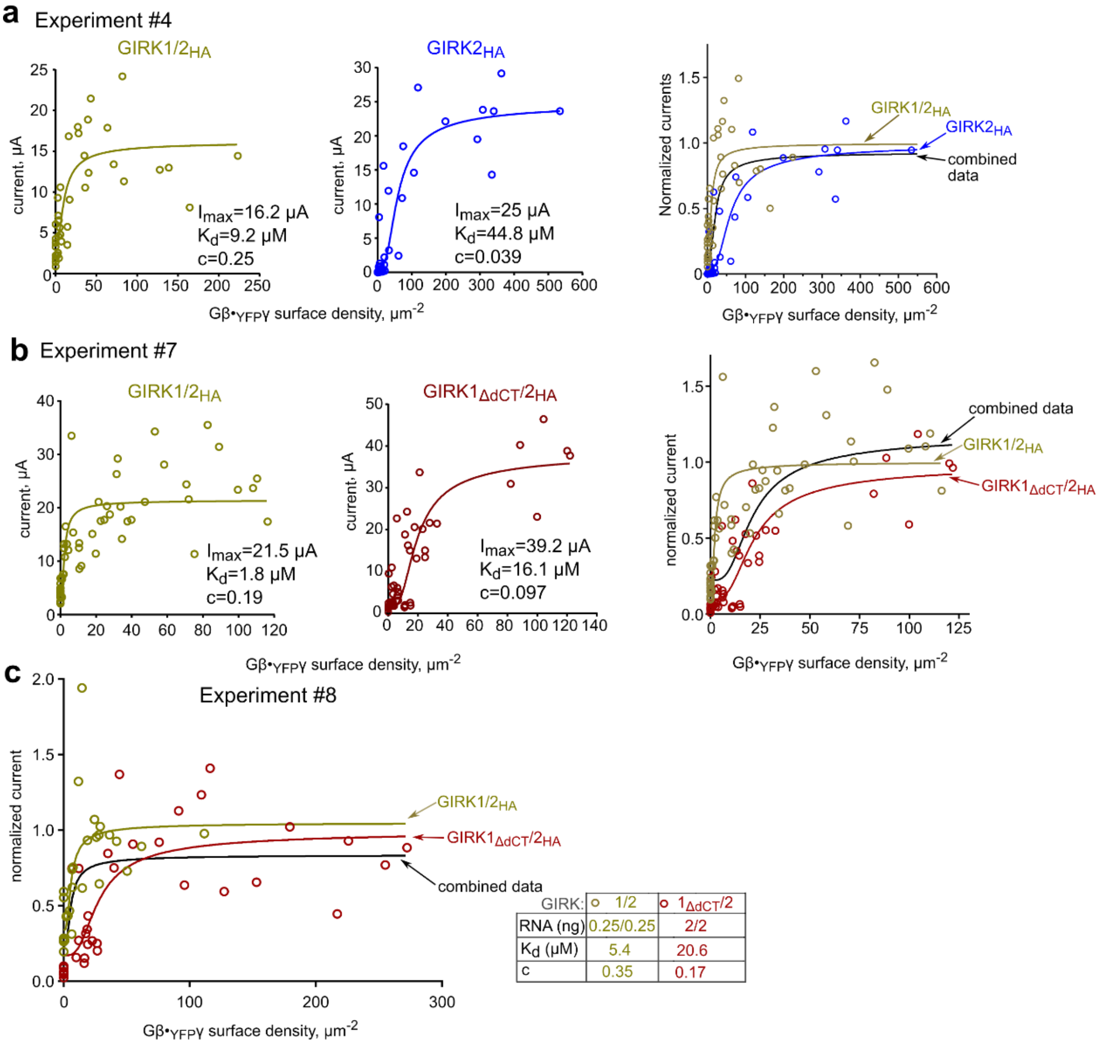


**Supplementary Fig. 7. GIRK2 and GIRK1$_{\Delta dCT}$ show lower apparent affinity to Gβγ than GIRK1/2: raw data in individual oocytes and WTM fits.** Data were fitted with the WTM model with μ=0.3; fit parameters are in the panels. **a,** activation of GIRK1/2$_{HA}$ (left; 0.25 ng RNA of each subunit) and GIRK2$_{HA}$ (middle; 5 ng RNA/oocyte) by Gβ·$_{YFP}$Gγ (0.2-10 ng Gβ, 5:1 Gβ:Gγ RNA). Right panel: WTM fits of currents normalized to maximal $I_{\beta\gamma}$ ($I_{max}$, Supplementary Table 6) and fitted as in Fig. 4 but, in this case, we also included a comparison with the combined data (experimental points for both channel compositions) corresponding to null hypothesis (no difference between the dose-response curves). The analysis was done according to GraphPad guide: https://www.graphpad.com/guides/prism/latest/curve-fitting/reg_comparing_fits_with_anova.htm. Briefly, the data for two channel compositions were fitted separately and together (combined data) to Eq. 5. The $K_d$, n and the SEM parameter of goodness of fit from the three fits were compared using one-way ANOVA and the p value was extracted from F-test and AICc test reports. The differences between the three sets were significant, F(2,162)=146460, p<0.0001 by F test; P<0.01 by AIC test). **b**, GIRK1/2$_{HA}$ (left;, 0.25 ng RNA/oocyte) and GIRK1$_{\Delta dCT}$/2-HA (middle; 2 ng RNA/oocyte) were activated by coexpression of Gβ (0.1-5 ng RNA/oocyte) and YFP-Gγ in 2:1 RNA ratio. Right panel: WTM fits of normalized currents (as in **a**) fitted to the WTM model (μ = 0.3) for GIRK2$_{HA}$, GIRK1/2$_{HA}$, and the combined data. The differences between fits were significant: F(2,206)=28358, P<0.0001; P<0.01 by AIC test. **c**, dose-dependent activation of GIRK1/2$_{HA}$ and GIRK2$_{HA}$ by Gβ·$_{YFP}$Gγ (experiment #8). Analysis and presentation of data are as the right panels in a and b. The differences between the fits were significant, F(2,116)=24243, P<0.0001. In a pairwise comparison between GIRK1/2$_{HA}$ and GIRK2$_{HA}$, the difference in $K_d$ was significant: F(1, 58)=12.15, p=0.0009.

*Fig. S8*

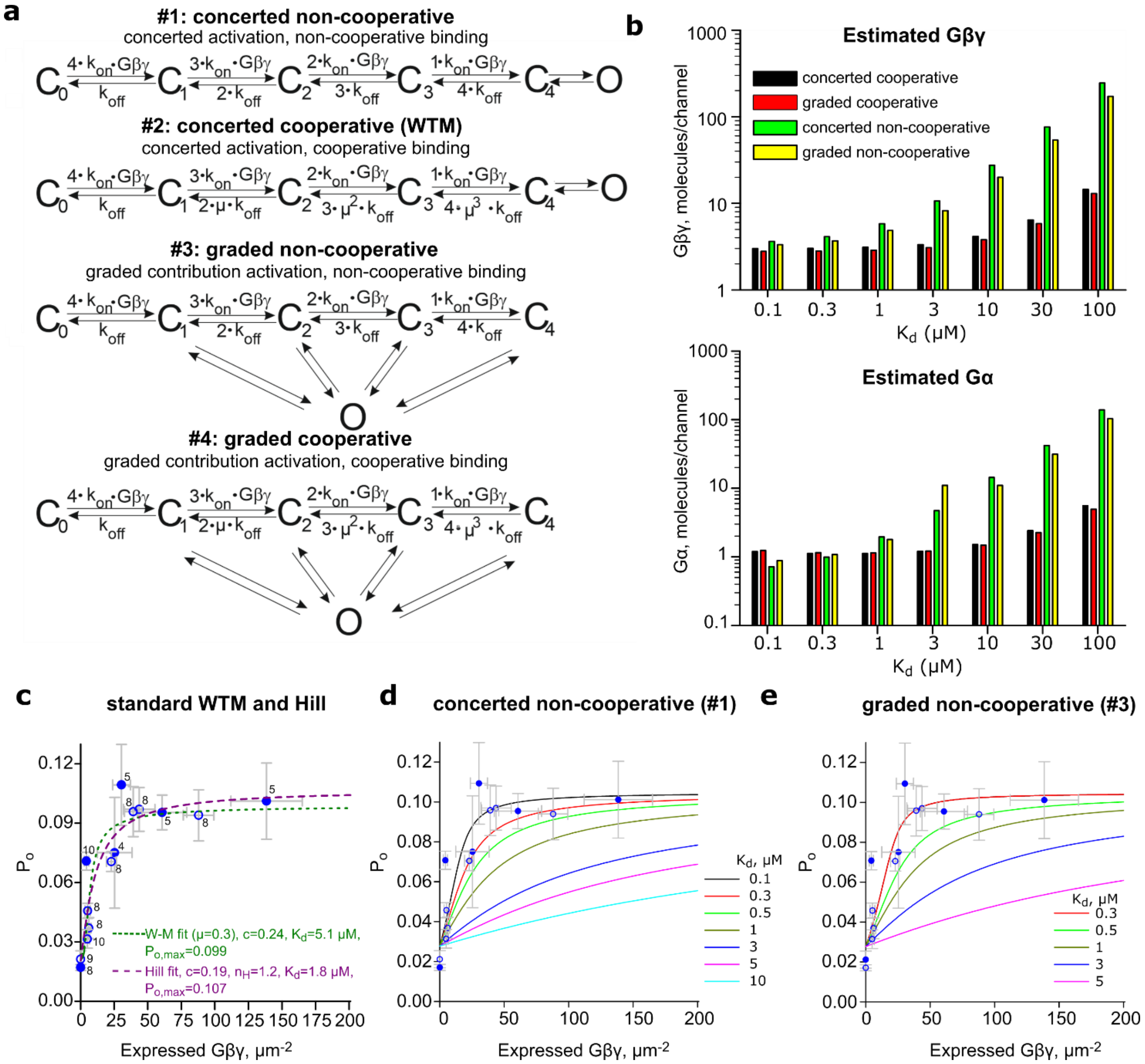


**Supplementary Fig. 8. Modeling basal and Gβγ-evoked GIRK1/2 activity. a**, the four kinetic models for Gβγ activation of GIRK1/2. Up to four Gβγ bind sequentially to the activation sites. In concerted models, the channel opens only when all four sites are Gβγ-occupied. In graded contribution models, occupation of the first site leads to opening, and binding to each additional site increases the $P_o$ in a more-than-additive manner[1]. In cooperative models, each subsequent Gβγ binds with a higher affinity than the previous one. In the non-cooperative models, the affinity of Gβγ to each binding site is the same. **b**, amounts of available basal endogenous Gα and Gβγ that determine $I_{basal}$ and $I_{evoked}$, per channel, were calculated with each of the kinetic models for a range of $K_d$ (see Supplementary Methods). **c**, data from two experiments with GIRK1/2$_{HA}$ (Supplementary Table 6), #4 (closed symbols) and #7 (open symbols; experiment of Fig. 4d), were pulled and fitted to the standard WTM and Hill models. Each point shows mean±SEM of $P_o$ (Y-axis) vs. Gβ$\cdot_{YFP}$γ surface density (X-axis). Numbers of cells are shown near the data points. The whole-cell current at each Gβγ dose was expressed as fraction of the maximal current obtained in WTM fit (24 and 13.4 μA in experiments #4 and #7) and transformed into $P_o$ assuming $P_{o,max}$=0.105 (Supplementary Table 3). **d**, **e**, simulated Gβγ dose-response curves for the concerted and graded non-cooperative models, with a range of $K_d$ values. The basal $P_o$ of ~0.3 is not a fitted value but emerges from basal levels of Gβγ and Gα (as shown in **b**). The data points are the same as in **c.** The simulations best align with data with $K_d$ between 0.1-0.5 μM.

*Fig. S9*

**a**

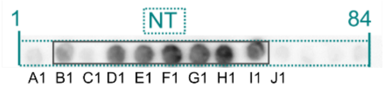


```
B1  RKFGDDYQVVTTSSSGSGLQPQGPG
C1  DYQVVTTSSSGSGLQPQGPGQGPQQ
D1  TTSSSGSGLQPQGPGQGPQQQLVPK
E1  GSGLQPQGPGQGPQQQLVPKKKRQR
F1  PQGPGQGPQQQLVPKKKRQRFVDKN
G1  QGPQQQLVPKKKRQRFVDKNGRCNV
H1  QLVPKKKRQRFVDKNGRCNVQHGNL
I1  KKRQRFVDKNGRCNVQHGNLGSETS
```

**b**

```
hGIRK2   1     makltesm tnvlegdsmd qdvespvaih qpklpkqard dlprhisrdr tkrkiqryvr kdgkcnvhhg nvretyrylt  78
mGIRK2A  1   mtmakltesm tnvlegdsmd qdvespvaih qpklpkqard dlprhisrdr tkrkiqryvr kdgkcnvhhg nvretyrylt  80

hGIRK2   79  difttlvdlk wrfnllifvm vytvtwlffg miwwliayir gdmdhiedps wtpcvtnlng fvsaflfsie tettigygyr  158
mGIRK2A  81  difttlvdlk wrfnllifvm vytvtwlffg miwwliayir gdmdhiedps wtpcvtnlng fvsaflfsie tettigygyr  160

hGIRK2   159 vitdkcpegi illliqsvlg sivnafmvgc mfvkisqpkk raetlvfsth avismrdgkl clmfrvgdlr nshiveasir  238
mGIRK2A  161 vitdkcpegi illliqsvlg sivnafmvgc mfvkisqpkk raetlvfsth avismrdgkl clmfrvgdlr nshiveasir  240

hGIRK2   239 aklikskqts egefiplnqt dinvgyytgd drlflvspli isheinqqsp fweiskaqlp keeleivvil egmveatgmt  318
mGIRK2A  241 aklikskqts egefiplnqt dinvgyytgd drlflvspli isheinqqsp fweiskaqlp keeleivvil egmveatgmt  320

hGIRK2   319 cqarssyits eilwgyrftp vltledgfye vdynsfhety etstpslsak elaelasrae lplswsvssk lnqhaelete  398
mGIRK2A  321 cqarssyits eilwgyrftp vltledgfye vdynsfhety etstpslsak elaelanrae lplswsvssk lnqhaelete  400

hGIRK2   399 eeeknleeqt erngdvanle neskv 423
mGIRK2A  401 eeeknpeelt erng             414
```

**Supplementary Fig. 9. Illustration of analysis of peptide array data and comparison of hGIRK2 with mGIRK2A. a**, the method used to designate the approximate boundaries of Gβγ-binding regions from peptide array scans, exemplified with the GIRK1 NT Gβγ-binding region (highlighted in yellow). The boundaries were arbitrarily defined before the last 10 a.a. of the first 25-mer Gβγ-labeled peptide, and after the first 10 a.a. of the last Gβγ-labeled peptide. In cases when the starting or the last peptide of the array bound Gβγ, 5 N-terminal a.a. were not counted as part of the binding site. **b**, alignment of a.a. sequences of mouse GIRK2A (mGIRK2A) used in most experiments in this report, and hGIRK2 used for the peptide arrays. There are 2 a.a. differences (highlighted in yellow). In addition, hGIRK2 is 2 a.a. shorter in its N terminus and contains 11 a.a. in its dCT not present in mGIRK2A. The last a.a. of mGIRK2A (G414) corresponds to G412 of hGIRK2.

*Fig. S10*

**a**

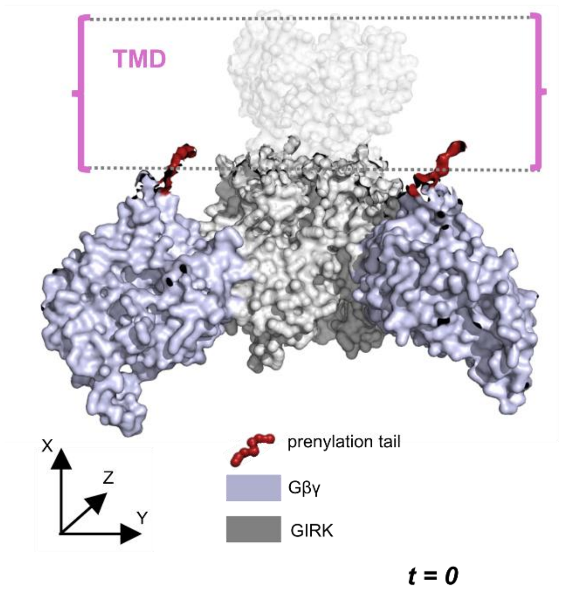


**c**

**Interactions of $G\gamma_{prenyl}$ with Gβ residues (% of simulation run time)**

| | Non-truncated G2NC | | $G2NC_{trunc}$ | |
|---|---|---|---|---|
| Run # | % Gβ (a.a. 314-338) interactions | Top a.a. involved | % Gβ (a.a. 314-338) interactions | Top a.a. involved |
| 1 | 2.12 | W338 (99%), K336 (1%) | 0.7 | W338 (99%) |
| 2 | 2.22 | W338 (95%), F334 (5%) | 0.8 | W338 (95%), F334 (5%) |
| 3 | 0.4 | W338 (100%) | 0.12 | W338 (100%) |
| 4 | 0.4 | W338 (99%) | 0.62 | W338 (99%) |
| 5 | 0.38 | W338 (100%) | 0.13 | W338 (98%) |
| 6 | 0.56 | W338 (99%), F334 (1%) | 0.72 | W338 (99%) |
| 7 | 0.16 | W338 (97%), K336 (3%) | 0.66 | W338 (98%) |
| 8 | 0.18 | W338 (99%) | 0.31 | W338 (99%) |
| 9 | 0.5 | W338 (99%) | 0.82 | W338 (99%) |
| 10 | 0.01 | W338 (71%), K336 (47%) | 0.43 | W338 (48%), K336 (34%), F334 (17%) |
| **Total** | **26.3** | | **53.1** | |

**b**

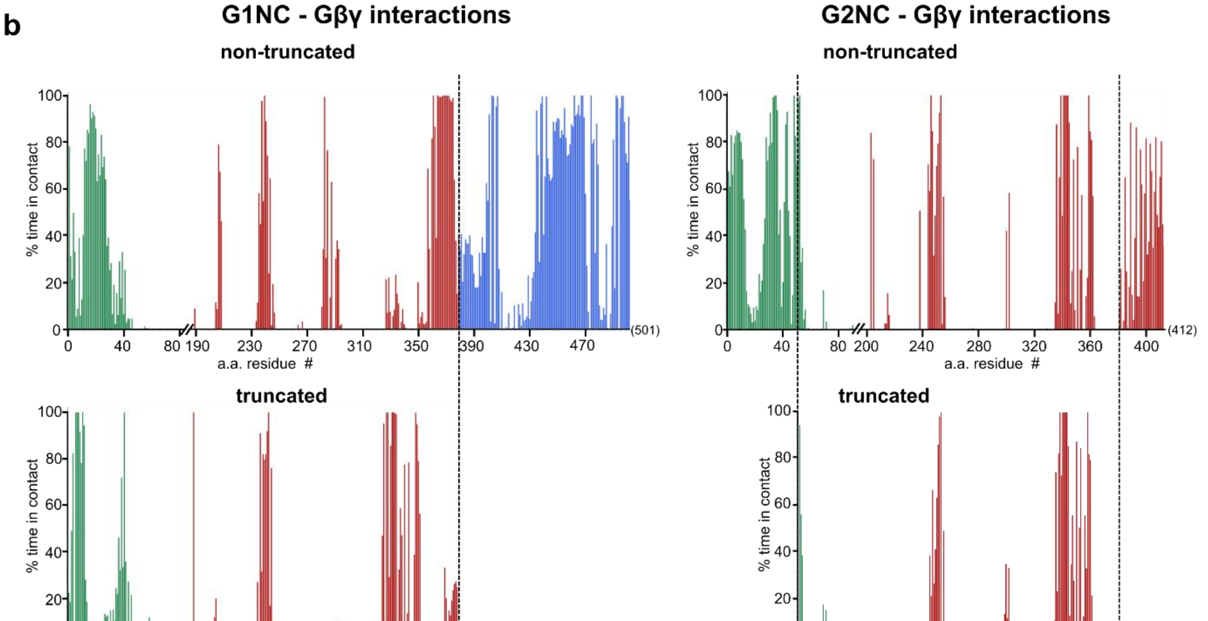


**Supplementary Fig. 10. MD simulations of binding of prenylated Gβγ to full-length and truncated G1NC and G2NC, and of $G\gamma_{prenyl}$ to Gβ. a**, coarse-grained models of G1NC, G2NC, and their truncated versions, which include the bound Gβγ subunits and the prenylation tail ($G\gamma_{prenyl}$). The image shows the experimental system, exemplified for the GIRK2/Gβγ complex at the beginning of the MD simulation (t=0). **b**, histograms showing % of time spent by each a.a. within full-length (non-truncated) and truncated G1NC and G2NC in contact with Gβγ across all runs. **c**, summary of interactions of $G\gamma_{prenyl}$ with the hydrophobic residues of Gβ in the G2NC-Gβγ system.

*Fig. S11*

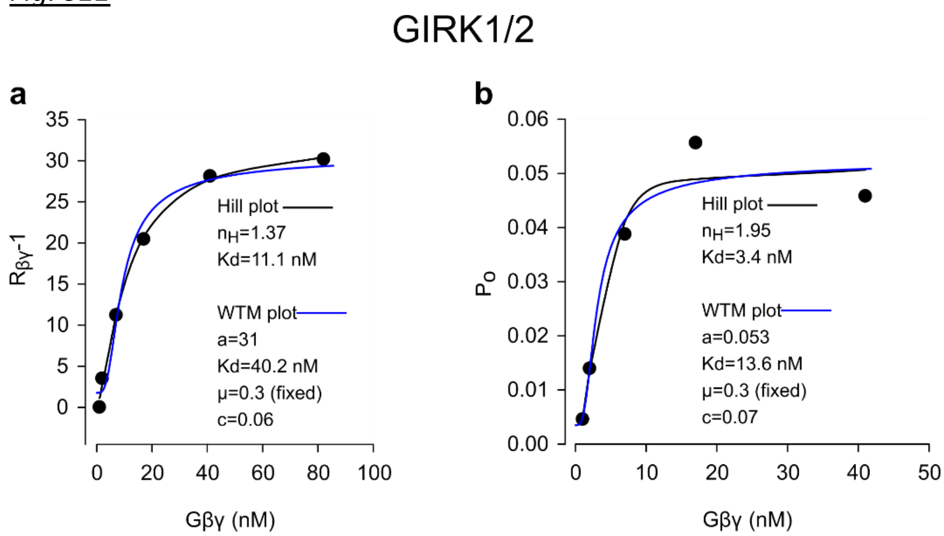


**Supplementary Fig. 11. Dose-dependent activation of GIRK1/2 by Gβγ added to the bathing solution in excised patch experiments.** Original data reported in Peleg et al., 2002[9] were fitted to Hill and WTM models. **a**, results from multichannel patches (>3 channels/ patch; n=4-8 patches for each point). $R_{\beta\gamma}$ is calculated as ($NP_o$ at the peak of activation by Gβγ)/($NP_o$ during the last minute before addition of Gβγ). Channel activity is presented as $R_{\beta\gamma}$-1. **b**, results from patches with 1-3 channels where the exact $P_o$ could be calculated (n=3-8 except the lowest does of Gβγ, where n=1). For additional details, see Fig. 7 in Peleg et al. 2002[9]. Note that basal activity of GIRK1 these patches was 3-7% of maximal $P_o$, which is lower than in our usual whole-cell records. There are two reasons for that: first, activation by Gβγ is stronger for low expression levels of GIRK1/2[1, 9] (to achieve low channel density for single-channel recordings, we injected 5-20 pg RNA vs. 50-250 pg in standard whole-cell experiments, e.g. Fig. 3, Supplementary Fig. 3). Second, Gβγ was applied 3 min after excision when basal activity already decays by ~50% compared to cell-attached mode (Fig. 5). Comparing WTM fits of GIRK1/2 activation by coexpressed Gβγ in whole oocytes ($K_d$=5.5 μM, Fig. 4f) and by purified Gβγ in excised oocyte's patches[9] ($K_d$=13-40 nM) suggests a partition coefficient between 140 and 425.

### Supplementary Tables

#### *Supplementary Table 1. Dissociation constants of Gβγ effectors, including GIRKs.*

| Gβγ-binding protein | $K_d$ | Method and Gβγ used | Ref. |
|---|---|---|---|
| $G\alpha^{GDP}$ (various Gα proteins) | 0.2-27 nM | Fluorescent flow cytometry with bovine brain Gβγ in detergent solutions | 10 |
| PH-PLCγ1<br>Ras-GRF<br>SOS-PH | 318 nM<br>108 nM<br>208 nM | SPR; **prenylated** $G\beta_1\gamma_1$ | 11 |
| KCTD12 (H1 domain) | 185 nM | Isothermal titration calorimetry; **non-prenylated** $G\beta_1\gamma_2$ | 12 |
| D2 (in I-II linker of $Ca_V2.1$ α1 subunit)<br>AID, as above | 24 nM<br><br>63 nM | Pull-down of *ivt* $G\beta_1\gamma_2$ (presumably prenylated) by GST-fused segments of I-II linker on glutathion affinity resin | 13 |
| phosducin | 42 nM | SPR; **prenylated** $G\beta_1\gamma_1$ | 14 |
| GRK2 (βARK1) | 25 nM | Purified **prenylated** $G\beta_1\gamma_2$ added to GRK2 affinity beads (not in lipid phase) | 15 |
| PLCβ2 | 3.2 µM | lipid bilayers, binding curve from FRET, **prenylated** Gβγ concentration in membrane was estimated directly | 16 |
| cardiac $I_{KACh}$<br>GIRK1<br>GIRK4 | 55 nM<br>125 nM<br>50 nM | Immunoprecipitated GIRKs bound to Gβγ on the surface of immunobeads; competition with radiolabeled **prenylated** Gβγ added in detergent solution. | 17 |
| GIRK4, short C-terminal peptide (209-225) | 60 nM | competition of synthetic peptides with atrial GIRK1/4 for binding of **prenylated** $G\beta_1\gamma_2$ in detergent solution | 18 |
| GIRK4 C-terminus | ≤790 nM | SPR; **prenylated** $G\beta_1\gamma_2$ | 19 |
| GIRK1 cytosolic domain, N- and C-terminally truncated | 250 µM | NMR in solution; **non-prenylated** $G\beta_1\gamma_2$ | 20 |
| Purified $GIRK2_{trunc}$ | 1.9 mM (~300 µM in high $Na^+$) | lipid bilayers, dose-response of channel's activity, **non-prenylate**d $G\beta_1\gamma_2$ | 21 |

Abbreviations: AID, α-subunit binding domain; βARK1, β-adrenergic receptor kinase 1; GRK2, G protein-dependent receptor kinase 2; NMR, nuclear magnetic resonance; PH, plekstrin homology domain; PLC, phospholipase C; Ras-GRF, Ras-specific guanine nucleotide exchange factor; SOS, son-of-sevenless protein; SPR, surface plasmon resonance.

***Supplementary Table 2. The values of $K_d$ calculated for sequential cooperative Gβγ binding to GIRK2.***

$K_{d1}$, $K_{d2}$, $K_{d3}$ and $K_{d4}$ are $K_d$ values, in μM, for the binding of the 1$^{st}$, 2$^{d}$, 3$^{d}$ and 4$^{th}$ Gβγ, respectively. The numbers in yellow rectangle correspond to estimates of Wang et al.[21] with 0 to 4 bound $Na^+$ ions per GIRK2 channel. The values of $K_{d1}$ in the following rows have been chosen arbitrarily for illustration purposes and are calculated for the case of constant $Na^+$ concentration with cooperativity factor for Gβγ sequential binding (μ) fixed at 0.3. For calculation of $Na^+$-dependent changes the cross-cooperativity coefficient between $Na^+$ and Gβγ was 0.63[21]. Calculations were done according to the WTM model (Eqn. 5, Methods).

| Na ions/ channel | $K_{d1}$ | $K_{d2}$ | $K_{d3}$ | $K_{d4}$ (μM) |
|---|---|---|---|---|
| 0 | 1900 | 570 | 171 | 51 |
| 1 | 1197 | 359 | 108 | 32 |
| 2 | 754 | 226 | 68 | 20 |
| 3 | 475 | 143 | 43 | 13 |
| 4 | 299 | 90 | 27 | 8 |
| | 30 | 9 | 2.7 | 0.8 |
| | 20 | 6 | 1.8 | 0.54 |
| | 10 | 3 | 0.9 | 0.27 |
| | 5 | 1.5 | 0.45 | 0.135 |
| | 3 | 0.9 | 0.27 | 0.08 |
| | 1 | 0.3 | 0.09 | 0.027 |
| | 0.5 | 0.15 | 0.045 | 0.0135 |
| | 0.2 | 0.06 | 0.018 | 0.0054 |
| | 0.05 | 0.015 | 0.0045 | 0.00135 |

***Supplementary Table 3. Single channel parameters of the various channels***

| | $P_o$ | | | $i_{single}$ (pA) | | |
|---|---|---|---|---|---|---|
| Channel | mean | SEM | n | mean | SEM | n |
| GIRK2 | 0.088 | 0.009 | 38 | 1.98 | 0.065 | 32 |
| IRK1-xFP | 0.81 | 0.028 | 5 | 1.38 | 0.03 | 7 |
| GIRK1/2$_{HA}$ | 0.103 | 0.022 | 4 | 2.64 | 0.081 | 5 |
| YFP-GIRK1/2 + Gβγ (5 ng RNA) | 0.105[1] | | | 2.8[1] | | |

Supplementary Table 3. Single channel parameters of the various channels involved in the calculation of channel surface density, calibration of YFP in the PM from whole-cell currents of various channels, and in simulations. We used known $P_o$ and $i_{single}$ for YFP-GIRK1/2 and determined $P_o$ and $i_{single}$ experimentally under identical conditions for xFP-IRK1, GIRK2 and GIRK1/2$_{HA}$. The values of $P_o$ and $i_{single}$ are from cell-attached patches, at $V_m$=-80 mV, in 146 $[K^+]_{out}$. For GIRK2, average $i_{single}$ was derived from patches with $P_o$>0.05, to avoid filtering effect with short openings.

***Supplementary Table 4. Gβ surface density.***

| *What has been measured* | mean | SEM | N |
|---|---|---|---|
| Endogenous Gβγ in uninjected oocytes– this study | 30 | 11 | 4 |
| Endogenous Gβγ in uninjected oocytes – previous study* | 24 | 4 | 4 |
| Endogenous Gβγ in uninjected oocytes (all data combined) | 28 | 5 | 8 |
| Endogenous Gβγ in oocytes expressing GIRK2 | 35 | 12 | 3 |
| Expressed Gβγ in oocytes injected with RNAs of GIRK2+**Gβγ**** | 35 | 13 | 3 |
| Expressed Gβγ in oocytes injected with RNAs of GIRK2+ $G\beta{\cdot}_{YFP}G\gamma$ ** | 35 | 15 | 3 |
| Expressed Gβγ in oocytes injected with RNAs of GIRK2+ **Gβγ or** $G\beta{\cdot}_{YFP}G\gamma$ combined** | 35 | 9 | 6 |
| Expressed YFP-Gβ in oocytes injected with RNA of YFP-Gβ and WT Gγ* | 28 | 6 | 4 |

Supplementary Table 4. Gβ surface density, in $\mu m^{-2}$, in naïve oocytes, and surface density of expressed Gβ in oocytes injected with RNAs of Gβ or YFP-Gβ (5 ng) and Gγ (1-2ng) or YFP-Gγ (2-2.5 ng). All surface density measurements were done using the quantitative Western blot (qWB) method.

* Yakubovich et al., 2015[1]

** net expression: Gβγ measured in oocytes expressing GIRK2 alone was subtracted from total Gβ reading.

N is the number of experiments.

***Supplementary Table 5. n, N from experiments of Figs. 2 and 4.***

| Fig. 2h,i | Gβγ coexpressed with: | | | |
|---|---|---|---|---|
| | GIRK2: n= | | GIRK1/2: n= | |
| Gβγ RNA (ng) | GMP | Whole oocyte | GMP | Whole oocyte |
| 0.2 | 25 | 36 | 8 | 22 |
| 0.5 | 27 | 36 | 19 | 32 |
| 1 | 42 | 36 | 24 | 33 |
| 2 | 13 | 37 | 5 | 33 |
| 5 | 47 | 32 | 24 | 32 |
| Number of experiments | N=2<br>N=3 | N=3 | N=1<br>N=2 | N=3 |
| Fig. 4a-c | Gβγ coexpressed with: | | | |
| | GIRK2: n= | | GIRK1/2: n= | |
| Gβγ RNA (ng) | surface density | whole-cell current | surface density | whole-cell current |
| 0 | - | 7 | - | 8 |
| 0.2 | 10 | 10 | 12 | 10 |
| 0.5 | 11 | 9 | 6 | 4 |
| 1 | 10 | 9 | 8 | 5 |
| 2 | 9 | 5 | 8 | 5 |
| 5 | 8 | 3 | 8 | 5 |
| 10 | 10 | 5 | - | - |
| Number of experiments | N=1 (all data from one experiment) | | | |

For Fig. 4a-c: In this analysis, we included oocytes measured for both YFP-Gγ fluorescence and $I_{\beta\gamma}$, as well as those assessed only for YFP-Gγ expression. 5 ng Gβ was present in all experiments. In the upper table the numbers highlighted in yellow and in cyan correspond to two different sets of experiments.

*Supplementary Table 6. The complete fitting results to the WTM model*

| GIRK2 | | | | | WTM model, μ=0.3 | | | | | | | | channel density, $\mu m^{-2}$ |
|---|---|---|---|---|---|---|---|---|---|---|---|---|---|
| | | | | | Individual cells | | | Groups | | | Summary | | |
| **Exp. #** | **method** | **calibration** | **GIRK2** | **See Fig.** | **c** | **Kd (μM)** | **Imax (μA)** | **c** | **Kd (μM)** | **Imax (μA)** | **Kd (μM)** | **c** | |
| 1 | c.a. patch | YFP-GIRK1/2 | wt | 3 | 0.037 | 17.1 | Po=0.19 | | | | 17.3 | 0.037 | (0.91) |
| 2 | whole cell | YFP-GIRK1/2 | HA | | | | | 0.044 | 58.5 | 8.6 | 58.4 | 0.044 | 12.1 |
| 3 | whole cell | IRK1-YFP | HA | | 0 | 14.7 | 15.4 | | | | 14.7 | 0 | 21.6 |
| 4 | whole cell | IRK1-YFP | HA | 4a-c | 0.039 | 44.8 | 25 | 0.052 | 45.0 | 24 | 44.8 | 0.039 | 35.1 |
| 5 | whole cell | Gβγ from GMPs | wt | Supp. 5 | | | | 0.010 | 18.4 | 17.4 | 18.4 | 0.01 | 12.7 |
| 6 | whole cell | IRK1-YFP | wt | | 0.050 | 34.3 | 5.63 | 0.000 | 23.8 | 5.1 | 34.3 | 0 | 4.1 |
| | | | | **mean** | **0.032** | **27.7** | **15.3** | **0.027** | **36.4** | **13.8** | **31.3** | **0.022** | **17.1** |
| | | | | **n** | **4** | **4** | **3** | **4** | **4** | **4** | **6** | **6** | **5.0** |
| | | | | **SEM** | **0.009** | **6.2** | **4.56** | **0.011** | **8.1** | **3.7** | **6.6** | **0.008** | **5.3** |
| **GIRK1/2** | | | | | **WTM model, μ=0.3** | | | | | | | | |
| 4 | whole cell | IRK1-YFP | HA | 4a-c | 0.250 | 9.2 | 16.2 | 0.24 | 7.0 | 14.8 | 9.2 | 0.25 | |
| 7 | whole cell | IRK1-YFP | HA | 4d | 0.190 | 1.8 | 21.5 | 0.17 | 5.1 | 24 | 1.81 | 0.19 | |
| 8 | whole cell | IRK1-YFP | wt | | 0.350 | 5.4 | 12.8 | 0.31 | 5.0 | 13.4 | 5.4 | 0.35 | |
| | | | | **mean** | **0.263** | **5.5** | **16.8** | **0.24** | **5.7** | **17.4** | **5.5** | **0.263** | |
| | | | | **n** | **3** | **3** | **3** | **3** | **3** | **3** | **3** | **3** | |
| | | | | **SEM** | **0.038** | **1.7** | **2.06** | **0.033** | **0.5** | **2.7** | **1.7** | **0.038** | |
| **$GIRK1_{\Delta dCT}$/GIRK2** | | | | | **WTM model, μ=0.3** | | | | | | | | |
| 7 | whole cell | IRK1-YFP | HA | 4d | 0.097 | 16.1 | 39.2 | 0.06 | 14.1 | 38.9 | 16.125 | 0.097 | |
| 8 | whole cell | IRK1-YFP | wt | | 0.17 | 20.6 | 26.1 | 0.14 | 21.2 | 26.3 | 20.584 | 0.17 | |
| | | | | **mean** | **0.134** | **18.4** | **32.6** | **0.1** | **17.7** | **32.6** | **18.355** | **0.134** | |
| | | | | **n** | **2** | **2** | **2** | **2** | **2** | **2** | **2** | **2** | |
| | | | | **SEM** | **0.026** | **1.6** | **4.64** | **0.028** | **2.5** | **4.5** | **1.6** | **0.026** | |

**Table S6-continued**

| GIRK2 | | WTM model, µ=0.44 | | | | | | | | Free µ, c=0.03 | |
|---|---|---|---|---|---|---|---|---|---|---|---|
| | | Individual cells | | | Groups | | | Summary | | Individuals cells | |
| Exp. # | See Fig. | c | Kd (µM) | Imax (µA) | c | Kd (µM) | Imax (µA) | Kd (µM) | c | µ | Kd (µM) |
| 1 | 3 | 0.024 | 7.4 | Po=0.196 | | | | 7.4 | 0.024 | 0.44 | 7.4 |
| 2 | | | | | 0.039 | 26.4 | 9.2 | 26.4 | 0.039 | | |
| 3 | | 0 | 2.7 | 8.4 | | | | 2.8 | 0 | fit unstable | |
| 4 | 4a-c | 0.03 | 18.5 | 25.6 | 0.029 | 12.6 | 21.4 | 12.6 | 0.029 | 0.62 | 8.2 |
| 5 | Supp. 5 | | | | 0.006 | 9.6 | 20 | 9.6 | 0.006 | | |
| 6 | | 0.07 | 11.2 | 3.5 | 0 | 11.3 | 5.5 | 11.2 | 0.07 | fit unstable | |
| | **mean** | **0.03** | **10** | **12.5** | **0.019** | **15.0** | **14.05** | **11.3** | **0.028** | **0.53** | **7.8** |
| | **n** | **4** | **4** | **3** | **4** | **4** | **4** | **6** | **6** | **2** | **2** |
| | **SEM** | **0.013** | **3.1** | **5.5** | **0.008** | **3.3** | **3.42** | **3.1** | **0.009** | **0.06** | **0.3** |
| **GIRK1/2** | | **WTM model, µ=0.44** | | | | | | | | | |
| 4 | 4a-c | 0.23 | 3.3 | 16.2 | 0.22 | 3.1 | 15.2 | 3.3 | 0.23 | | |
| 7 | 4d | 0.19 | 0.8 | 21.9 | 0.17 | 2.5 | 24.9 | 0.8 | 0.19 | | |
| 8 | | 0.34 | 2.4 | 13.1 | 0.31 | 2.2 | 13.6 | 2.4 | 0.34 | | |
| | **mean** | **0.25** | **2.2** | 17.1 | **0.23** | **2.6** | **17.9** | **2.2** | **0.25** | | |
| | **n** | **3** | **3** | **3** | **3** | **3** | **3** | **3** | **3** | | |
| | **SEM** | **0.04** | **0.6** | **2.1** | **0.03** | **0.2** | **2.9** | **0.6** | **0.04** | | |
| **GIRK1$_{\Delta dCT}$/GIRK2** | | **WTM model, µ=0.44** | | | | | | | | | |
| 7 | 4d | 0.09 | 7.5 | 41.3 | 0.06 | 6.6 | 41.1 | 7.5 | 0.09 | | |
| 8 | | 0.15 | 8.6 | 26.5 | 0.095 | 8.7 | 26.9 | 8.6 | 0.15 | | |
| | **mean** | **0.12** | **8.0** | **33.9** | **0.078** | **7.6** | **34** | **8.0** | **0.12** | | |
| | **n** | **2** | **2** | **2** | **2** | **2** | **2** | **2** | **2** | | |
| | **SEM** | **0.02** | **0.4** | **5.2** | **0.012** | **0.8** | **5** | **0.4** | **0.02** | | |

**Supplementary Table 6.** Experimental details and the WTM model fit parameters for all Gβγ dose-dependence experiments. $I_{max}$ is the fitted maximal current. Cooperativity factor µ was fixed at 0.3 or 0.44. Fits with free µ (with fixed c=0.03) were also performed for GIRK2 data, but stable fits were obtained only in two experiments. GIRK2 channel surface density in experiments #2-6 was calculated from $I_{max}$ (fit with µ=0.3) assuming $P_{o,max}$=0.19. In experiment #1 whole-cell currents in the 5 ng Gβ RNA group were 640±136 nA (n=8), corresponding to 0.91 $\mu m^{-2}$ channel density. All currents in whole-cell mode were recorded in 24 mM $[K]_{out}$ solution. Data are presented for individual cells and groups (where available). In all experiments we measured the surface density of Gβ$_{\cdot YFP}$Gγ except #5, where we monitored relative changes in PM-attached Gβ in GMPs (instead of YFP-Gγ) and assumed Gβ density of 40 $\mu m^{-2}$ with 5 ng Gβ RNA (from Fig. 3). In experiment #2, surface expression of GIRK2$_{HA}$ was measured in groups of cells expressing the different concentration of Gβγ (see Supplementary Methods), and currents were corrected for changes in channel expression.

***Supplementary Table 7. p values for pull-down experiments***

| | p | q | DF |
|---|---|---|---|
| G1NdCT vs. G1NC | 0.0001 | 4.820 | 51 |
| G1NdCT vs. G1NCΔC1 | 0.3787 | 1.913 | 51 |
| G1NdCT vs. G1NCΔC2 | 0.0189 | 3.215 | 51 |
| G1NdCT vs. G1NCΔC3 | 0.0134 | 3.337 | 51 |
| G1NdCT vs. G1CT | <0.0001 | 7.222 | 51 |
| G1NdCT vs. G1(1-40)dCT | <0.0001 | 6.532 | 51 |
| G1NdCT vs. G1(40-84)dCT | 0.9632 | 0.9404 | 51 |
| G1NdCT vs. Sumo dCT | <0.0001 | 6.853 | 51 |
| G1NdCT vs. Sumo NT | <0.0001 | 7.418 | 51 |
| | p | q | DF |
| G1NC vs. G1NCΔC1 | 0.7085 | 1.389 | 51 |
| G1NC vs. G1NCΔC2 | 0.9752 | 0.8313 | 51 |
| G1NC vs. G1NCΔC3 | 0.9998 | 0.4212 | 51 |
| G1NC vs. G1CT | 0.4784 | 1.707 | 51 |
| G1NC vs. G1NdCT | 0.0001 | 4.820 | 51 |
| G1NC vs. G1(1-40)dCT | 0.0932 | 2.562 | 51 |
| G1NC vs. G1(40-84)dCT | 0.0744 | 2.659 | 51 |
| G1NC vs. Sumo dCT | 0.1012 | 2.527 | 51 |
| G1NC vs. Sumo NT | 0.0109 | 3.389 | 51 |

Supplementary Table 7; for Fig. 7: Statistical comparison of Gβγ binding in constructs containing different regions of GIRK1 from pull-down experiments. The table presents p-values (p), t-values (q), and degrees of freedom (DF) for pairwise comparisons using Dunnett's test. G1NC or G1NdCT were used as controls in separate analyses to compare all other samples against them.

***Supplementary Table 8. $K_d$ and Hill coefficients from fits to Hill equation for dose-response relations from excised patch measurements.***

| channel | Gβγ type | preparation | $K_d$ (nM) | $n_H$ | reference |
|---|---|---|---|---|---|
| $I_{KACh}$ (GIRK1/4) | $G\beta_1$ or $G\beta_2$ with $G\gamma_2$, $G\gamma_5$ or $G\gamma_7$ | atrial myocyte | 4-11 | 1.5 | 17 |
| GIRK1/3 | Bovine brain | *Xenopus* oocyte | 11 | 1.5 | 22 |
| GIRK 1/4 | Bovine brain | *Xenopus* oocyte | 10 | 1.5 | |
| neuronal (GIRK 1/2?) | $G\beta_1\gamma_2$ | locus coeruleus neuron | 3.78 | 2.03 | 23 |
| GIRK1/2 | $G\beta_1\gamma_2$ | *Xenopus* oocyte | 11 | 1.37-2.04 | ref. 9 and Fig. S11 |
| GIRK 1/5 (GIRK5 of *Xenopus*) | $G\beta_1\gamma_2$ | *Xenopus* oocyte | 1.8-4.75 | 1.21-1.31 | 24, 25 |
| $I_{KACh}$ (GIRK1/4) | Bovine brain | atrial myocyte | 6 | 3.12 | 26 |

*Supplementary Table 9. DNA constructs used in this work*

| Construct name | Vector | Species | Remarks | Accession number |
|---|---|---|---|---|
| GIRK1 | pGEM-HJ | rat | | NP_113798.1 |
| YFP-GIRK1 | pGEM-HJ | rat | YFP in NT | |
| GIRK1Δ123(dCT) | pGEM-HJ | rat | GIRK1 (1-378) | |
| G1NC WT | pMXT | rat | GIRK1 NT(1-84)-Linker (QSTASQST)-CT(185-501) | |
| G1NCΔdCT | pMXT | rat | GIRK1 NT(1-84)-Linker (QSTASQST)-CT(184-380) | |
| G1NdCT | pMXT | rat | GIRK1 NT(1-84)-Linker (QSTASQST)-dCT(381-501) | |
| G1NCΔC1 | pMXT | rat | GIRK1 NT(1-84)-Linker (QSTASQST)-CT(254-501) | |
| G1NCΔC2 | pMXT | rat | GIRK1 NT(1-84)-Linker (QSTASQST)-CT(184-253, 321-501) | |
| G1NCΔC3 | pMXT | rat | GIRK1 NT(1-84)-Linker (QSTASQST)-CT(184-319, 371-501) | |
| G1N(1-40)dCT | pMXT | rat | GIRK1 NT(1-40)-Linker (QSTASQST)-dCT(381-501) | |
| G1N(40-84)dCT | pMXT | rat | GIRK1 NT(40-84)-Linker (QSTASQST)-dCT(381-501) | |
| Sumo-G1NT | pGEM-HJ | rat | Sumo-GIRK1 NT(1-84) | |
| G1CT | pMXT | rat | GIRK1 CT(184-501) | |
| Sumo-G1dCT | pMXT | rat | Sumo-GIRK1 dCT(381-501) | |
| GIRK2 | pGEM-HJ | mouse | GIRK2A, 414 a.a. | NP_001020755.1 |
| G2NC | PGBXW | mouse | GIRK2 NT(1-95) Linker (QSTASQST) CT(194-414) | |
| GIRK2$_{trunc}$ | pGEM-HJ | mouse | 52-380 a.a | |
| G2NC$_{trunc}$ | pGEM-HJ | mouse | GIRK2 NT(52-95) Linker (QSTASQST) CT(194-380) | |
| GIRK2-HA | pGEM-HJ | mouse | HA tag in P-loop --MDHI-HA-EDPS-- | |
| GIRK2-CFP | pGEM-HJ | mouse | CFP in CT | |
| M2R | pGES | human | | NP_001006631.1 |
| IRK1-YFP, IRK1-CFP | pGSB | mouse | YFP or CFP in CT | NP_032451.1 |
| Gβ1 wt | pGEM-HE | bovine | | NP_786971.2 |
| Myr-Gβ1 | pGEM-HJ | bovine | myristoylated Gβ1 | |
| ***Supplementary Table 9 continued*** | | | | |
| YFP-Gβ1 | pGEM-HJ | Bovine | YFP in NT with a Lys-Ser linker | |
| Split Venus2-Gβ1 | pGEM-HJ | human | Split Venus2 in NT | |
| Gβ1-6Gly-YFP | pGEM-HJ | bovine | YFP in CT, fused to Gβ1 by a linker of GGGGGG (6 glycines) | |

| | | | | |
|---|---|---|---|---|
| CFP-2Gly-Gβ1 | pGEM-HJ | bovine | YFP in NT, fused to Gβ by a linker of GG (2 glycines) | |
| Gγ2 | pGEM-HJ | bovine | | P63212.2 |
| Gγ2 C68S | pGEM-HE | bovine | | |
| Split Venus1-Gγ2 | pGEM-HJ | bovine | Split Venus1 in NT | |
| YFP-Gγ2 | pMXT | bovine | YFP in NT | |
| $G\alpha_{i3}$ | pGEM-HJ | human | | NP_006487.1 |
| Phosducin | pGEM-HE | bovine | myristoylated phosducin | NP_002588.3 |

Mouse GIRK2 (GIRK2A, 414 a.a.), $GIRK2_{HA}$, C-terminally CFP labeled GIRK2 (GIRK2-CFP), bovine $G\beta_1$, bovine $G\gamma_2$, human m2R, rat GIRK1, N-terminally YFP labeled GIRK1 (YFP-GIRK1), N-terminally YFP labeled $G\gamma_2$ (YFP-$G\gamma_2$), N-terminally YFP labeled $G\beta_1$ (YFP-Gβ, with a Lys-Ser linker), G1NC (the cytosolic N-a.a. 1-84 and C-termini a.a 183-501 of GIRK1 connected by an 8-a.a. linker, QSTASQST), G2NC (the full cytosolic N- a.a. 1-95 and C-termini a.a. 194-404 of GIRK2 connected by a 2-a.a. linker, Lys-Leu), full length human $G\alpha_{i3}$ and myristoylated bovine phosducin were described previously[8, 27-29]. TEVC, two-electrode whole-cell voltage clamp.

***Supplementary Table 10. Antibodies used in this study***

| Antibody | SOURCE | IDENTIFIER |
|---|---|---|
| Donkey IgG | Jackson ImmunoResearch Labs | Cat. No. 017-000-003; RRID:AB_2337256; Lot 131795 |
| Rabbit polyclonal anti-Gβ (T-20) | Santa Cruz Biotechnology | Cat. No. sc-378; RRID:AB_631542; currently discontinued |
| GNB1 | GeneTex | GTX114442; RRID:AB_10619473; Lot 43565 |
| Goat Anti-Rabbit IgG H&L–DyLight650 | Abcam | ab96886; RRID:AB_10680254; Lot GR3228258-6 |
| Anti-$His_6$ Peroxidase mouse monoclonal, clone BMG-His-1 | Roche | Cat. No. 11 965 085 001; RRID: AB_514487 |
| Goat Anti-Rabbit IgG, (H+L) HRP conjugated | Jackson ImmunoResearch Labs | Cat. No. 111-035-144, RRID:AB_2307391 |
